# Small molecule-controlled gene expression: Design of drug-like high affinity small molecule modulators of a custom-made riboswitch

**DOI:** 10.1101/2025.01.21.634073

**Authors:** Vera Hedwig, Maike Spöring, Gary Aspnes, Dirk Gottschling, Holger Klein, Matthias Klugmann, Sebastian Kreuz, Benjamin Ries, Gisela Schnapp, Sandra Scharsich, Jörg S. Hartig, Oliver Hucke

**Affiliations:** Department of Chemistry, University of Konstanz, Universitätsstraße 10, 78457 Konstanz, Germany; Konstanz Research School Chemical Biology (KoRS-CB), University of Konstanz, Universitätsstraße 10, 78457 Konstanz, Germany; Boehringer Ingelheim Pharma GmbH & Co. KG, Birkendorferstr.65, 88397 Biberach an der Riß, Germany

## Abstract

Riboswitches are regulatory RNA structures that modulate gene expression in response to a small molecule. Until now, efforts to design ligand analogs were motivated by their potential antibiotic activity. However, riboswitches are ideally suited as tools for gene therapy, enabling precise control of gene ex-pression without the need of potentially immunogenic regulatory proteins. Developing synthetic RNA switches starting from natural riboswitches will require engineering both, the ligand and the RNA sequence in order to achieve full orthogonality i.e., sensitivity to the designed small molecule modulator, but not to the natural ligand. We present the structure-based design of a drug-like small molecule ligand of the thiamine pyrophosphate (TPP) aptamer, BI-5232. BI-5232 is structurally highly diverse from the natural ligand TPP but rivals its binding affinity (K_D_ = 1.0 nM). Importantly, in our design the pyrophosphate of TPP was replaced by an uncharged heterocycle that interacts with the PP helix in an unprecedented way, as revealed by Molecular Dynamics simulations. Subsequently, we altered the aptamer sequence to drastically reduce its affinity to TPP while retaining binding properties for our designed ligand. Based on the developed orthogonal small molecule/RNA aptamer interaction we finally constructed orthogonal ribozyme-based ON- and OFF-switches of gene expression in human cell lines. Such systems are valuable additions to the synthetic toolbox for conditionally controlling gene expression with potential applications in next-generation gene therapies.

## Introduction

Riboswitches are promising tools to regulate gene expression for scientific and medical purposes. Notably, the independency of protein cofactors for their regulatory activity as well as their small size and modular character render them attractive for gene therapies.(1, 2) However, there remains a demand for aptamers binding therapeutically applicable ligands that possess drug-like properties such as high bioavailability, safety and tolerability. Up to date, the ligand-repertoire of switches with functionality in human cells is limited to substances that are present in cells or cause toxic side effects when applied at higher concentrations.(3, 4)

In order to generate aptamers and ligands suited for therapeutic purposes, aptamers can be selected against drug-like compounds, in-vitro.(4) Alternatively, natural aptamers could be re-engineered to display non-natural ligand selectivity and decreased affinity for their natural ligands. For antibiotic purposes, a variety of riboswitches has been targeted with ligand analogs in the past.(5, 6) The TPP riboswitch class was one of the earliest riboswitches identified (7) and is the most widely known, being distributed among all three domains of life.(5, 8, 9) Especially the *E. coli thiM* aptamer has been studied intensively and was subject of several structural and mechanistic studies providing insights into the mechanism of ligand binding.(10-16) It is composed of five stem structures (P1-P5) arranged in two helical sections connected via a three-way junction to a closing stem (P1) that is stabilized upon ligand binding (Figure 1).(10, 13) When TPP is present, the 4-amino-5-hydroxymethyl-2-pyrimidine (HMP) moiety is recognized by the J3-2 junction of the ‘pyrimidine-sensor helix’ (Py-helix) while the pyrophosphate is bound as a complex with two Mg^2+^-ions by the ‘pyrophosphate-sensor helix’ (PP-helix) (Figure 1).(10, 13) Herein, TPP acts as a bridge that connects the two helical regions and locks them in a ‘Y-shape’, which is further stabilized by tertiary interactions.(10, 13)

**Figure 1:**
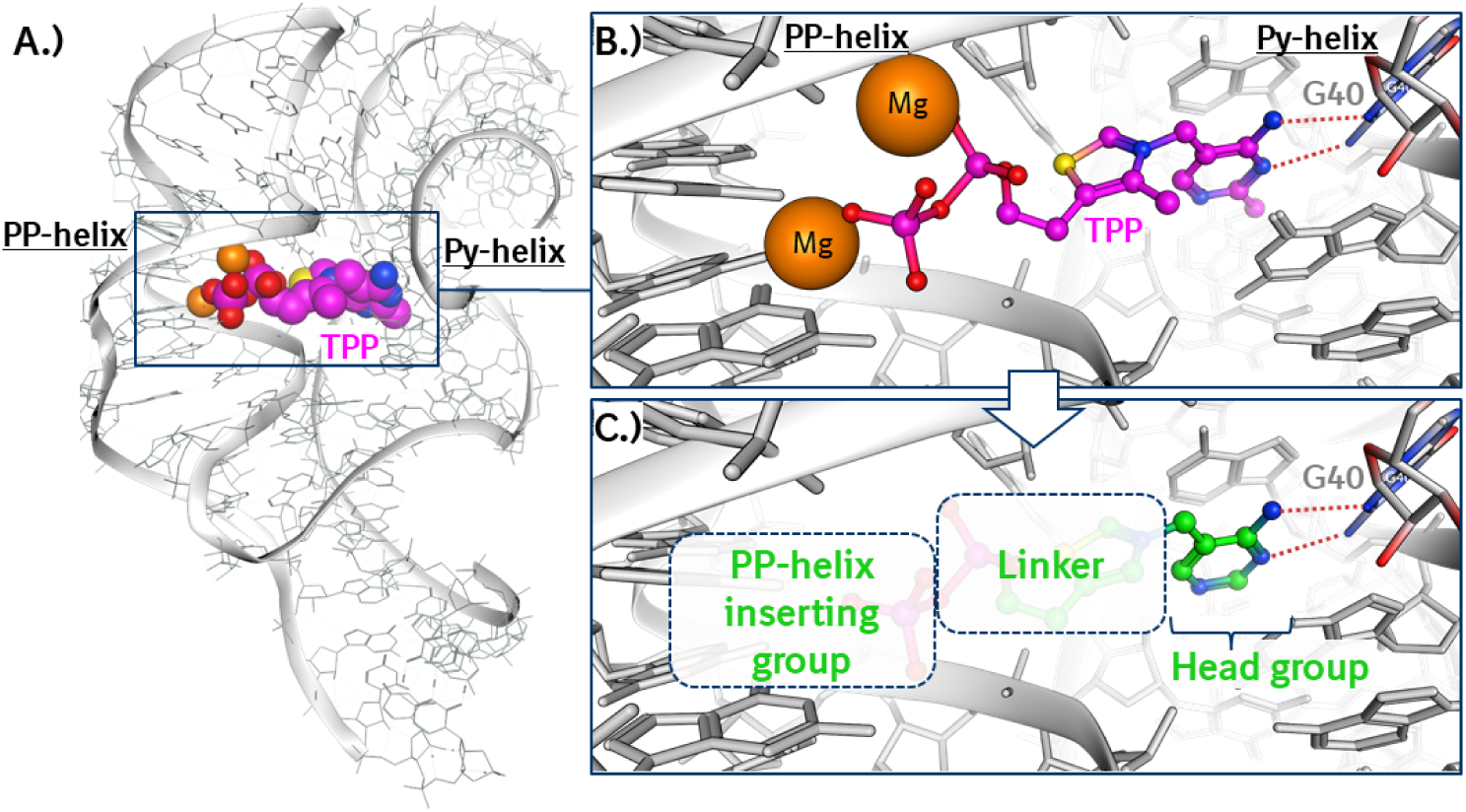
**A.** Binding mode of TPP in the thiM aptamer as observed in an X-ray structure (PDB code 2GDI) (10), used as starting point for design. **B**. Close-up of the TPP binding mode. Mg ions mediating polar interactions with the PP-helix and hydrogen bonds (dashed red lines) to G40 in the Py-helix are indicated. **C**. The design strategy was to identify a linker structure that allows the insertion of a drug-like replacement of the pyrophosphate group into the PP helix. Head group key H-bonds with G40 (red dashed lines) were meant to be conserved.

TPP analogs are able to bind to the *thiM* aptamer as well, including the monophosphorylated and unphosphorylated variants thiamine monophosphate and thiamine as well as pyrithiamine, benfotiamine, and amprolium.(13, 17) Several studies have investigated the determinants of ligand binding to the TPP aptamer and assessed the antibiotic potential of TPP analogs targeting the TPP riboswitch. Chen *et al.* have dissected the TPP molecule into three parts (pyrimidine, thiazole and pyrophosphate moieties), sequentially exchanged them by mostly closely related analogous structures and evaluated their potency to displace thiamine from the aptamer.(17) The same group also performed a fragment-based screen to identify alternative molecules that are bound by the TPP aptamer.(18, 19) Lünse *et al.* analyzed thiamine analogues that bear a triazole instead of a thiazole moiety as alternative thiM riboswitch inducers.(20) These triazole-based compounds were somewhat effective thiamine analogs, in spite of the loss in binding affinity that is known to be a consequence of replacing the thiazolium ring with a triazole.(17) However, phosphorylation or the addition of phosphate-mimicking groups was required for *in vivo* activity.(20) Recently, Zeller *et al*. showed that the low nanomolar affinity (*K_D_* = 19 nM) of a close analog of TPP is achieved when thiamine (*K_D_* = 11 µM) is linked with the pyrophosphate analog MDP (*K_D_* = 1.2 mM), in spite of a thermodynamically unfavorable structural change of the RNA.(21, 22) The same group used SHAPE-MaP RNA structure probing to identify two thiM aptamer binding fragments that are structurally unrelated to TPP (22) resulted in a novel drug-like ligand with 620 nM affinity, without fully reaching to the PP helix, and interacting with a changed conformation of the aptamer pocket.

So far, efforts to explore new chemical matter for the thiM aptamer were focused either on examining the SAR around TPP with close analogs or on fragment-based approaches. Here we develop novel druglike replacements of TPP as thiM ligands with affinities in the low nanomolar range, using structure-guided computational design. Designs followed our main hypothesis that ligands need to interact efficiently with both the Py- and the PP-helices, to achieve high affinity (Figure 1). Initially, analogs targeting the Py-binding pocket were rationally designed. Suitable linker structures connected to heterocycles that reach into the PP-helix were developed and combined with the Py-binding fragments. The resulting nonnative ligands bind the TPP aptamer with affinities up to 1.0 nM, are cell-permeable and show improved pharmacological properties. We show that these analogs induce gene expression controlled by artificial TPP riboswitches in mammalian cells. Moreover, strategic mutations of the pyrophosphate binding pocket of the aptamer allowed us to modify its binding properties by improving its selectivity for the ligand analogs versus TPP. Eventually, truly orthogonal RNA/ligand pairs that are characterized by binding affinities similar to the naturally evolved interaction, were obtained. This work demonstrates the feasibility of re-engineering naturally occurring RNA/ligand pairs to enlarge the repertoire potentially suitable for biotechnological or therapeutic applications

## Results

### Structure-guided design of drug-like TPP riboswitch modulators

The binding mode in the published co-crystal structure of TPP bound to the thiM aptamer (PDB code: 2GDI) (10) served as a starting point for the design of drug-like thiM aptamer ligands. Our efforts were guided by the available knowledge of the relevance of the different parts of the TPP molecule for thiM binding (17, 18, 21), and by predicted ADME/PK properties (e.g. membrane permeability, clogP and metabolic stability). Property predictions were done with machine learning models established at Boehringer Ingelheim ((23); see SI for details). We measured binding affinities to the thiM aptamer with Surface Plasmon Resonance (SPR) spectroscopy (24). A thiM aptamer with a point mutation (G40C, Figure 1) served as a control to identify molecules that bind specifically to the Py-helix (15) and do not bind non-specifically to RNA. (Table 1).

**Table 1:**
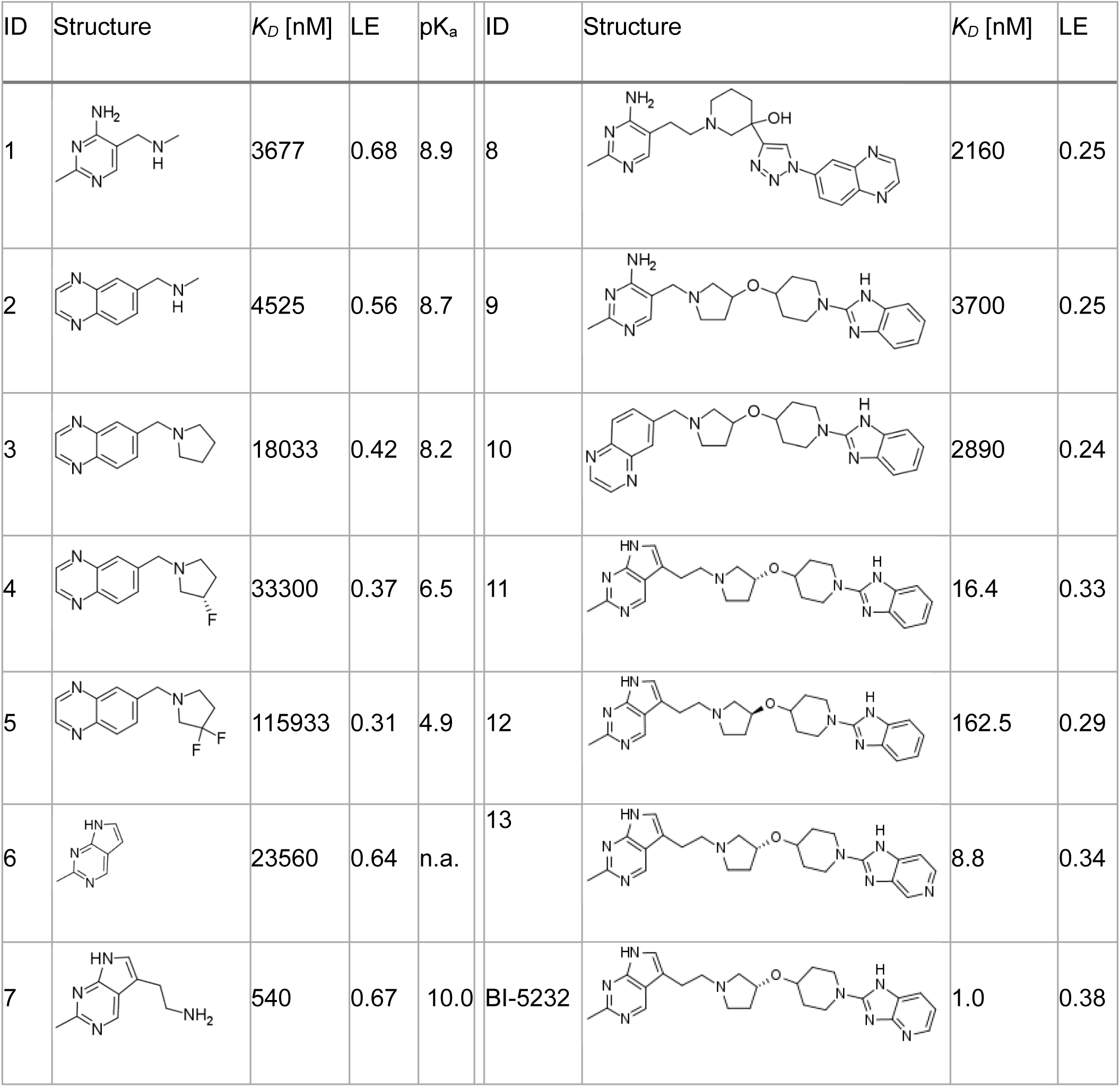
Binding affinities of head groups and of designed thiM aptamer modulators. *K_D_*: Dissociation constant to the wild type thiM aptamer, determined by SPR (*K_D_* > 200 µM for G40C mutant for all compounds). LE: Ligand efficiency (LE = 1.4 x (-log *K_D_*)/[heavy atoms]). pK_a_: Calculated pK_a_ values of the basic amine.

| ID | Structure | $K_D$ [nM] | LE | $\text{pK}_a$ | ID | Structure | $K_D$ [nM] | LE |
| --- | --- | --- | --- | --- | --- | --- | --- | --- |
| 1 |  | 3677 | 0.68 | 8.9 | 8 |  | 2160 | 0.25 |
| 2 |  | 4525 | 0.56 | 8.7 | 9 |  | 3700 | 0.25 |
| 3 |  | 18033 | 0.42 | 8.2 | 10 |  | 2890 | 0.24 |
| 4 |  | 33300 | 0.37 | 6.5 | 11 |  | 16.4 | 0.33 |
| 5 |  | 115933 | 0.31 | 4.9 | 12 |  | 162.5 | 0.29 |
| 6 |  | 23560 | 0.64 | n.a. | 13 |  | 8.8 | 0.34 |
| 7 |  | 540 | 0.67 | 10.0 | BI-5232 |  | 1.0 | 0.38 |

A significant part of our efforts to design more drug-like ligands was based on replacement of the highly charged pyrophosphate in TPP which significantly impairs cell permeability of the active ligand. The network of highly polar Mg-ion and water mediated interactions of the pyrophosphate with the PP helix (10, 13, 19, 22, 25) is essential for high affinity binding of TPP to the aptamer, with measured dissociation constants of thiamine monophosphate and thiamine increasing by approximately 1.4 and 3 orders of magnitude respectively (supplementary information, table S3) versus TPP. We recognized there would be a significant challenge to finding nonionizable replacements for the pyrophosphate but hypothesized that a neutral aromatic ring system may be able to intercalate in the nearby RNA base stacking and efficiently replace interactions of the pyrophosphate with the inner surface of the PP-helix.

The other key design aspect was the identification of an alternative linker between the Py-helix binding head group and the planned intercalating group which would replace the permanent positively charged thiazolium of TPP which we viewed as an additional potential liability to cell permeability. We had envisioned potentially using a triazole group as part of the linker to enable efficient exploration through click chemistry. However, our SPR data for both the isosteric uncharged triazole and thiophene analogs of thiamine (compounds S4 and S5 in Table S3) respectively showed a 20-fold and > 70-fold reduction in binding affinity. These data agreed with similar findings by Chen et al. showing nearly 100-fold and 300-fold reductions in binding affinity for the triazole and thiophene pyrophosphate bearing analogs of TPP (17). These findings in conjunction with the structural data, wherein the TPP thiazolium nitrogen is 3.6 Å from the negatively charged phosphate of C73, confirm the importance of a positively charged group appropriately positioned in the linker. As such, we hypothesized linkers bearing appropriately placed basic aliphatic amines could lead to improved permeability without a significant negative impact on binding affinity and incorporated this in our hit identification plan. Ultimately, we identified two head groups as foundations for virtual screening of potential ligands. In addition to the natural 5-aminomethyl-2-methyl-4-amino pyrimidine fragment derived from TPP (compound 1, table 1) we selected an additional potential head group based on 6-aminoquinoxaline, identified as highly efficient binder (*K_D_* = 22 mM, LE = 0.58) in a fragment screen reported by Cressina *et al.* (18). We modified the fragment hit and selected 6-aminomethylquinoxaline (compound 2, table 1) based on a binding model introducing a basic amine which would a.) place the positive charge in a similar location to TPP and b.) provide a suitable vector to grow towards the PP helix. We found both of these fragments to be highly ligand efficient binders to the thiM aptamer with the thiamine derived fragment with binding affinities of 3.7 and 4.5 µM respectively for compounds 1 and 2. In order to further test whether the aminomethyl of compound 2 might have a similar electrostatic interaction to the thiazolium of TPP, we synthesized a series of pyrrolidine containing analogs of compound 2 with fluorine substituents to modulate basicity (des-fluoro, 3-fluoro and 3,3-difluoropyrrolidnes, compounds 3-5 in Table 1). While the pyrrolidine was a less potent and efficient fragment, these showed a clear dependence of binding affinity on pK_a_ of the basic amine, suggesting similar placement to the thiazolium.

As basis for a computational screen, we designed two initial virtual structures into the aptamer binding pocket that contained the key structural elements desired based on our working hypothesis and on our findings (Figure 2): The quinoxaline head group was combined with a linker with a basic motif, leading to a terminal aromatic system for insertion into the PP helix. Two different molecules were used to allow two separate ways of projecting into the PP helix. To address the question of which region of the PP helix might be most suitable for insertion, we selected two different regions to increase our chances of success. These two initial structures served as “seeds” for computational bioisostere searches for drug-like and synthetically accessible linker structures (software Spark (25), see Experimental section in supplementary material). The search hits included 368 compounds from the Boehringer Ingelheim BICLAIM database which includes validated information on synthetic accessibility.(26, 27) Their PK/ADME properties were predicted, and visual inspection led to a selection of 44 structures. The bound conformations of these were assessed by energy minimizations in absence of the receptor, and in some cases with torsion profile calculations. This inspired the synthesis of three compounds (8-10), with K_D_ values in the single digit µM range.

**Figure 2.**
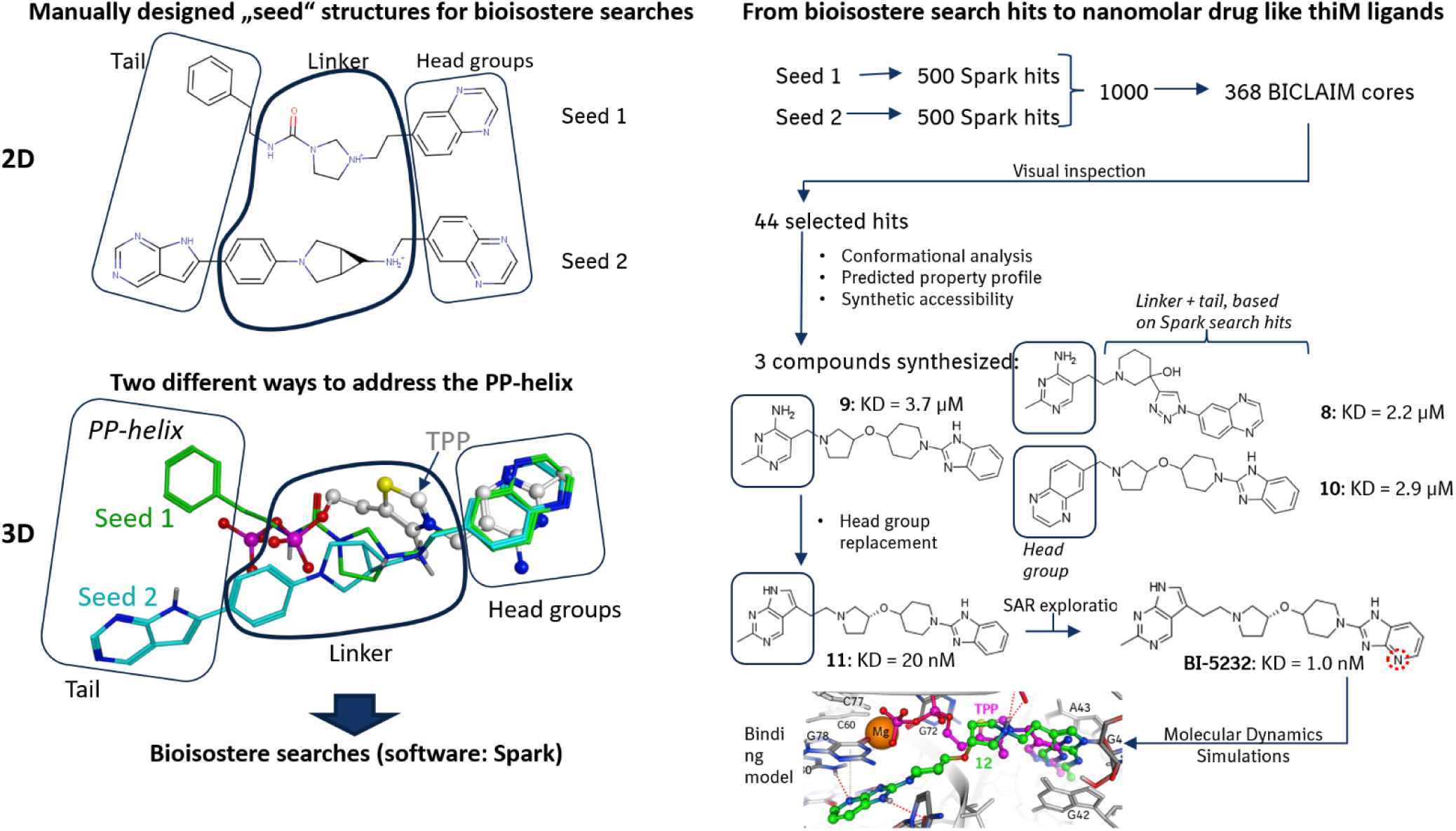
Overview of the computational design approach leading to BI-5232. Two “seed” structures were built into the TPP binding pocket of the thiM aptamer, as observed in X-ray structure of Serganov et. al (PDB code 2GDI (10), left hand side panel): Seed structures 1 and 2 offered the possibility to project aromatic ring systems into two different regions of the pyrophosphate (PP) helix of the aptamer. Right hand side panel: These two seeds served as starting points for computational bioisostere searches which led to the synthesis of 8, 9 and 10. Installing a new head group afforded a > 100-fold increase in binding affinity (compound 11). SAR exploration of the “tail” group resulted in BI-5232 with a KD value of 1.0 nM. A binding model for BI-5232 was derived that is in line with SAR observations and that proved stable in Molecular Dynamics simulations (see Figure 4 for details).

Through a parallel effort to identify alternative head groups we synthesized compound 6, a bicyclic derivative of the TPP pyrimidine head group with excellent binding efficiency (LE = 0.64, Table 1). Addition of a basic ethylamine substituent to mimic the aminomethyl of our other fragments (compound 7, table 1) led to significantly improved affinity (*K_D_* = 540 nM) with a modest gain in ligand efficiency (LE = 0.67). A direct substitution of our original head groups in compounds 9 and 10 as well as identification of the eutomeric linker afforded compound 11, containing an ionizable amine as a thiazolium mimetic but with the anionic pyrophosphate replaced by an uncharged moiety. This compound provided a significant improvement in binding affinity and ligand efficiency over the screening hits (*K_D_* = 16.4 nM, LE = 0.33) and an affinity only one order of magnitude lower than the natural ligand. The dystomer, compound 12, showed an 8-fold reduced affinity (*K_D_* = 163 nM; LE = 0.29).

As a final step in modulating binding affinity and properties, we turned our focus to the benzimidazole tail. Assuming this group behaves as hypothesized, we felt we might further refine potency through either improving the π-stacking or identifying additional interactions (hydrophobic, hydrogen-bonding, etc.) to the aptamer or both. It has been shown that molecular interaction energies can be influenced significantly by dipolar interactions between aromatic rings.(28) We were gratified to find that both aza-benzimidazole isomers further improved binding affinity, with compound 13 showing a 2-fold improvement (*K_D_* = 8.8 nM, LE = 0.34) and BI-5232 leading to a 20-fold improvement in potency and significant increase of the ligand efficiency (*K_D_* = 1.0 nM, LE = 0.38). BI-5232 is a drug-like thiM aptamer ligand with an affinity nearly as good as the natural ligand, in absence of the negative charges of the phosphate moiety and of the permanent positive charge of its thiazolium ring.

The impact of the property guided design is obvious from a direct comparison of BI-5232 with TPP (Figure 3): The polar surface area (PSA) is drastically reduced in BI-5232, and the clogD was raised to the druglike value of 1.8. As a result, we found low to moderate membrane permeability in MDCK PGP cells for BI-5232. The overall drug-likeness was quantified by the central nervous system multiparameter (CNS-MPO, (29, 30)) and by the quantitative estimation of drug-likeness (QED, (31)) scores. BI-5232 shows strong improvements in both regards, approaching the >0.5 threshold of most oral drugs in case of the QED score, and falling in the desirable range of >4.0 in case of the CNS-MPO score.

**Figure 3.**
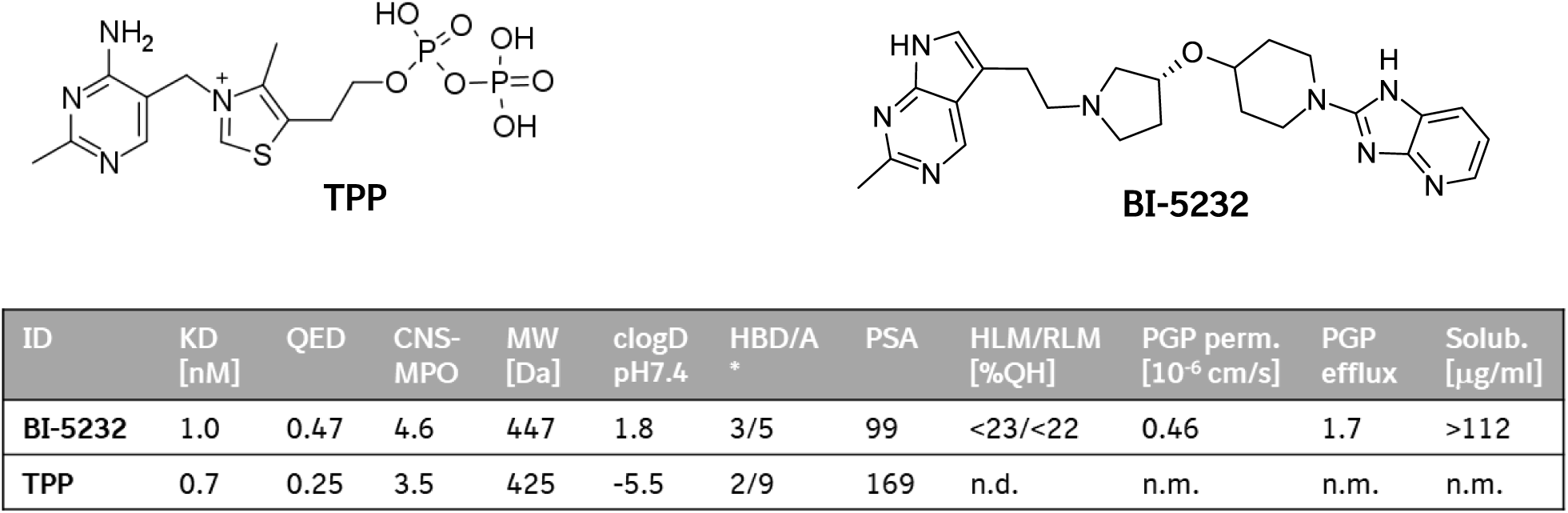
Property comparison of TPP and BI-5232. n.d. = not determined; n.m. = no valid measurement: the high polarity of TPP prevented analytical detection. Column headers: *K_D_* =dissociation constant, as measured by SPR; QED=quantitative estimation of drug-likeness (31); CNS-MPO= central nervous system multiparameter score (29, 30), MW=molecular weight; clogD=calculated logD value at pH7.4; HBD/A=number of hydrogen bond donors/acceptors; HLM/RLM=human/rat liver microsome stability; PGP perm.=membrane permeability in PGP overexpressing cells; PGP efflux=efflux ratio for PGP over-expressing PGP cells; Solub.=solubility.

We explored different binding hypotheses for BI-5232 in the thiM aptamer based on the X-ray structure of the aptamer TPP complex (PDB code 2GDI, (10)), with the benzimidazole derived ring system directed to the region of the PP-helix that interacts with the pyrophosphate of TPP (i.e. G76-G78 and G60-G63). None of these initial models proved stable when subjected to Molecular Dynamics (MD) simulations. During our efforts, Zeller et al. (22) published the X-ray structure of the thiM aptamer with the N-demethyl derivative of compound 2 (PDB code 7TZS). There, a partially changed structure of the aptamer binding pocket is observed, due to different conformations of G72 and C55.

This pointed us to a placement of BI–5232 with the bicyclic aromatic ring system in a sub–pocket pre-dominantly formed by G78 – G81 and C55 – C58 (Fig. 4). In this pocket, all three heteroatoms of the ring system are engaged in hydrogen bond interactions (Fig. 4B). The H-bond of the pyridine nitrogen atom with A80 can explain the gain in potency of BI-5232 compared to compounds 11 and 13 (Table 1) which differ from BI-5232 only by the absence or position of this nitrogen atom. This binding model proved highly stable in Molecular Dynamics simulations (see Experimental section/SI for details) and represents our binding hypothesis for BI-5232 in the thiM aptamer.

**Figure 4.**
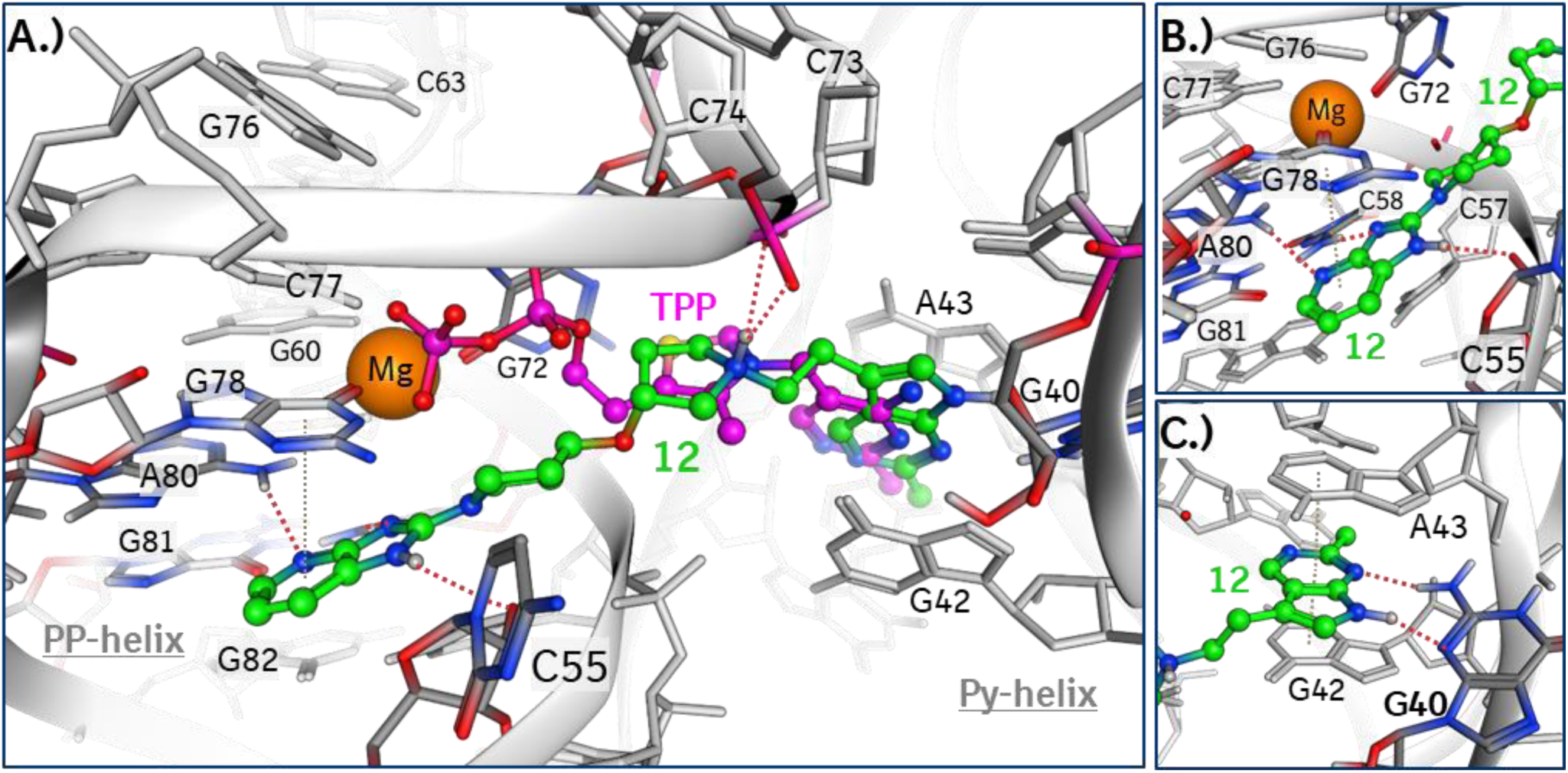
Model of BI-5232 bound to the thiM aptamer, based on 7TZS.pdb (22). **A.)** Comparison with the binding mode of TPP (superposed from 2GDI.pdb, (10)): The pyrophosphate of TPP interacts with the edge of the PP helix base stack via magnesium ion- and water-mediated interactions (not shown), engaging the nucleotides G76-G78 and G60-G63. In contrast, according to our binding model, BI-5232 inserts the bicyclic aromatic ring system into a pocket in the PP-helix region that is predominantly formed by nucleotides not involved in the TPP pyrophosphate interactions (G78 – G81 and C55 – C58). The positive charge of the pyrrolidine ring is interacting with the negatively charged phosphate groups connecting C73 and C74 and is superposing with the positively charged nitrogen of the thiazolium ring of TPP. **B.)** All three nitrogen atoms of the bicyclic ring system are engaged in polar interactions (dashed red lines). The pyridine ring is stacking with the nucleobase of G78 (grey dashed line). **C.)** The head group in the Py-helix shows the typical H-bond interactions with G40 (dashed red lines) and stacking interactions with G42 and A43.

### Evaluation of TPP analogs and generation of orthogonal thiM mutants discriminating against TPP

The iterative rounds of structure-guided design of new TPP analogs and the quantitative evaluation of their affinity towards the thiM aptamer yielded high affinity ligands that bind with dissociation constants in the low nanomolar range. Binding of these molecules depends on the Py-helix, as shown by the deleterious mutation G40C, and they were designed to potentially reach and stack into the PP-helix. In order to validate the actual binding mode of the TPP analogs, in-line probing analysis was performed.

In-line probing relies on the intrinsic instability of RNA phosphodiester bonds in unstructured regions, visualized by radioactive labeling and subsequent PAGE gel analysis.(32, 33) 5’-radioactively labeled thiM aptamer RNA was incubated with TPP and with the most potent TPP compounds 11, 13 and BI-5232. The compounds showed dose-dependent binding to the thiM aptamer while not altering the overall structure of the aptamer (Figure 5A and Figure S6). The band intensity for the L5 region, as well as for the nucleotides 22-26 of the P3 helix decreases with BI-5232 as with TPP (Figure 5A). This indicates that binding of BI-5232 to the thiM aptamer causes the aptamer to adopt the Y-shape that is also observed in the TPP-bound state. For compound 11 and 13 the same effect is visible at higher concentrations (Figure S6). Binding of the compounds 11, 13 and BI-5232 induces a strikingly different band pattern in the PP-helix region compared to TPP. It suggests that the compounds are reaching the PP-helix, as simulated during computational modeling, and bind via a different mode than the parental ligand. In presence of TPP, binding of the pyrophosphate group via coordination of Mg ions results in reduced cleavage around G60 and the P4/5 stem, while compounds 11, 13 and BI-5232 have the opposite effect (Figure 5A). In contrast, the proposed binding mode of the novel compounds involves nucleobases 55-58 reflected by reduced band intensities in the J2-4 loop region of the most potent compound BI-5232 (Figure 5A). A negative control was conducted with the G40C-mutated thiM aptamer. Consistent with the SPR data, no ligand-induced modulation of the in-line probing pattern was observed (Figure S4), showing that binding to the Py-helix is required for both TPP and the new compounds.

**Figure 5.**
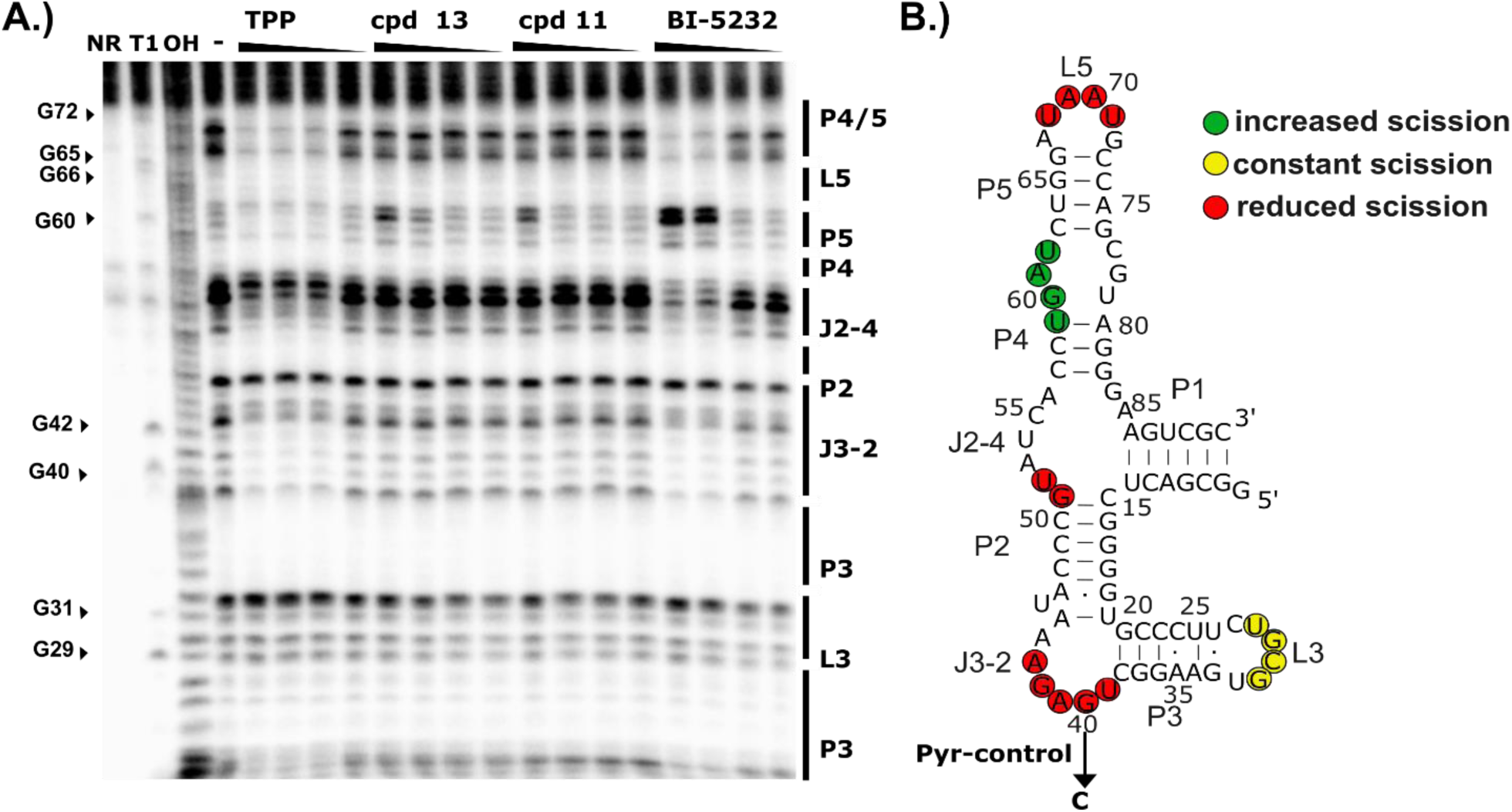
In-line probing of thiM aptamer with TPP and compounds 11, 13 and BI-5232. **A.)** PAGE gel analysis of in-line probing reaction of 5‘-^32^P-labeled TPP thiM aptamer incubated without ligand (-) or with TPP, 13, 11 and BI-5232 (1:5 dilutions from 500 nM). Untreated RNA (precursor, P), T1 RNase (T1) or alkaline digestion (OH) serve as controls. **B.)** Secondary structure of the thiM aptamer. Ligand-dependent increase (green) or decrease (red) of band cleavage in A.) are encircled, additionally highlighted is the PP-binding site (blue box).

The in-line probing data provides evidence that the designed ligands are not only binding tightly to the aptamer but seem to induce that Y-shaped aptamer conformation relevant for altering gene expression in riboswitches.(10, 34) In order to investigate whether the new ligands can be used as triggers of synthetic riboswitches, we engineered a TPP-dependent aptazyme. The architecture of this aptazyme is based on the tetracycline-dependent hammerhead (HHR) ON-switch reported by Beilstein *et al.* in 2015.(35, 36) When placed into the 3’-UTR of a eukaryotic mRNA, the aptazyme cleaves and degrades the mRNA in absence of the ligand while the mRNA is stable in presence of the ligand, turning gene expression on.

We connected the TPP-sensing thiM aptamer to the type III HHR and tested different communication modules (Figure S8). Upregulation of gene expression was observed both in cells cultivated in media supplemented with TPP and thiamine. The latter is taken up by the cells and pyrophosphorylated by thiamine pyrophosphokinase.(37, 38) The switching performance was rather low with an observed 2.1-fold induction at 500 µM thiamine for the ON-switch TPPK4 (Figure S8). We next tested the effect of compound 13 on the TPPK4 switch in a GFP/mCherry reporter system, where mCherry expression serves as transfection control (Figure S10). In this context, the compound induces TPPK4-controlled gene expression (Figure 6B). Importantly, comparable or even exceeding effects are already obtained at 100 µM of compound 13 compared to 500 µM thiamine. Introduction of the negative control mutation G40C in the Py helix abolished riboswitch response for both compound 13 and thiamine (Figure 6B).

**Figure 6.**
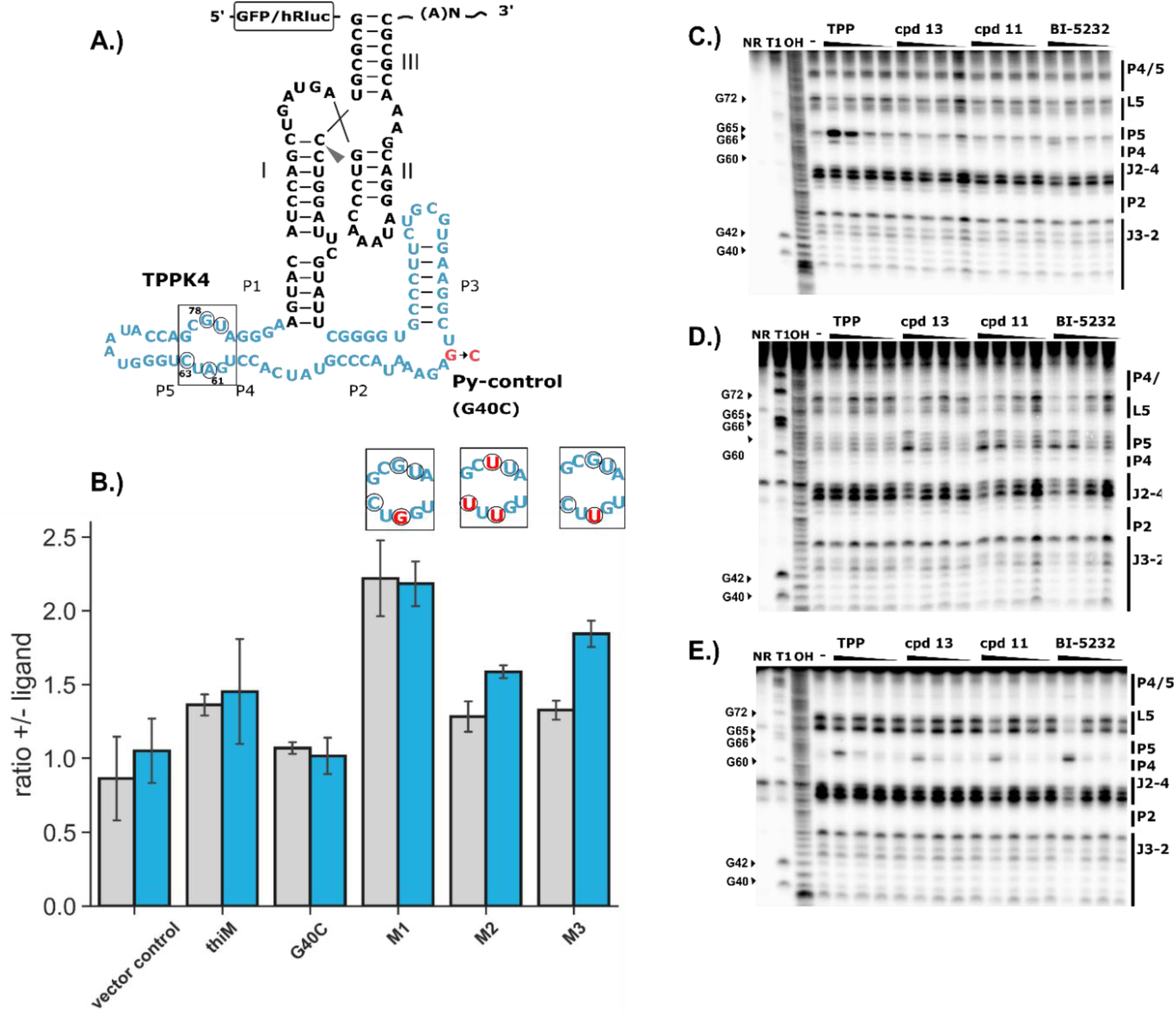
Screening for orthogonal mutants and in-line probing analysis. **A.)** Schematic depiction of the TPP-HHR aptazyme that is inserted into the 3’-UTR of an EGFP gene). **B.)** Gene expression of EGFP in HeLa cells after 3’-insertion of TPP-HHR aptazymes containing the TPP WT aptamer, a binding-deficient TPP aptamer with mutated Py-site (negative control) and mutants (M1-M3) identified in a screen of a library with a randomized PP-binding site. The randomized nucleotides are encircled, mutations are shown in red. Transfected Hela cells were treated with 500 µM thiamine (light gray) or 100 µM compound 13 (blue). Reporter gene expression was determined by flow cytometry 24 h post transfection. EGFP expression was divided by mCherry expression (transfection control), the fold change was determined by normalization on gene expression without thiamine. Error bars represent standard deviations of three independent measurements; each experiment performed in triplicates. **C.)-E.)** In-line probing reaction of 5‘-^32^P-labeled mutant thiM aptamer M1-3 in absence (-) or presence of TPP and compound 11,13 and BI-5232 (5:1 dilutions from 500 nM). Untreated RNA (precursor, P), T1 RNase (T1) or alkaline digestion (OH) serve as controls.

We next aimed to develop riboswitch sequences that are triggered selectively by the novel compounds but not by TPP. As design strategy we reasoned that binding of the novel compounds should be less affected by mutations in the Mg-complexing positions compared to the strict requirement of the conserved PP site for high-affinity TPP binding. We generated a library of TPPK4 HHR switches that contain four randomized positions within the PP-helix that are known to be required for Mg^2+^-mediated TPP binding which also showed a distinctly different modulation upon binding of compounds 11, 12 and BI-5232 compared to TPP (Figure 6A): Nucleotides A61, C63, C77, G78. A61 is coordinating one of the Mg^2+^ ions and forms stacking interactions with C63 and G60 that is as well directly coordinating a Mg^2+^-ion. Additionally, A61 is forming a H-bond to the oxo group of C77, which is stacking between G76 and G78 and thereby stabilizing the overall structure of the PP binding pocket. C77 and G78 are forming hydrogen bonds with the β-phosphate of TPP.

We screened the TPPK4 HHR library in HeLa cells as described before.(39) Several sequences that respond to compound 13 but dońt show changes in gene expression upon thiamine addition were identified (Figure 6B). We validated three of the most interesting mutants, confirming the initial results (Figure 6B). Interestingly, a mutation at nucleotide 61 (M1, A61G) did not reduce but rather enhance the switching efficiency with TPP and likewise with compound 12. Two mutants (M2, M3 Fig 6B) showed higher induction of gene expression upon compound addition. Aptamer M2 contains 3 mutations (A61U, C63U, G78U), while the difference in the ligand response of mutant 3 seems to rely solely on a single point mutation (A61U). Since the mutants 1-3 showed ameliorated response to the novel compounds, we analyzed the ligand binding properties of all three aptamers with TPP and the most promising compounds 11, 13 and BI-5232.

In order to investigate the interaction of the re-engineered aptamer sequences with the designed ligands we performed in-line probing experiments with the mutant thiM aptamers with compounds 11, 13 and BI-5232. (Figure 6C-E, Figures S5 and S7). All mutants show a decreased TPP-dependent structural modulation compared to the wild type thiM aptamer. The mutated aptamers retained the general thiM fold and upon compound addition showed enhanced band intensities in the PP-helix region while the band intensities decreased in the Py-helix and the regions in P3 and L5 reflecting the adoption of the Y-fold. The mutants mostly differ in the band pattern of the PP-helix and in the extent of compound-induced modulation of this area. Different than in the in-line probing of the WT aptamer (Figure 5A), M1 and M3 display a prominent band corresponding to the base 61 at higher concentrations of TPP and the compounds, indicating less protection at this position which might be caused by deficient Mg complexation (Figure 6C-E). In general, the change in the band pattern occurred at higher TPP concentrations compared to the developed compounds, indicating a more selective recognition of compounds 11, 13 and BI-5232.

The results were confirmed by SPR analysis of the mutant aptamers (Figure 7B-C and Table 2). Series of concentrations of TPP and compounds 11, 13, BI-5232 were measured and fitted to a 1:1 kinetic binding model. As reported before, the affinity for TPP of the wild type thiM aptamer was determined in the low nanomolar range.(18, 40) According to our measurements (Table 2) the mutant M2 binds TPP with a more than 900-fold reduced affinity (*K_D_*: 687 nM, vs. 0.74 nM for the natural aptamer) but retains high affinities for all three novel ligands. Compound 13 is bound by M2 with a *K_D_* of 1.8 nM which is an >4-fold improvement compared to the affinity towards the wildtype thiM sequence of *K_D_* = 8.8 nM. Also, for compound 11 that binds wildtype thiM with a *K_D_* of ∼ 16 nM the affinity is increased by a factor of ∼25 to a *K_D_* of 0.6nM, while BI-5232 is bound by the wildtype and mutated aptamer with similar affinities (M2 *K_D_*: 2.0 nM vs WT *K_D_*: 1.0 nM). The mutant M3 results in the most drastically reduced affinity for TPP (*K_D_*: 2200 nM), but only slightly lowered affinity for the analogs 11 and 13. Notably BI-5232 remains an efficient binder of M3 (*K_D_*: 2.5 nM).

**Figure 7.**
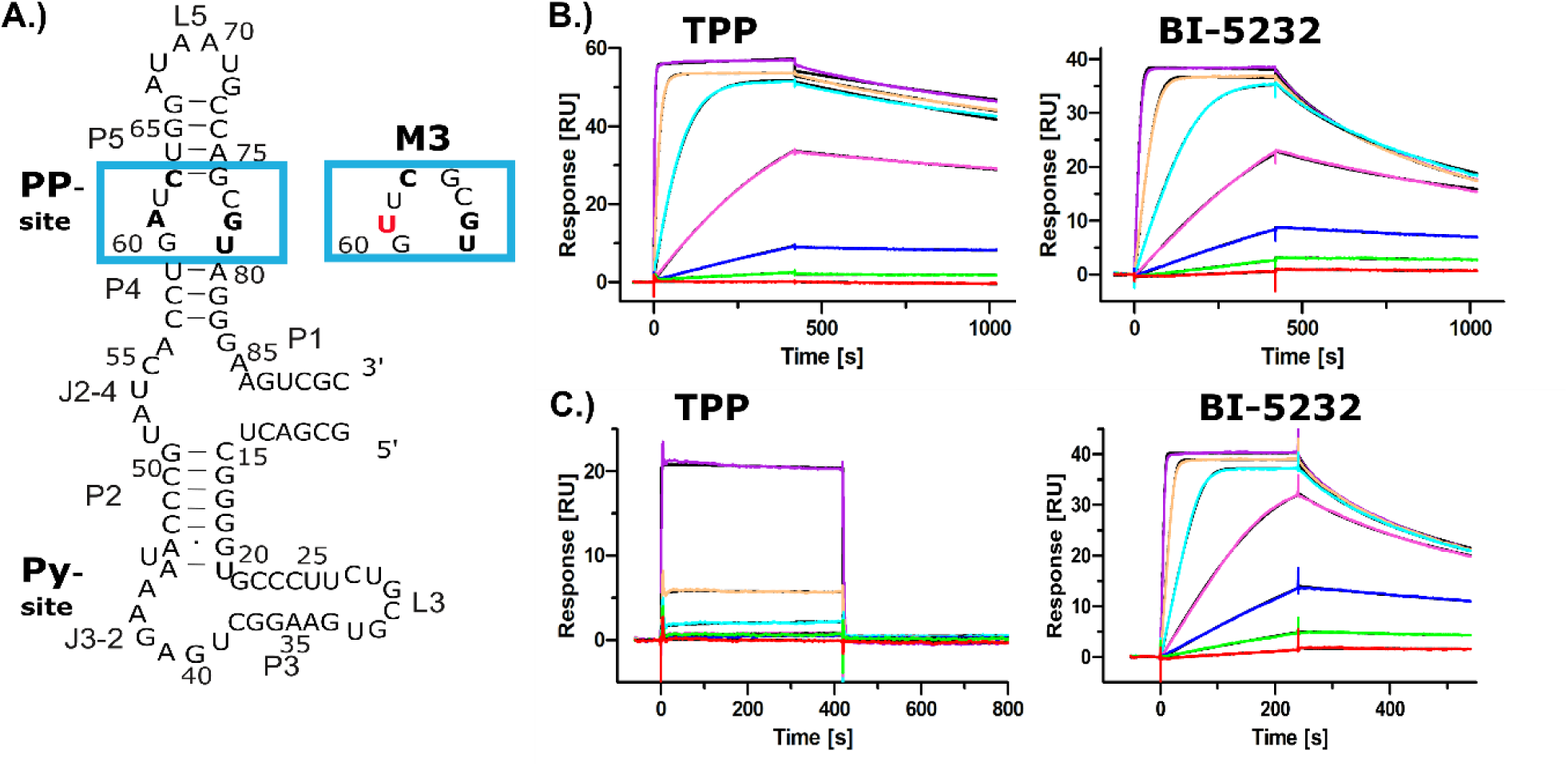
SPR binding studies of mutant 3 and thiM WT aptamers. **A.)** Secondary structure of the thiM aptamer. are encircled, additionally highlighted is the PP-binding site, the mutated version (A61U, M3) is shown on the right (blue box). SPR measurements: varying concentrations of TPP (left) and BI-5232 (right) bind to to thiM WT RNA **B.)** or mutant M3 RNA **C.)**. Aptamers were hybridized to a DNA-linker that was immobilized on a CM5 sensor chip via amine-coupling. The sensorgrams were fitted to a 1:1 kinetic binding model, resulting *K_D_* values are shown in Table 2.

**Table 2:** Binding affinities of TPP and of compound 11, 13 and BI-5232 for the wild type thiM aptamer and mutants M1-3. The dissociation constants (*K_D_*) were determined by SPR (additional details can be found in Table S2).

| Aptamer | $K_D$ [nM]<br>TPP | $K_D$ [nM]<br>cpd 11 | $K_D$ [nM]<br>BI-5232 | $K_D$ [nM]<br>cpd 13 |
| --- | --- | --- | --- | --- |
| thiM | 0.74 | 16.4 | 1.0 | 8.8 |
| Mutant 1 | 233 | 41 | 9.8 | 10.8 |
| Mutant 2 | 687 | 0.6 | 2.0 | 1.8 |
| Mutant 3 | 2200 | 49.7 | 2.5 | 34.1 |

Surprisingly also M1 has a reduced affinity for TPP, although improved switching was observed for the M1-riboswitch version, suggesting that the mutations have an additional effect on aptamer folding and switching into the ligand-bound conformation that cannot be explained by affinity alone. Taken together these results indicate a successful re-engineering of the thiM aptamer, creating orthogonal mutant aptamers that selectively bind the non-native ligands with high affinity and in some cases discrimination of TPP by a factor of more than 100-fold (M3, BI-5232).

Since M3 displays the lowest affinity for TPP while maintaining high affinity for BI-5232, we selected this combination in order to investigate whether the selectivity is retained in the synthetic HHR-based riboswitch. A concentration-dependent increase of reporter gene expression of a constructed TPPK4-M3 HHR riboswitch in a dual luciferase vector (Figure 8A-B and Figure S11) was observed for BI-5232 whereas no induction was obtained with thiamine (Figure 8B). This result demonstrates that the screened mutant aptamer M3 can indeed be utilized to construct an orthogonal, re-engineered riboswitch selective for BI-5232. Exploiting the modularity of riboswitch designs, we next investigated whether the selective aptamer can be adopted in a further aptazyme design. For this purpose, we connected the thiM M3 mutant aptamer to a HHR type I ribozyme from *Schistosoma mansonii*, creating an OFF-switch, similar to previous TPP riboswitch designs (Figure 8A).(41, 42) For this riboswitch, concentration-dependent downregulation of the luciferase reporter gene expression was observed upon increasing concentrations of BI-5232 whereas no effect was observed with thiamine (Figure 8D).

**Figure 8.**
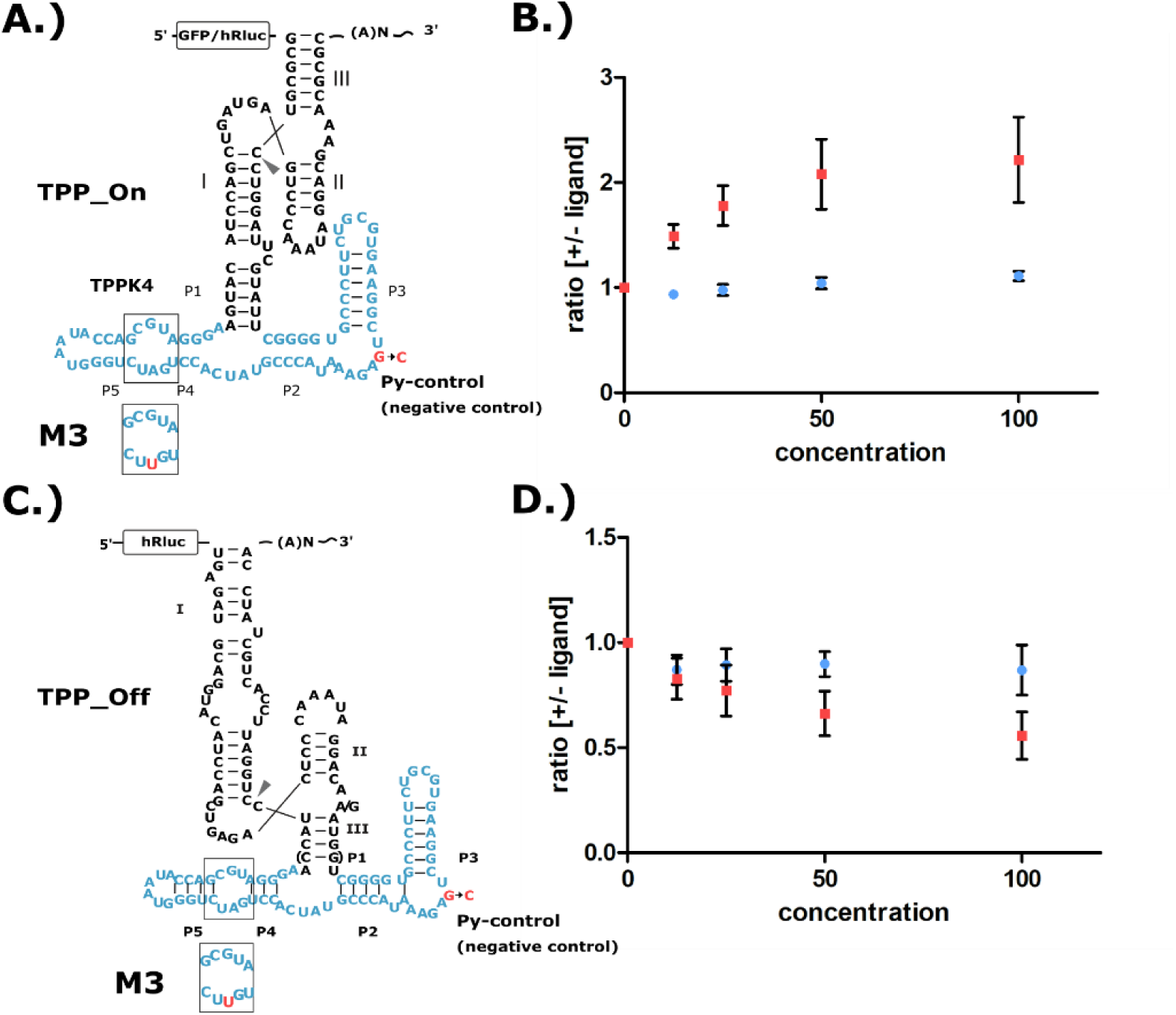
Reporter gene expression control of a luciferase reporter in HeLa cells by riboswitch mutant M3. **A.)** Sequence and secondary structure of the TPP-HHR ON-switch and PP-mutation of M3. **B.)** Dose-dependent gene expression in presence of BI-5232 and thiamine after transient expression with plasmids, expressing a Luciferase reporter under the control of mutant M3. Shown is the ratio of reporter gene expression in presence vs. absence of the respective ligand, BI-5232 (red) and thiamin (blue). **C.)** Schematic representation of the TPP-HHR OFF-switch design, showing the aptazyme sequence and **D.)** Concentration-dependent effect of BI-5232 (red) in comparison to thiamin (blue). Luminescence was measured 24 h after transfection. Error bars represent standard deviations of three independent experiment.

## Discussion

Riboswitches as tools for gene expression control hold great potential for medical applications in gene therapies, but aptamers found in nature often bind small molecules that occur in all cell types. In this work, we describe the development of novel, highly affine ligands for the natural thiM aptamer. We show that these ligands are inducing structural changes in the aptamer comparable to TPP and can thus trigger synthetic riboswitches engineered to function in mammalian cells. Although the compounds were designed to bind the aptamer in a similar manner as TPP, their chemical structures differ greatly. Moreover, analysis of their binding mode via in-line probing and a subsequent small-scale screen allowed us to develop aptamer mutants with drastically reduced affinities for TPP that still bind the developed compounds with high affinity. Finally, we designed orthogonal riboswitches that control gene expression in mammalian cells in response to the new compounds but do not respond to thiamine.

To date, attempts to enlarge the repertoire of riboswitches mainly focus on the de-novo generation of aptamers via SELEX, starting with pre-defined ligands. The *in-vitro* selected tetracycline aptamer, published by Berens *et al*. in 2001 turned out to meet the structural requirements to trigger riboswitches on several expression platforms in mammalian cells.(35, 43-45) Subsequent studies showed that these switches are also functional in *C. elegans* and allow the induction of adeno-associated viral vector (AAV) transgene expression in mice, underlining the general utility of riboswitch-application for gene therapies.(2, 44) Apart from this, several RNA aptamers with high affinities towards novel ligands were developed via SELEX-campaigns, e.g., for theophylline and doxycycline.(46, 47) Recently, Fukunaga *et al.* selected an aptamer for a derivative of potassium channel inhibitors that was suitable for the design of efficient synthetic riboswitches.(48) However, the generation of functional riboswitches based on most of these aptamers turned out to be challenging, due to insufficient structural rearrangements upon ligandbinding. To address this issue, Hoetzel and Suess proposed a combination of capture-SELEX, where the ligand is added for RNA elution, confirming the selection of aptamers that undergo conformational changes during ligand-binding, and in vivo screens to ultimately identify aptamer sequences that are suitable for the construction of efficient riboswitches.(49, 50)

Here, we show that redesigning natural riboswitch-ligands and subsequent modification of the binding domain results in novel aptamer-ligand combinations that can be directly used for the design of artificial riboswitches. Exploiting natural riboswitches holds the advantage that their aptamers have evolved for efficient structural changes upon ligand binding, which makes them more suitable for controlling expression platforms. Using a similar approach, Dixon *et al.* reengineered the *add A* riboswitch, which resulted in bacterial riboswitches with higher response for ammeline and azacytosine instead of their natural ligand, adenine.(51) So far, the design and identification of alternative riboswitch ligands have mostly been viewed as a strategy for discovering new antibiotics to disrupt the natural riboswitch function.(6, 52) In contrast, the herein developed ligands were designed to bind the aptamer as TPP, in the Py-binding pocket and the P4/5 stem, causing aptamer-folding similar to the natural ligand. Therefore, the ligands were likely to induce the structural change that is necessary to trigger switching on an expression platform. In fact, BI-5232 seems to induce aptamer folding more effectively than TPP, pointing out an alternative potential of riboswitch optimization via aptamer mutation of the J2-4 region. Moreover, even minor changes in the sequence were sufficient to reengineer the aptamer efficiently. Such aptamer variations can be identified easily in low throughput screening approaches.

The developed artificial TPP aptazymes possess only moderate switching efficiency. Potentially the structural change of the TPP-aptamer is not ideal for this type of riboswitch construct. The TPP response of both M3-riboswitches is much less pronounced, which might be a consequence of the strongly reduced TPP affinity of mutated aptamer sequences. However, the dynamic range does not represent the ligandbinding affinity, as the mutations might also influence the structural rearrangements during aptamer folding.

The mutated aptamer was utilized for construction of two different types of riboswitches with ON- and OFF-reactivity, demonstrating that it is generally functional in the mammalian cellular environment. In addition, several other artificial riboswitch platforms have been developed recently, including regulatory elements influencing splicing, polyadenylation, RNA interference or translational frameshifting.(44, 45, 53-57) The continuing development of such novel expression platforms opens several novel design strategies for the implementation of orthogonal ligand/aptamer pairs such as the ones introduced here.

Complementing former efforts to identify drug-like TPP replacements for the thiM aptamer, which were focused fragment-based approaches (see introduction), we used a structure guided computational design approach. Our working hypothesis was that ligands need to interact efficiently with both, the Py- as well as the PP-helices, to achieve high affinity, and that the highly polar pyrophosphate of TPP can be replaced with a heterocyclic ring system that forms stacking interactions with the PP-helix. With BI-5232 we achieved 1 nM binding affinity with a molecule that shows excellent drug-like properties (Figure 3).

We propose a binding model for BI-5232 that is supported by our SPR data and by MD simulations. In this model, the interaction with PP helix occurs not in the region that is of prime importance for the binding of the pyrophosphate group of TPP. Key polar interactions are instead formed by A80, C58 and C55. This is in line with the observation that M1 (A61G), M2 (A61G+C63U+G78U) and M3 (A61U) show no major effect on the binding affinity of the novel ligands.

It should be noted that initial binding models for BI-5232 were based on the thiM aptamer structure obtained in complex with TPP by Serganov et al. (PDB code 2GDI, (10)). These models proved to be unstable in MD simulations. The proposed binding model was built using the more recent structure of Zeller et al. (PDB code 7TZS, (22)) This structure contains a small ligand that interacts only with the Py-helix. This leaves room for the nucleobase of G72 to move into the pocket region that in case of TPP is occupied by the pyrophosphate. Similar conformations of G72 are observed in all four structures of Zeller et al., all containing ligands that do not reach the PP helix. In our binding hypothesis the nucleobase of G72 provides an important interaction surface for the linker region of BI-5232. We therefore propose that the conformation of the thiM aptamer binding pocket in absence of TPP, as found in the recent X-ray structures by Zeller et al., provides a superior basis for structure-based design of TPP riboswitch modulators.

Tran *et al*. also replaced a phosphate of the natural nucleotide ligand (ZMP) of the ZTP riboswitch by heterocycles forming stacking interactions with RNA nucleobases.(58) They arrived at ligands with affinities similar to the natural ligand, in this case in the low µM range. A group at Merck identified ribocil as a drug-like ligand with low nanomolar affinity of the FMN riboswitch using a phenotypic screening approach.(58, 59) To our knowledge, this work for the first time shows the viability of structure-based rational design of drug like riboswitch modulators rivaling the affinity of natural ligands in the low nanomolar range. Our designed tool compound, BI-5232, is freely available to the scientific community via the OpnMe platform.(60, 61) We hope that it will be instrumental to the scientific community in exploring controlled gene expression in combination with thiM aptamer derived riboswitches.

### Experimental Section

The experimental section can be found in the supporting information of this article.

## ASSOCIATED CONTENT

Experimental section

SI Figure 1: Stability of the thiM aptamer structures during the simulations

SI Figure 2: Stability of the ligand conformations during the simulations of the complexes with the thiM aptamer

SI Figure 3: Hydrogen bond interaction count between the thiM aptamer and TPP and BI-5232, as observed during the simulations

SI Figure 4: PAGE-gel analysis of in line probing with G40C mutant (negative control)

SI Figure 5: PAGE gel analysis of in-line probing reactions of thiM mutants M1 and M2

SI Figure 6: PAGE gel analysis of in-line probing reactions of thiM WT aptamer

SI Figure 7: PAGE gel analysis of in-line probing reaction of the mutant aptamer M3

SI Figure 8: Generation of TPP-responsive ON-switching aptazymes on the basis of the TetKx-platform

SI Figure 9: Screening of TPPKx aptazymes with mutated PP-helix.

SI Figure 10: Plasmid map of EGFP-mCherry vector used to test TPPKx aptazymes in combination with the investigated analogs.

SI Figure 11: Plasmid map of luciferase expressing vector used to determine activity of TPPKx aptazymes.

SI Table 1: Primer sequences for TPP riboswitch insertion in the 3’-UTR of reporter genes

SI Table 2: SPR data of TPP WT aptamer and aptamer mutants

SI Table 3: SPR binding constants (K_D_) for TPP (1) and derivatives Synthesis of compounds

This material is available free of charge via the Internet at http://pubs.acs.org.

## AUTHOR INFORMATION

### Corresponding Authors

Jörg S. Hartig; Tel: +49 7531 88 4575;

Oliver Hucke - Boehringer Ingelheim Pharma GmbH & Co. KG, 88397 Biberach an der Riß, Germany;

### Notes

The authors declare no conflict of interest.

## ACKNOWLEDGMENT

We thank Dmitry Galetsky, Astrid Joachimi and Yvette Hoevels for technical assistance.

