## Supplementary Information for "Small molecule-controlled gene expression: Design of drug-like high affinity small molecule modulators of a custom-made riboswitch"

### Table of content

1. Experimental Section
1.1. Structure guided design 3
1.1.1. Molecular Modeling 3
1.1.2. Bioisostere searches 3
1.1.3. Torsion profiles 3
1.1.4. Prediction of compound properties 3
1.1.5. Molecular Dynamics simulations 3
1.2. In-Line probing 4
1.3. Plasmid construction 4
1.4. Cell culture experiments 4
1.5. Flow cytometry 4
1.6. Surface Plasmon resonance 4

2. Supporting Figures 5

SI Figure S1: Stability of the thiM aptamer structures during the simulations 5
SI Figure S2: Stability of the ligand conformations during the simulations of the complexes with the thiM aptamer. 6
SI Figure S3: Hydrogen bond interaction count between the thiM aptamer and TPP and BI-5232, as observed during the simulations. 6
SI Figure S4: PAGE-gel analysis of in line probing with G40C mutant (negative control) 7
SI Figure S5: PAGE gel analysis of in-line probing reactions of thiM mutants M1 and M2 7
SI Figure S6: PAGE gel analysis of in-line probing reactions of thiM WT aptamer 8
SI Figure S7: PAGE gel analysis of in-line probing reaction of the mutant aptamer M3 8
SI Figure S8: Generation of TPP-responsive ON-switching aptazymes on the basis of the TetKx-platform 9
SI Figure S9: Screening of TPPKx aptazymes with mutated PP-helix. 10
SI Figure S10: Plasmid map of EGFP-mCherry vector used to test TPPKx aptazymes in combination with the investigated compounds 10
SI Figure S11: analogs Plasmid map of luciferase expressing vector used to determine activity of TPPKx aptazymes 12

3. Supporting Tables
SI Table S1: Primer sequences for TPP riboswitch insertion in the 3'-UTR of reporter genes 13
SI Table S2: SPR data of TPP WT aptamer and aptamer mutants 14
SI Table S3: SPR binding constants ($K_D$) for TPP (1) and derivatives 14

4. Synthesis of compounds 15

5. HPLC traces of key compounds

6. $^1$H NMR of key compounds

7. Chiral chromatography for compounds 11 and 12

8. References 34

#### 1. Experimental section

##### 1.1. Structure guided design

###### 1.1.1. Molecular Modeling of “seed” structures for bioisostere searches

The two “seed” structures used as input for subsequent bioisostere searches were obtained on basis of the X-ray structure of the complex of TPP with the thiM aptamer published by Serganov et al. (PDB code: 2GDI,[1]). They were designed in the TPP binding cavity of the aptamer using the “Builder” module of the software “Molecular operating environment (MOE)”.[2] The modeling was carried out using the MMFF94x forcefield and R-Field electrostatics for minimizations.

###### 1.1.2. Bioisoster searches

Bioisoster searches were conducted with the software Spark.[3] Spark searches databases of molecular fragments for building blocks that are similar in shape and electrostatics to a defined substructure of the starting molecule, and that have a connectivity that is geometrically compatible with the substructure to be replaced. Searches were done in the “Chembl_common”, “Common”, “VeryCommon” and “cores_BICLAIM” databases, using default parameters. The latter is a Boehringer Ingelheim proprietary collection of core reagents that can be chemically enumerated to cover a space of billions of compounds using validated synthetic approaches.[4, 5] The 500 highest ranked hits from the Spark searches were saved for each of the two starting structures (“seed” structures). Within the set of 1000 hits, 368 came from the BICLAIM-cores database. These were prioritized, their PK/ADME properties were predicted, and they were visually inspected, leading to a selection of 44. The conformations of these were assessed by checking for large deviations from the proposed conformation upon energy minimization. In some cases, conformations were checked with torsion profiles. A final list of 23 compounds was subject to selection by the chemistry team. Ultimately, three structures were considered for synthesis. They inspired the synthesis of compounds 8, 9, 10 (Table 1 and Figure 2).

###### 1.1.3. Torsion profiles

Torsion profiles were calculated with Density Functional Theory (DFT) calculations at the wB97XD level of theory using a cc-pVDZ basis set. The software Gaussian 09[6] was used for this purpose.

###### 1.1.4. Prediction of in vitro ADME/PK properties

Boehringer Ingelheim has established a state-of-the-art machine learning platform for the prediction of numerous substance properties that are highly relevant for the suitability of small molecules as therapeutics .[7] Using this platform, the following properties were calculated for all compounds considered for synthesis: Caco2 permeability, Caco2 efflux, MDCK PGP permeability, MDCK PGP efflux, liver microsome stability (human, rat, mouse), plasma protein binding (human, rat, mouse), solubility, cytochrome P450 3A4 inhibition. Further, the following properties were calculated as basis for a combined numerical score to assess suitability for passing the blood brain barrier (CNS MPO score[8]): cLogP, cLogD, most basic pKa, molecular weight, TPSA, number of H-bond donors. The predicted property profile was considered during compound design and for the selection of compounds for synthesis.

###### 1.1.5. Molecular Dynamics Simulations

*Parametrization and System:*
The Molecular dynamics simulations were conducted using the AMBER99SB-ildn Force Field.[9] The ligands to the riboswitch, Thiamine Pyrophosphate (TPP) and BI-5232, were parametrized with OpenFF 2.0.0.[10] A rectangular box with periodic boundaries was employed, and a minimum distance of $1\,nm$ between the complex and the boundary condition was maintained. The system was solvated with explicit TIP3P water.[11] The system was neutralized with sodium ions, which were randomly placed in the solvent. The solvation, box generation and ion neutralization performed with the GROMACS 2021 tools and for the parametrization the OpenFF-toolkit was used.[12, 13]
*Simulation Preparation:*
The GROMACS 2021 molecular dynamics engine was used for all simulation steps.[12] The nonbonded neighbor search was performed grid-based with the Verlet cutoff-scheme, and a short-range neighbor distance limit of $1.2\,nm$ was applied.[14] The van der Waals interactions used a cutoff of $1.2\,nm$ for short-range interactions. PME was used for the long-range electrostatics, and a cutoff of $1.2\,nm$ was used for short-range interactions.[15]
The system coordinates were optimized using a steepest descent algorithm with a convergence criterion of forces smaller than $1000.0\,kJ/mol/nm$, a step size of $0.01$, and a maximum step number of $50000$ steps.[16] All following steps used the leap-frog integrator with a timestep of $0.002\,fs$. Bonds involving hydrogens were constrained with the linear constraint solver for molecular simulations (LINCS), with iteration set to $1$ and order to $4$.[17]

The system equilibration was started under NVT conditions and coupled with the V-rescale thermostat to reach a temperature of $298\ K$.[18] The simulation length was $100\ ps$, and position restraints were applied to the complex. Afterward, an NPT equilibration step was performed including a Berendsen Barostat and a target system pressure of $1\ bar/n$. The complex was restrained with position restraints.[19]
A production run of $100\ ns$ was conducted without any restraints, of which $10\ ns$ were used for equilibration. Every $10\ ps$, a coordinate snapshot was stored, resulting in $10\ 000$ snapshots per run. The entire simulation protocol workflow was repeated three times for the BI-5232 complex, TPP complex, and apo simulations, resulting in a total of $870\ ns$ of production runs.
*Simulation Analysis:*
Analysis was carried out using PyMol and Gromacs tools.[12, 20] Plotting and statistical analysis were performed using Matplotlib, Pandas, and Numpy.[21-23]

##### 1.2. In-Line probing

The aptamer sequence was amplified in a PCR with T7 promotor-containing primers and in-vitro transcribed using T7 RNA polymerase (NEB). After PAGE-Gel purification, ~80 pmol of RNA was dephosphorylated with Shrimp Alkaline Phosphatase (NEB), labelled with [γ-32P]-ATP (Hartmann Analytic) and PAGE-purified. In-line probings were performed as previously described [24, 25]. [γ-32P]-labeled RNA (1 kBq) was incubated with 20 mM MgCl2, 100 mM KCl and 50 mM Tris–HCl (pH 8.3) and the ligand (TPP and TPP-analogs) or H2O for ∼48 h and separated on a denaturing 8 %PAGE-gel. The band pattern was visualized in a phosphorimager (GE Healthcare Life Sciences) after 20-40 h of exposure and analyzed using Quantity One.

##### 1.3. Plasmid construction

The TPPK4-HHR riboswitch sequence was inserted into a psiCHECK-2 vector, encoding Renilla luciferase (hRluc) and firefly luciferase (hluc+) via overhang extension PCR. For FACS experiments a vector encoding for EGFP and mCherry (Figure) was used and the riboswitch was inserted via Gibson Assembly. The construct was introduced into the 3'-UTR of the EGFP gene in the exact same position as in the psiCHECK-2 vector, while mCherry expression served for transfection control. All primers used for PCR were ordered from Sigma-Aldrich and are listed in (Table S1). Plasmids were purified using a Zyppy Plasmid Miniprep kit (Zymo Research).

##### 1.4. Cell culture experiments

Hela cells were cultivated in Dulbecco's Modified Eagle's Medium (Gibco DMEM, Fisher Scientific) supplemented with 10% (v/v) fetal calf serum and 1% (v/v) penicillin/streptomycin (Fisher Scientific) and incubated at 37 °C in a 5% CO2 humidified atmosphere. The day before plasmid transfection, 15000 cells/well in 100 µl medium/well were seeded in a 96-well plate. For transient transfection a Lipofectamine3000 kit (Thermo Fisher Scientific) was used. After incubation of 4 h at 37°C, the medium was replaced by Dulbecco's Modified Eagle's Medium without thiamine-HCl (Gibco DMEM, Fisher Scientific) or Dulbecco's Modified Eagle's Medium without thiamine-HCl (Gibco DMEM, Fisher Scientific), supplemented with the respective ligand (thiamine-HCl or the alternative TPP-aptamer ligands). Twenty-four hours after transfection, luciferase activity or EGFP/mCherry expression was assessed. The luciferase expression of psiCHECK-2 transfected cells was measured using the Dual-Luciferase Reporter Assay System (Promega) according to the manufacturer's protocol. The luminescence was quantified in a Spark multimode microplate reader (Tecan), applying a settle time of 0 ms and an integration time of 2000 ms.

##### 1.5. Flow cytometry

After twenty-four hours of incubation at 37 °C, transfected cells were trypsinated, washed with PBS and resuspended in PBS supplemented with FBS (5% (v/v)), EDTA (1 mM) and HEPES (25 mM). Expression of EGFP and mCherry was analyzed using a Beckman Coulter CytoFLEX™ flow cytometer (blue laser, 488 nm and yellow-green laser, 561 nm) and FlowJo v10 software.

##### 1.6. Surface Plasmon Resonance

SPR experiments were carried out at 25 °C on a BiacoreT200 instrument. Immobilization of a single stranded DNA (5'-CGTCGCAGATCGTGTCTTCC[Am C7) to CM5 chips (Cytiva) was performed as described by Liu et al..[26] RNA was captured via the single stranded oligonucleotide. The utilized RNA (in vitro transcribed and PAGE purified) consists of the natural TPP aptamer sequence from the *thiM* riboswitch [2], a short linker ($U_3$) and a hybridization sequence complementary to the immobilized DNA ( 5´ TAATACGACTCACTATAGGAAGACACGATCTGCGACGTTTGCGACTCGGGGTGCCCTTCTGCGTGAAG GCTGAGAAATACCCGTATCACCTGATCTGGATAATGCCAGCGTAGGGAAGTCGC 3´).

As a control, a TPP aptamer with an inactivating G40C mutation was used. Sequence of M1, M2 and M3 are:

TAATACGACTCACTATAGGAAGACACGATCTGCGACGTTTGCGACTCGGGGTGCCCTTCTGCGTGAA GGCTGAGAAATACCCGTATCACCTG**G**TCTGGATAATGCCAGCGTAGGGAAGTCGC;

TAATACGACTCACTATAGGAAGACACGATCTGCGACGTTTGCGACTCGGGGTGCCCTTCTGCGTGAAG GCTGAGAAATACCCGTATCACCTG**T**T**T**TGGATAATGCCAGC**T**TAGGGAAGTCGC;

TAATACGACTCACTATAGGAAGACACGATCTGCGACGTTTGCGACTCGGGGTGCCCTTCTGCGTGAAG GCTGAGAAATACCCGTATCACCTG**T**TCTGGATAATGCCAGCGTAGGGAAGTCGC.

Binding assays were performed with a flow rate of 30 µL/min in 10 mM HEPES pH 7.4, 100 mM NaCl, 50 mM KCl, 5 mM $MgCl_2$, 0.05 % (v/v) Tween 20, 2 % (v/v) DMSO. Compounds were tested in a single cycle experiment at five different concentrations (e.g. 320 nM, 1.6 µM, 8 µM, 40 µM, 200 µM), with 2 min association and a final dissociation time of 15 min. For regeneration, 50 mM EDTA was used. TPP was used at concentrations of 625 pM, 1,25 nM, 2,5 nM, 5 nM and 10 nM).

Sensograms were evaluated using Biacore T200 Evaluation Software. The responses at the different concentrations were either fitted using a steady state affinity model or direct curve fitting using a 1:1 interaction model to determine the $K_D$ value as well as kinetic parameters ($k_a$ and $k_d$).

#### 2. Supporting Figures

***Complex stability and riboswitch fold stability in MD-simulations***

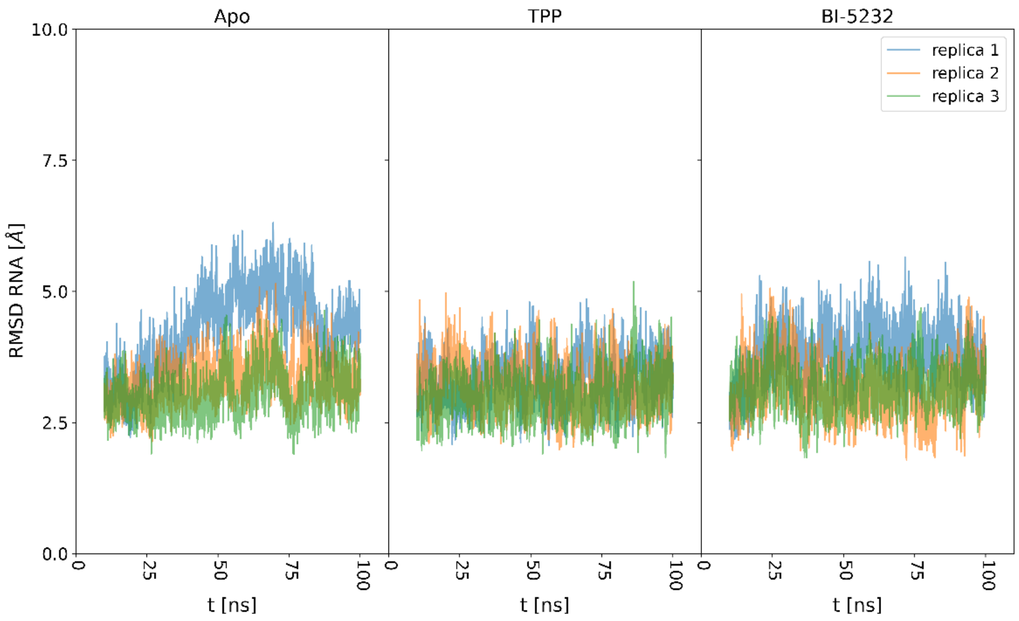


***Supplementary Figure S 1:*** *Stability of the thiM aptamer structures during the simulations. Shown is the root mean square deviation (RMSD) of the RNA atom coordinates over the simulation time compared to the energy minimized initial structure. The low average RMSD value of 3.2 Å for the TPP complex simulations and 3.3 Å for the BI-5232 complex simulations, shows that only minor changes of the overall riboswitch structure occur. In replica 1 of the Apo structure a change in the loop region of residues 25-20 leads to an increase of the RMSD for a approximately 30 ns, leading to an average RMSD of 3.5 Å. However, the binding site is not affected by this change.*

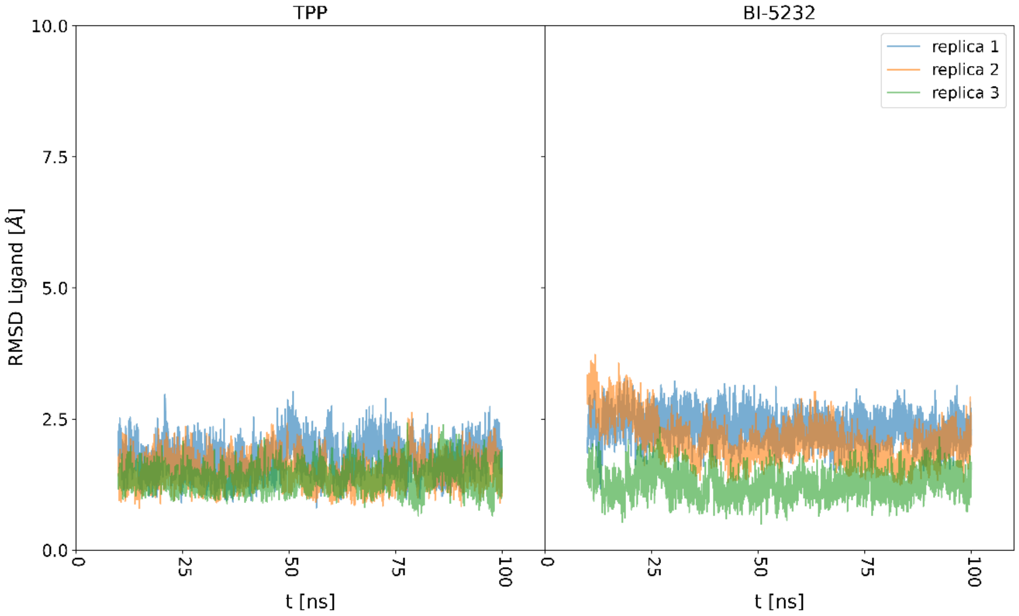


***Supplementary Figure S 2:*** *Stability of the ligand conformations during the simulations of the complexes with the thiM aptamer, compared to the energy minimized initial structures. The positions of TPP and BI-5232 are preserved during the simulations. The slightly increased RMSD for BI-5232 (average: 1.9 Å) in comparison to TPP (average: 1.5 Å) is a consequence of the flexibility of the piperidine ring of BI-5232.*

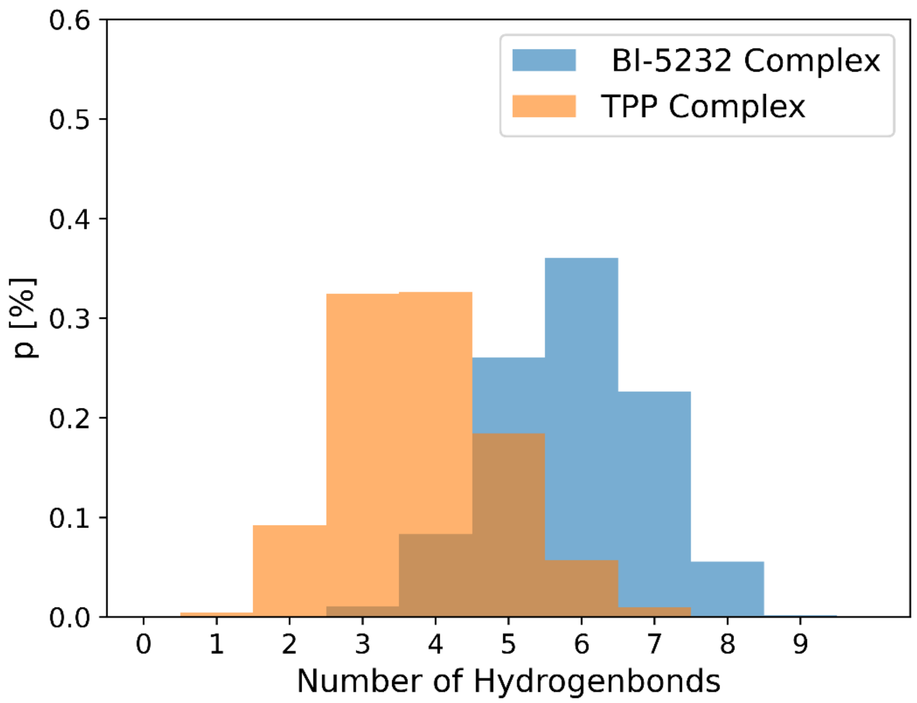


***Supplementary Figure S 3:*** *Hydrogen bond interaction count between the thiM aptamer and TPP and BI-5232, as observed during the simulations. BI-5232 shows on average two additional H-bonds as a result of the insertion of the bicyclic aromatic "tail" into the PP helix (see Fig. 3 in the main text).*

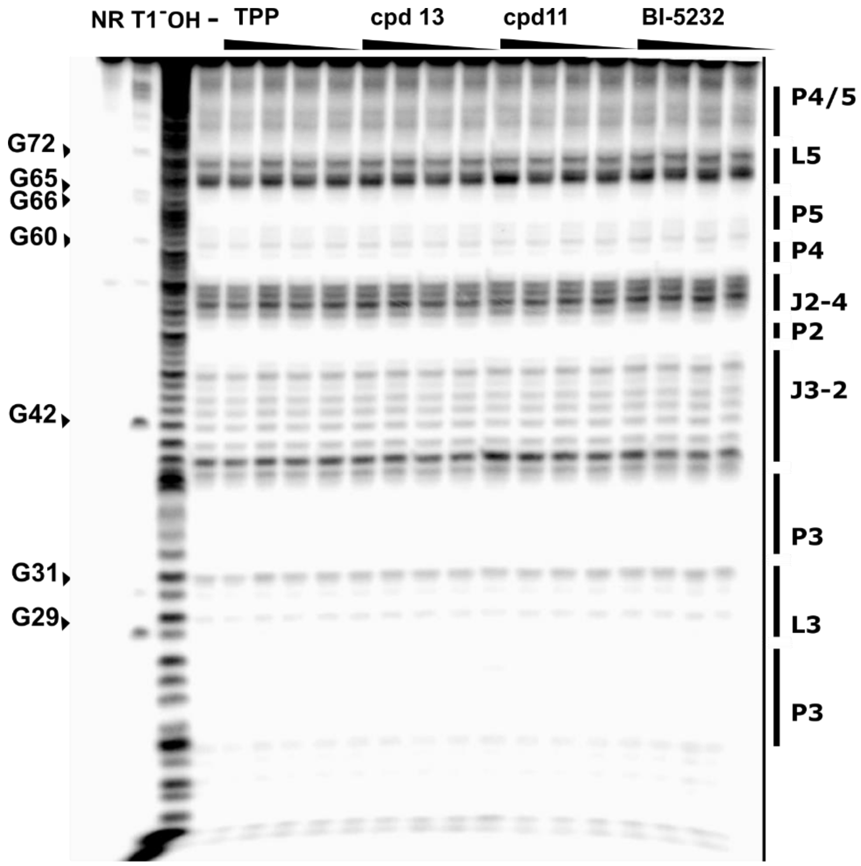


***Supplementary Figure S 4:*** *PAGE-gel analysis of in line probing with G40C mutant (negative control). The 5‘ $^{32}$P-labeled RNA was incubated without ligand (-) or with TPP, 13 11 or BI-5232 1:5 dilutions from 5 nM). 5‘ $^{32}$P-labeled RNA left untreated (precursor, P), treated with T1 RNase (T1) or digested in alkaline condition (OH) serve as controls*

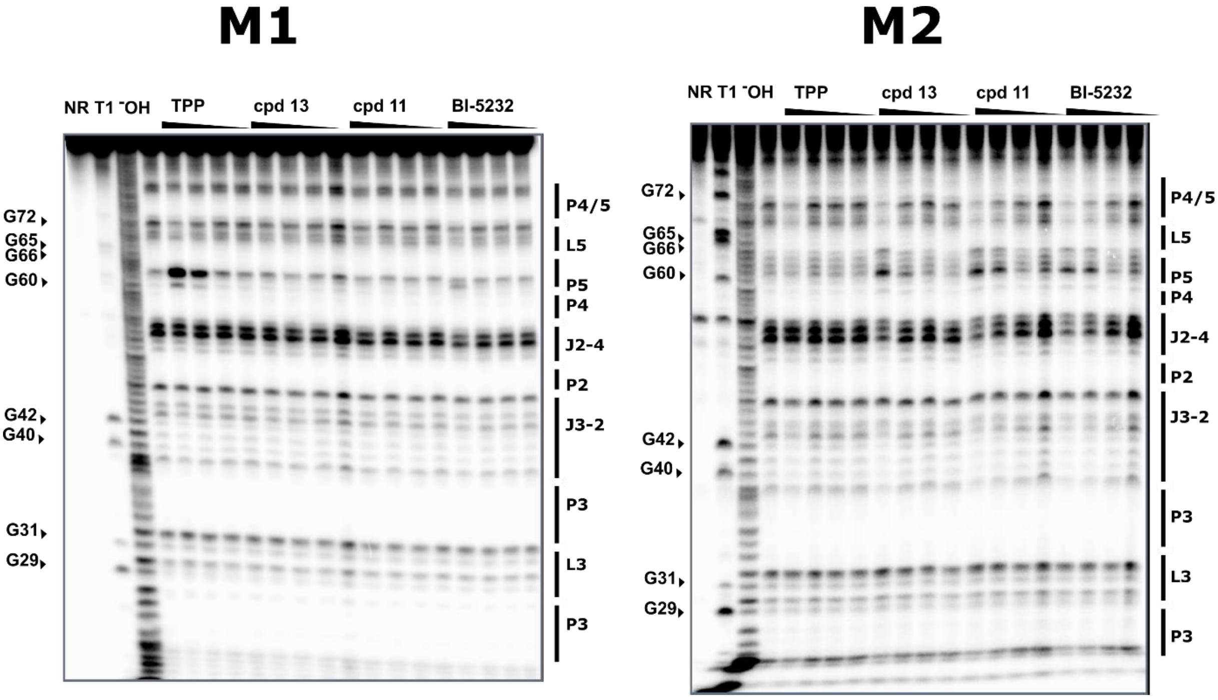


***Supplementary Figure S 5:*** *PAGE gel analysis of in-line probing reactions of thiM mutants 1 (left) and 2 (right) with TPP and compounds BI-5232, 11 and BI-5232.The 5‘ $^{32}$P-labeled thiM aptamer was incubated without ligand (-) or with TPP, BI-5232, 11 or BI-5232 (1:5 dilutions from 5 nM). 5‘ $^{32}$P-labeled RNA left untreated (precursor, P), treated with T1 RNase (T1) and alkaline digestion (OH) serve as controls.*

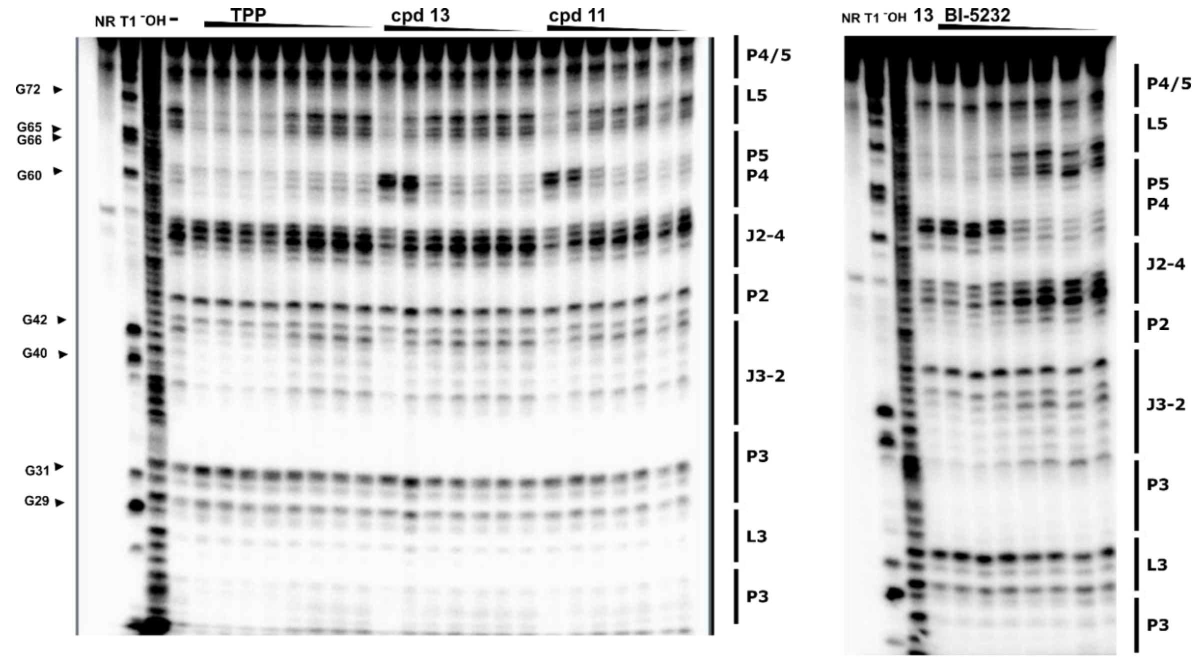


***Supplementary Figure S 6:*** *PAGE gel analysis of in-line probing reactions of thiM WT aptamer with TPP and the analogs BI-5232, 11 and BI-5232. The 5‘ $^{32}$P-labeled thiM aptamer was incubated without ligand (-) or with TPP, BI-5232, 11 or 12 (TPP: 10 µM and 1:5 dilutions from 5 µM, analogs: 1:5 dilutions from 5 µM). 5‘ $^{32}$P-labeled RNA left untreated (precursor, P), treated with T1 RNase (T1) or digested in alkaline condition (OH) serve as controls*

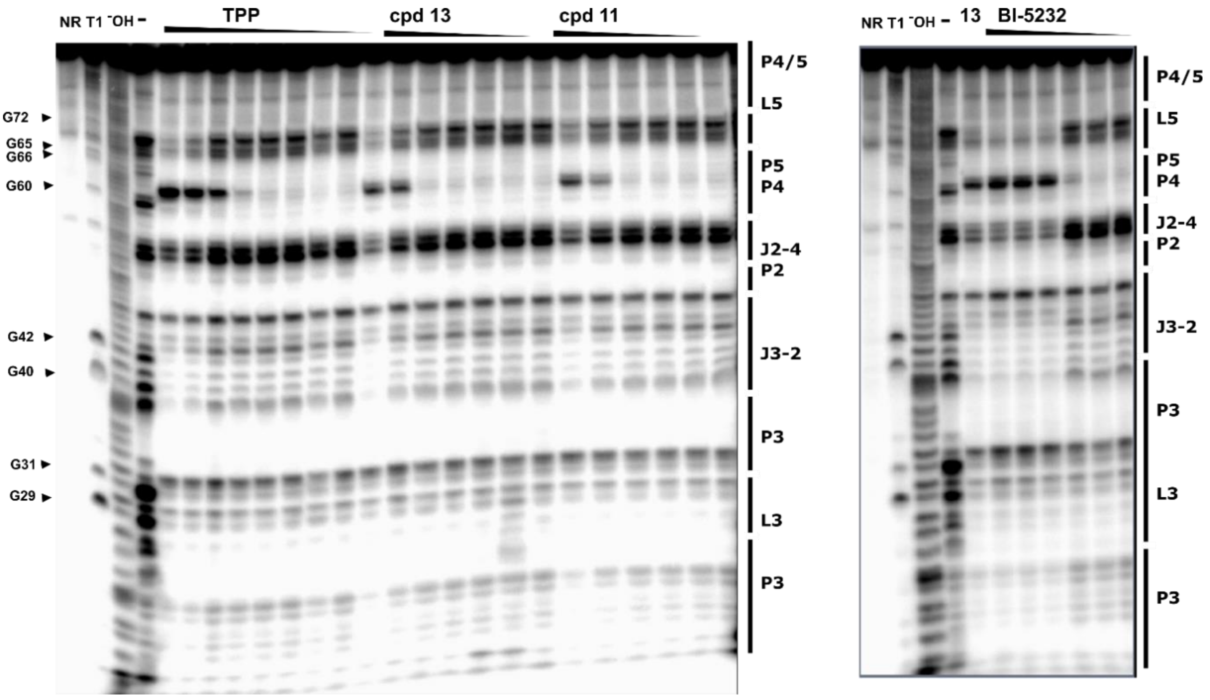


***Supplementary Figure S 7:*** *PAGE gel analysis of in-line probing reaction of the mutant aptamer M3 with TPP and the analogs BI-5232, 11 and BI-5232. The 5‘ $^{32}$P-labeled RNA was incubated without ligand (-) or with TPP, BI-5232, 11 or BI-5232 (TPP: 10 µM and 1:5 dilutions from 5 µM, analogs: 1:5 dilutions from 5 µM). 5‘ $^{32}$P-labeled RNA left untreated (precursor, P), treated with T1 RNase (T1) or digested in alkaline condition (OH) serve as controls*

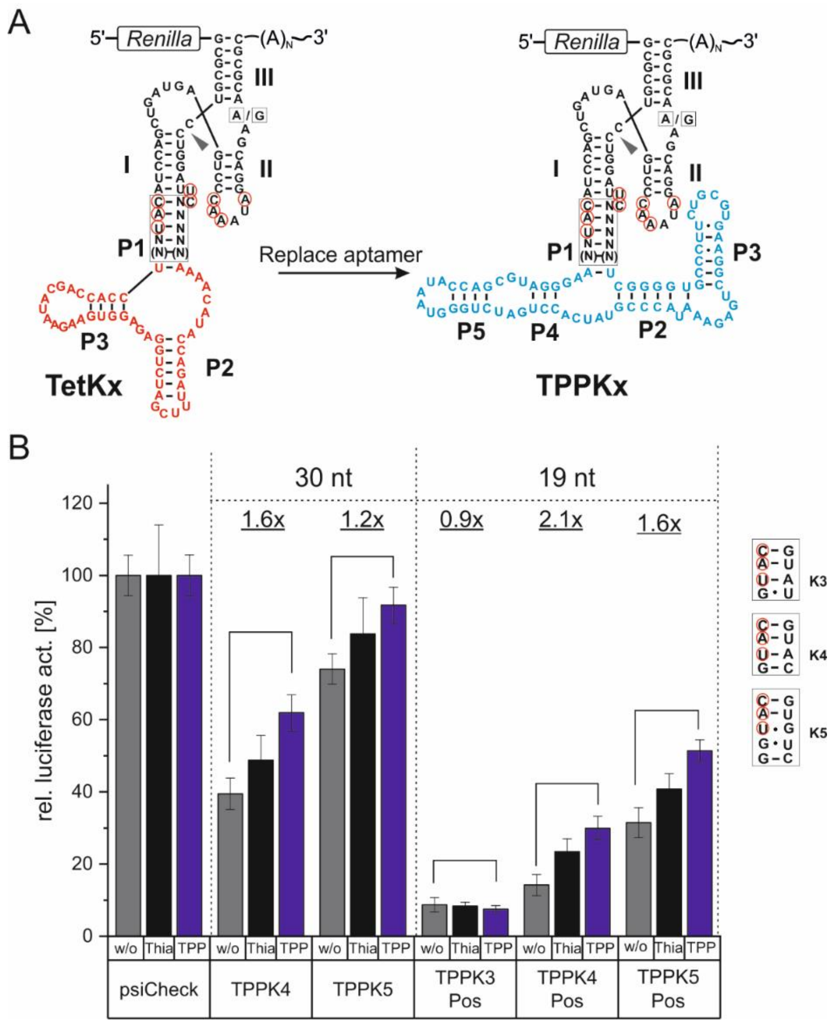


***Supplementary Figure S 8:*** *Generation of TPP-responsive ON-switching aptazymes on the basis of the TetKx-platform. (A) Secondary structure of TPPKx aptazymes derived from the TetKx-platform through replacement of the tetracycline aptamer by the TPP aptamer from the thiM motif. (B) Luciferase activity measurement of constructs with TPPKx aptazymes inserted 30 nt or 19 nt downstream of the stop codon of a Renilla luciferase. The communication modules of TPPKx aptazymes are depicted on the right. Transfected HeLa cells were incubated in the absence (grey, thiamine-depleted medium) or presence of 500 µM thiamine (black) or 500 µM TPP (blue). A construct without any regulatory element serves as control and was set to 100 %. Numbers above the bars indicate the dynamic range of regulation. Shown is mean ± SD, n=3 replicates.*

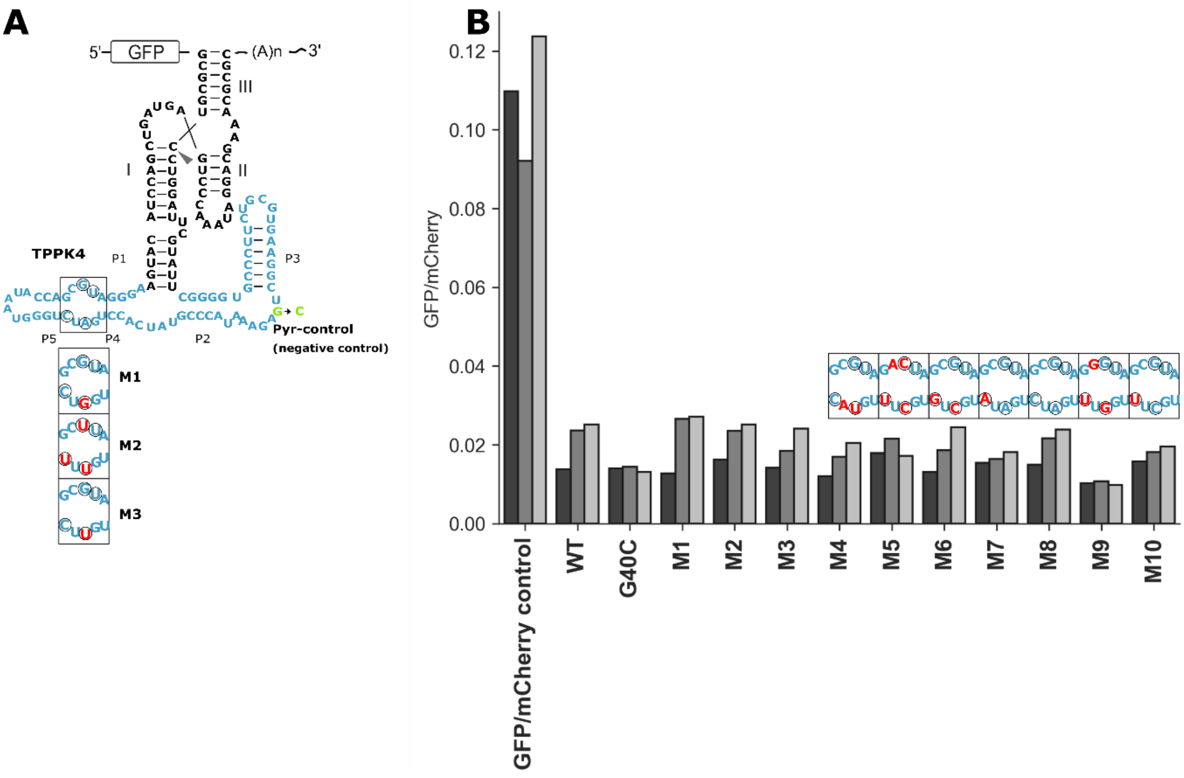


***Supplementary Figure S 9:*** *Screening of TPPKx aptazymes with mutated PP-helix. Relative EGFP fluorescence of constructs containing the aptazyme 19 nt downstream of an EGFP gene. (A) Secondary structure of the TPP-responsive ON-switching aptazyme. Nucleotides A61, C63, C77, G78 were randomized. (B) HeLa cells were transfected with different mutants, incubated in the absence (black, thiamine-depleted medium) or presence of 500 µM thiamine (grey) or 500 µM compound 11light grey). The fluorescence was measured via flow cytometry 24 h after transfection. Pos. control: vector without aptazyme, G40C: aptazyme with mutated Pyr-binding pocket. Mutants 1-10 (M1-M10) were sequenced to determine the effects of individual mutations. M4, M5 and M9 contain mutations that are not part of the randomized sequence; M8 is identical to the original TPP WT sequence that was used as a template.*

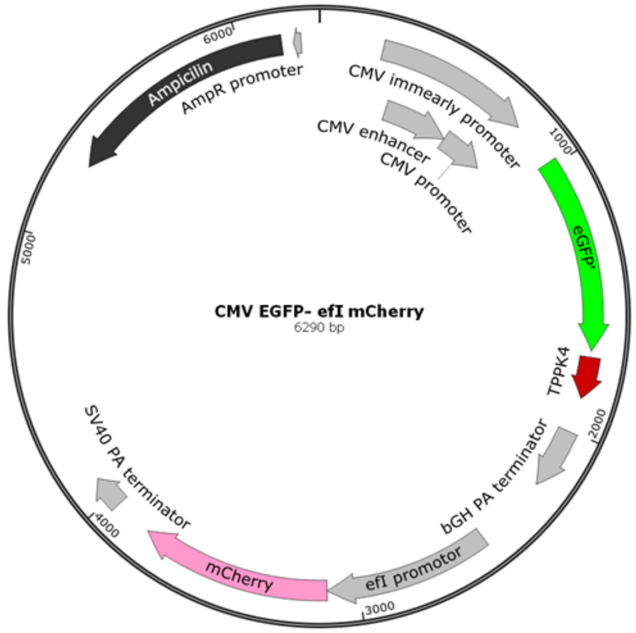


***Supplementary Figure S 10****: Plasmid map of EGFP-mCherry vector used to test TPPKx aptazymes in combination with the investigated compounds.*

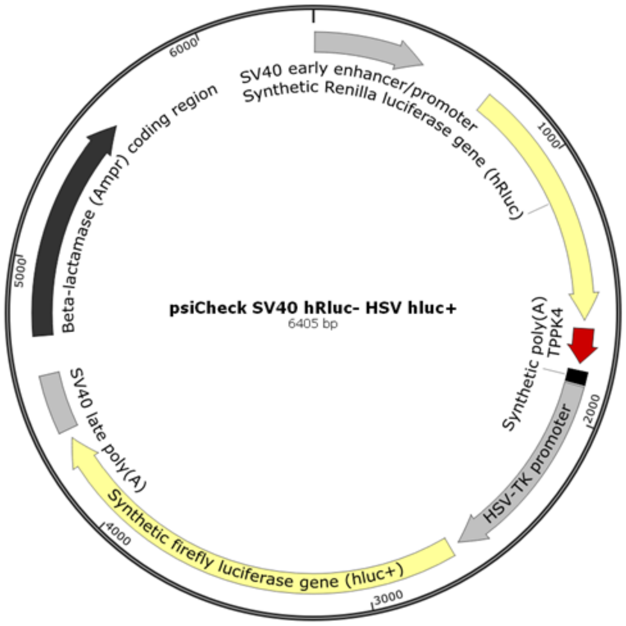


***Supplementary Figure S 11:*** *Plasmid map of luciferase expressing vector used to determine activity of TPPKx aptazymes.*

#### 3. Supplementary Tables

*Supplementary Table S 1:* ***Primer sequences for TPP riboswitch insertion in the 3'-UTR of reporter genes.*** *Aptamer sequences are highlighted in green, communication modules in yellow. Nucleotides that were randomized to screen for analog selectivity are highlighted in pink*

| | |
|---|---|
| WT_On fwd | [Phos]CCGTATCACCTGATCTGGATAATGCCAGCGTAGGGAAGTACATCCAGCTG ATGAGTCCCAAATA |
| WT_On rev | GTATTTCTCAGCCTTCACGCAGAAGGGCACCCCGAGTACGAATCCAGGACGCGC |
| Screen_fwd | [Phos]CCGTATCACCTNATCTGGATAATGCCAGNNTAGGGAAGTACATCCAGCTGA TGAGTCCCAAATA |
| Screen_rev | GTATTTCTCAGCCTTCACGCAGAAGGGCACCCCGAGTACGAATCCAGGACGCGC |
| WT_Off fwd | [Phos]CCCGTATCACCTGATCTGGATAATGCCAGCGTAGGGAACCATCCTGGATT CCACTGCTATCCACA |
| WT_Off rev | TATTTCTCAGCCTTCACGCAGAAGGGCACCCCGACCATTTCGTCCTATTTGGGAC TCGTCAG |
| Mutant 3 Off fwd | [Phos]CCCGTATCACCTGTTCTGGATAATGCCAGCGTAGGGAACCATCCTGGATTC CACTGCTATCCACA |
| Mutant 3 Off rev | TATTTCTCAGCCTTCACGCAGAAGGGCACCCCGACCATTTCGTCCTATTTGGGAC TCGTCAG |

*Supplementary Table S 2:* ***SPR data of thiM aptamer and aptamer mutants***

| Compound ID | Aptamer | ka [1/Ms] ± SD (x $10^6$) | kd [1/s] ± SD (x $10^{-2}$) | $K_D$ [nmol/l] ± SD |
|---|---|---|---|---|
| 13 | WT | 7.77 ± 0.71 | 6.75 ± 1.00 | 8.8 (4) ± 1.5 |
| | M1 | 9.65 ± 1.58 | 10.30 ± 1.59 | 10.8 (3) ± 1.1 |
| | M2 | 6.49 ± 0.51 | 1.15 ± 0.18 | 1.8 (3) ±0.04 |
| | M3 | | | 34.1 (3) ± 0.4 |
| 11 | WT | 3.36 ± 0.22 | 5.49 ± 0.40 | 16.4 (4) ± 0.6 |
| | M1 | 4.12 ± 0.71 | 17.12 ± 5.00 | 40.9 (3) ± 4.1 |
| | M2 | 5.88 ± 0.24 | 0.37 ± 0.20 | 0.6 (6) ± 0.2 |
| | M3 | 7.28 ± 4.00 | 35.72 ± 18.00 | 49.7 (4) ± 3.9 |
| BI-5232 | WT | 3.49 ± 1.69 | 0.34 ± 0.12 | 1.0 (5) ± 0.2 |
| | M1 | 2.62 ± 1.08 | 2.43 ± 0.59 | 9.8 (5) ± 1.2 |
| | M2 | 2.16 ± 0.96 | 0.39 ± 0.14 | 2.0 (5) ± 0.5 |
| | M3 | 2.47 ± 0.35 | 0.61 ± 0.08 | 2.5 (4) ± 0.3 |
| TPP (S1) | WT | 0.47 ± 0.02 | 0.03 ± 0.001 | 0.7 (2) ± 0.1 |
| | M1 | 0.39 ± 0.01 | 9.23 ± 0.20 | 232.9 (2) ± 10.9 |
| | M2 | 0.73 ± 0.28 | 44.32 ± 4.00 | 687 (2) ± 210.5 |
| | M3 | | | 2200.5 (2) ± 158.5 |

***Supplementary Table S 3:* SPR binding constants ($K_D$) of TPP and derivatives for the thiM aptamer (WT)***. *The loss in binding affinity of compounds S4 and S5 versus compound S3 underlines the importance of the positive charge of the thiazolium ring. The reduced affinities of compounds S2, S3 compared to TPP (S1) show the strong contribution of the pyrophosphate moiety to binding.*

| Compound ID | Structure | $K_D$ [nM] |
|---|---|---|
| **S1(TPP)** | 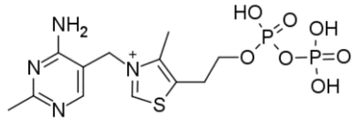 | 0.7 |
| **S2** | 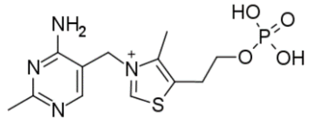 | 118 |
| **S3** | 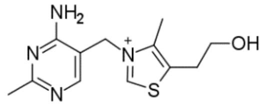 | 2794 |
| **S4** | 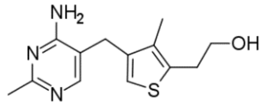 | >200000 |
| **S5** | 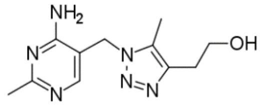 | 59960 |

### 4. Compound Synthesis

##### General Methods and Materials

All commercially available chemicals were used as received with purity as reported from the supplier. Anhydrous solvents were either purchased or prepared according to standard procedures[S1] and stored over molecular sieves under argon. Unless stated otherwise, all reactions were carried out in oven-dried (at 120 °C) glassware under an inert atmosphere of argon. A Biotage Initiator Classic microwave reactor was used for reactions conducted in a microwave oven. Reactions were monitored by TLC on aluminum-backed plates coated with Merck Kieselgel 60 F 254 with visualization under UV light at 254 nm, and with HPLC-MS analysis (for HPLC-MS methods, *see below.*). Unless stated otherwise, crude products were purified by flash column chromatography on silica (using a Biotage IsoleraOne, Biotage IsoleraFour or CombiFlash® Teledyne Isco system) or by (semi)-preparative reversed-phase HPLC (Agilent Acquity or Waters instruments). Unless specified otherwise, the purity of all final compounds was determined to be ≥ 95% by LC-MS. Nuclear magnetic resonance (NMR) spectra were recorded at room temperature ((22 ± 1 °C), on a Bruker Avance 400 spectrometer with tetramethylsilane as an internal reference. Chemical shifts δ are reported in parts per million (ppm). $^{1}$H NMR spectra were referenced to the residual partially non-deuterated solvent signal of DMSO (δ = 2.50 ppm). Coupling constants *J* are reported in Hz, and splitting patterns are described as br = broad, s = singlet, d = doublet, t = triplet, q = quartet, quin = quintet and m = multiplet. MarvinSketch software version 20.19.1 was used to generate compound names.

##### LC-MS Methods

**Method 1:**

| | | | | | | |
|---|---|---|---|---|---|---|
| Instrument: | **Agilent 1200 & 1956A** | | | | | |
| Column: | **Xbridge Shield RP18 2.1*50mm,5um** | | | | | |
| Column temperature: | **40℃** | | | | | |
| Mobile phase A(MPA) | **H2O+10mM NH4HCO3** | | | | | |
| Mobile phase B(MPB) | **Acetonitrile** | | | | | |
| Flow rate: | **0.8 mL/min (0.00-2.50min) 1.0 mL/min (2.51-3.00min)** | | | | | |
| Gradient Ratio: | **Time(min)** | **0.00** | **0.30** | **2.10** | **2.48** | **2.50** |
| | **MPA(%)** | **100** | **100** | **40** | **40** | **100** |
| | **MPB(%)** | **0** | **0** | **60** | **60** | **0** |
| Detection: | **DAD** | | | | | |
| MS Mode: | **Positive** | | | | | |
| MS Range: | **100-1000** | | | | | |

**Method 2:**

| | | | | | | |
|---|---|---|---|---|---|---|
| Instrument: | Agilent 1200 & 1956A | | | | | |
| Column: | Xbridge Shield RP18 2.1*50mm,5um | | | | | |
| Column temperature: | 40℃ | | | | | |
| Mobile phase A(MPA) | H2O+10mM NH4HCO3 | | | | | |
| Mobile phase B(MPB) | Acetonitrile | | | | | |
| Flow rate: | 0.8 mL/min (0.00-2.50min) 1.0 mL/min (2.51-3.00min) | | | | | |
| Gradient Ratio: | Time(min) | 0.00 | 0.30 | 2.10 | 2.48 | 2.50 |
| | MPA(%) | 100 | 100 | 40 | 40 | 100 |
| | MPB(%) | 0 | 0 | 60 | 60 | 0 |
| Detection: | DAD | | | | | |
| MS Mode: | Negative | | | | | |
| MS Range: | 100-1000 | | | | | |

**Method 3:**

| | | | | | | |
|---|---|---|---|---|---|---|
| Instrument: | Agilent 1200 & G6120B | | | | | |
| Column: | Kinetex C18 30*2.1mm,5μm | | | | | |
| Column temperature: | 40℃ | | | | | |
| Mobile phase A(MPA) | H2O+0.04 %(v/v) TFA | | | | | |
| Mobile phase B(MPB) | ACN+0.02 %(v/v) TFA | | | | | |
| Flow rate: | 1.0 mL/min(0-1.80min),1.2ml/min(1.81-2.00) | | | | | |
| Gradient Ratio: | Time(min) | 0.00 | 1.00 | 1.80 | 1.81 | 2.00 |
| | MPA(%) | 95 | 5 | 0 | 95 | 95 |
| | MPB(%) | 5 | 95 | 100 | 5 | 5 |
| Detection: | 220 nm 254 nm | | | | | |
| MS Mode: | Positive | | | | | |
| MS Range: | 100-1000 | | | | | |

**Method 4:**

| | | | | | | |
|---|---|---|---|---|---|---|
| Instrument: | Agilent 1200 & 1956A | | | | | |
| Column: | Xbridge Shield RP18 2.1*50mm,5um | | | | | |
| Column temperature: | 40℃ | | | | | |
| Mobile phase A(MPA) | H2O+10mM NH4HCO3 | | | | | |
| Mobile phase B(MPB) | Acetonitrile | | | | | |
| Flow rate: | 1.0 mL/min (0.00-2.48min) 1.2 mL/min (2.50-3.00min) | | | | | |
| Gradient Ratio: | Time(min) | 0.00 | 2.00 | 2.48 | 2.50 | 3.00 |
| | MPA(%) | 90 | 20 | 20 | 90 | 90 |
| | MPB(%) | 10 | 80 | 80 | 10 | 10 |
| Detection: | 220 nm | | | | | |
| MS Mode: | Positive | | | | | |
| MS Range: | 100-1000 | | | | | |

**Method 5:**

| | | | | | | |
|---|---|---|---|---|---|---|
| Instrument: | Agilent 1200 & 610MS | | | | | |
| Column: | Luna C18, 2.0*50mm 5um | | | | | |
| Column temperature: | 40℃ | | | | | |
| Mobile phase A(MPA) | H2O + 0.0375%TFA | | | | | |
| Mobile phase B(MPB) | Acetonitrile + 018% TFA | | | | | |
| Flow rate: | 0.8 mL/min (0.00-2.50min) 1.0 mL/min (2.51-3.00min) | | | | | |
| Gradient Ratio: | Time(min) | 0.00 | 0.40 | 3.40 | 3.85 | 8.86 |
| | MPA(%) | 99 | 99 | 0 | 0 | 99 |
| | MPB(%) | 1 | 1 | 100 | 100 | 1 |
| Detection: | DAD | | | | | |
| MS Mode: | Positive | | | | | |
| MS Range: | 100-1000 | | | | | |

**Method 6:**

| | | | | | | |
|---|---|---|---|---|---|---|
| Instrument: | SHIMADZU M20A& MS2010EV | | | | | |
| Column: | Venusil XBP-C18 , 2.1x50mm, 5μm | | | | | |
| Column temperature: | 50℃ | | | | | |
| Mobile phase A(MPA) | H2O+0.04 %(v/v) TFA | | | | | |
| Mobile phase B(MPB) | ACN+0.02 %(v/v) TFA | | | | | |
| Flow rate: | 1.0 mL/min | | | | | |
| Gradient Ratio: | Time(min) | 0.00 | 2.20 | 2.48 | 2.50 | 3.00 |
| | MPA(%) | 90 | 10 | 10 | 90 | 90 |
| | MPB(%) | 10 | 90 | 90 | 10 | 10 |
| Detection: | 220 nm 254nm | | | | | |
| MS Mode: | Positive | | | | | |
| MS Range: | 100-1000 | | | | | |

**Method 7:**

| | | | | | | |
|---|---|---|---|---|---|---|
| Instrument: | Shimadzu LC-20AD&MS 2010 | | | | | |
| Column: | Luna-C18 2.0x30mm, 3μM | | | | | |
| Column temperature: | 40℃ | | | | | |
| Mobile phase A(MPA) | H2O+0.04 %(v/v) TFA | | | | | |
| Mobile phase B(MPB) | ACN+0.02 %(v/v) TFA | | | | | |
| Flow rate: | 0.8 mL/min(0.01-1.51min,1.2mL/min(1.52-2.00min) | | | | | |
| Gradient Ratio: | Time(min) | 0.01 | 1.15 | 1.65 | 1.66 | 2.00 |
| | MPA(%) | 90 | 10 | 10 | 90 | 90 |
| | MPB(%) | 10 | 90 | 90 | 10 | 10 |
| Detection: | 220 nm 254nm | | | | | |
| MS Mode: | Positive | | | | | |
| MS Range: | 100-1000 | | | | | |

**Method 8:**

| | |
|---|---|
| Instrument: | Shimadzu LC-30AD&MS 2020 |
| Column: | Kinetex EVO C18 30*2.1mm, 5um |
| Column temperature: | 40℃ |
| Mobile phase A(MPA) | H2O+0.04 %(v/v) TFA |
| Mobile phase B(MPB) | ACN+0.02 %(v/v) TFA |
| Flow rate: | 1.5 mL/min (0.00-1.50min) |

| Gradient Ratio: | | | | | | |
|---|---|---|---|---|---|---|
| | Time(min) | 0.01 | 0.80 | 1.20 | 1.21 | 1.50 |
| | MPA(%) | 95 | 5 | 5 | 95 | 95 |
| | MPB(%) | 5 | 95 | 95 | 5 | 5 |

| | |
|---|---|
| Detection: | 220 nm 254nm |
| MS Mode: | Positive |
| MS Range: | 100-1000 |

**Method 9:**

| | | | | | |
|---|---|---|---|---|---|
| Device description: | Waters Acquity, QDa Detector | | | | |
| Column: | XBridge C18_3.0 x 30 mm_2.5 µm | | | | |
| Column producer: | Waters | | | | |
| Description: | | | | | |
| Gradient/Solvent Time [min] | % Sol [Water 0.1% NH3] | % Sol [Acetonitrile] | Flow [ml/min] | Temp [°C] | Back pressure [PSI] |
| 0.0 | 95.0 | 5.0 | 1.5 | 60.0 | |
| 1.3 | 0.0 | 100.0 | 1.5 | 60.0 | |
| 1.5 | 0.0 | 100.0 | 1.5 | 60.0 | |
| 1.6 | 95.0 | 5.0 | 1.5 | 60.0 | |

**Method 10:**

| | | | | | |
|---|---|---|---|---|---|
| Device description: | Agilent 1200 with DA- and MS-Detector | | | | |
| Column: | XBridge C18_3.0 x 30 mm_2.5 µm | | | | |
| Column producer: | Waters | | | | |
| Description: | | | | | |
| Gradient/Solvent Time [min] | % Sol [Water 0.1% NH3] | % Sol [Acetonitrile] | Flow [ml/min] | Temp [°C] | Back pressure [PSI] |
| 0.0 | 97.0 | 3.0 | 2.2 | 60.0 | |
| 0.2 | 97.0 | 3.0 | 2.2 | 60.0 | |
| 1.2 | 0.0 | 100.0 | 2.2 | 60.0 | |
| 1.25 | 0.0 | 100.0 | 3.0 | 60.0 | |
| 1.4 | 0.0 | 100.0 | 3.0 | 60.0 | |

**Method 11:**

| | | | | | |
|---|---|---|---|---|---|
| Device description: | Waters Acquity, QDa Detector | | | | |
| Column: | XBridge C18_3.0 x 30 mm_2.5 µm | | | | |
| Column producer: | Waters | | | | |
| Description: | | | | | |
| Gradient/Solvent Time [min] | % Sol [Water 0.1% NH3] | % Sol [Acetonitrile] | Flow [ml/min] | Temp [°C] | Back pressure [PSI] |
| 0.0 | 95.0 | 5.0 | 1.5 | 60.0 | |
| 1.3 | 0.0 | 100.0 | 1.5 | 60.0 | |
| 1.5 | 0.0 | 100.0 | 1.5 | 60.0 | |
| 1.6 | 95.0 | 5.0 | 1.5 | 60.0 | |

**Method 12:**

| | | | | | |
|---|---|---|---|---|---|
| Device description: | Agilent 1200 with DA- and MS-Detector | | | | |
| Column: | Sunfire C18_3.0 x 30 mm_2.5 µm | | | | |
| Column producer: | Waters | | | | |
| Description: | | | | | |
| Gradient/Solvent Time [min] | % Sol [Water 0.1% TFA (v/v)] | % Sol [Acetonitrile] | Flow [ml/min] | Temp [°C] | Back pressure [PSI] |
| 0.0 | 97.0 | 3.0 | 2.2 | 60.0 | |
| 0.2 | 97.0 | 3.0 | 2.2 | 60.0 | |
| 1.2 | 0.0 | 100.0 | 2.2 | 60.0 | |
| 1.25 | 0.0 | 100.0 | 3.0 | 60.0 | |
| 1.4 | 0.0 | 100.0 | 3.0 | 60.0 | |

**Method 13**

| | | | | | |
|---|---|---|---|---|---|
| Device description: | Waters Acquity, QDa Detector | | | | |
| Column: | Sunfire C18_3.0 x 30 mm_2.5 µm | | | | |
| Column producer: | Waters | | | | |
| Description: | | | | | |
| Gradient/Solvent Time [min] | % Sol [Water 0.1% TFA (v/v)] | % Sol [Acetonitrile 0.08% TFA (v/v)] | Flow [ml/min] | Temp [°C] | Back pressure [PSI] |
| 0.0 | 95.0 | 5.0 | 1.5 | 60.0 | |
| 1.3 | 0.0 | 100.0 | 1.5 | 60.0 | |
| 1.5 | 0.0 | 100.0 | 1.5 | 60.0 | |
| 1.6 | 95.0 | 5.0 | 1.5 | 60.0 | |

**Mobile phase preparations**
Examples:

- The mobile phase "Water 0.1% TFA (v/v)" is prepared by adding 1 ml of a commercially available TFA solution to 999 ml water.
- The mobile phase "Water 0.1% $NH_3$" is prepared by adding 4 ml of a commercially available concentrated ammonium hydroxide solution (25 wt%) to 996 ml water.

**Syntax for column description**
Description_Dimensions ID x length_Particle Size
**Convention:** Sections separated by underscores; blanks between numbers and unit; ID and length in mm, particle size in µm; ID and particle size always with one digit
e.g.: XBridge C18_4.6 x 50 mm_3.5 µm

**Chiral SFC method 1**

| | | | | | |
|---|---|---|---|---|---|
| Device description: | Agilent 1260 SFC with DAD and MS | | | | |
| Column: | Chiralpak® IG_4.6 x 250 mm_5 µm | | | | |
| Column producer: | Daicel | | | | |
| Description: | | | | | |
| Gradient/Solvent Time [min] | % Sol [scCO2] | % Sol [ETOH 20mM NH3] | Flow [ml/min] | Temp [°C] | Back pressure [PSI] |
| 0.0 | 60.0 | 40.0 | 4.0 | 40.0 | 2175.0 |
| 10.0 | 60.0 | 40.0 | 4.0 | 40.0 | 2175.0 |

#### Synthesis of compounds

##### 2-methyl-5-[(methylamino)methyl]pyrimidin-4-amine (1)

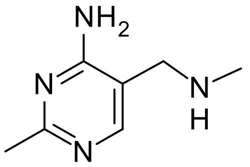

**1A** (61 mg, 0.45 mmol) and methylamine (0.22 mL 2 mol/l in THF, 0.45 mmol) were dissolved in acetic acid (0.02 mL, 0.45mmol) and DCM (2 mL) and stirred 1h at rt. NaBH(OAc)3 (0.15 g, 0.67mmol) was added and the mixture was stirred at rt o/n. The mixture was evaporated, the residue was dissolved in DMF and was purified by rp-chromatography (XBridge, gradient of ACN/water/NH3) and lyophilized to obtain the compound **1** (18.5 mg, 27 % yield)

LC-MS (*Method 10*): $^{t}R$ = 0.32 min, $[M+H]^+$ 152
$^{1}H$ NMR (400 MHz, DMSO-$d_6$) δ ppm 7.85 (s, 1 H) 6.68 (br s, 2 H) 3.47 (s, 2 H) 2.28 (s, 3 H) 2.22 (s, 3 H)

##### 4-amino-2-methylpyrimidine-5-carbaldehyde (1A)

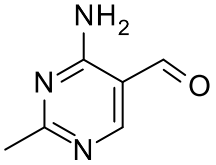

(4-amino-2-methyl-5-pyrimidinyl)methanol (0.5 g, 3.59 mmol) was dissolved in DCM (30 mL). Activated manganese dioxide (2186.6 mg, 25.15 mmol) was added and the mixture was stirred at rt o/n. The mixture was filtrated over Celite, the filter cake was washed with DCM+MeOH and the filtrate was evaporated to obtain compound **1A** (0.47 g, 95% yield)
LC-MS (*method 10*): $^{t}R$ = 0.23 min, $[M+H]^+$ 138

##### methyl[(quinoxalin-6-yl)methyl]amine (2)

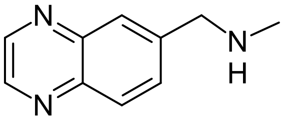

6-Bromomethyl-quinoxaline (80 mg, 0.36 mmol) was dissolved in EtOH (2 mL). Methylamine (0.54 mL 2 mol/L in THF, 1.08 mmol) was added and the mixture was stirred 1h at 60 °C. The solution was purified by reverse phase-chromatography (XBridge, gradient of ACN/water/NH3). Corresponding fractions were lyophilized to obtain compound **2** (14.7 mg, 24 % yield)

LC-MS (*method 10*): $^{t}R$ = 0.59 min, $[M+H]^+$ 174
$^{1}H$ NMR (400 MHz, DMSO-$d_6$) δ ppm 8.91 (dd, *J*=8.36, 1.90 Hz, 2 H) 8.04 (d, *J*=8.62 Hz, 1 H) 8.00 (d, *J*=0.89 Hz, 1 H) 7.84 (dd, *J*=8.62, 1.90 Hz, 1 H) 3.90 (s, 2 H) 2.32 (s, 3 H)

##### 6-[(pyrrolidin-1-yl)methyl]quinoxaline (3)

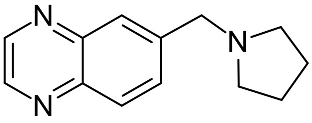

Quinoxaline-6-carbaldehyde (66.7 mg, 0.42 mmol) and pyrrolidine (30 mg, 0.42 mmol) were dissolved in THF (3 mL) and the mixture was acidified with acidic acid to pH 5. The mixture was stirred 0.5 h at rt. NaBH(OAc)3 (111.75 mg, 0.52 mmol) was added and the mixture was stirred 3 days at rt. The mixture was evaporated, the residue was dissolved in MeOH, filtered and the clear solution was purified by rp-chromatography (XBridge, gradient of ACN/water/NH3). Corresponding fractions were lyophilized to obtain the compound **3** (27.2 mg, 30 % yield)

LC-MS (*method 10*): $^{t}R$ = 0.76 min, $[M+H]^+$ 214
$^{1}H$ NMR (400 MHz, DMSO-$d_6$) δ ppm 8.90 - 8.93 (m, 2 H) 8.05 (d, *J*=8.62 Hz, 1 H) 7.98 (d, *J*=1.01 Hz, 1 H) 7.82 - 7.87 (m, 1 H) 3.83 (s, 2 H) 1.73 (spt, *J*=3.27 Hz, 4 H)
4 H: 2 x -$CH_2$- from the Pyrrolidine are in the DMSO peak (predicted chemical shift in Marvin: 2.60 ppm)

**6-{[(3R)-3-fluoropyrrolidin-1-yl]methyl}quinoxaline(4)**

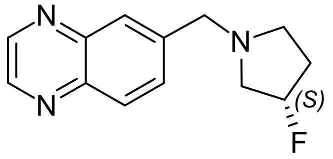

quinoxaline-6-carbaldehyde (63.0 mg, 0.40 mmol) and (3S)-3-fluoropyrrolidine hydrochloride (50 mg, 0.40 mmol) were dissolved in THF (3 mL) and the mixture was acidified with acidic acid to pH 5. The mixture was stirred 0.5 h at rt. NaBH(OAc)3 (105.5 mg, 0.50 mmol) was added and the mixture was stirred at rt o/n. The mixture was evaporated, the residue was dissolved in MeOH, filtered and the clear solution was purified by rp-chromatography (XBridge, gradient of ACN/water/NH3). Corresponding fractions were lyophilized to obtain the compound **4** (12.2 mg, 13 % yield)

LC-MS (*method 10*): $^{t}R$ = 0.71 min, $[M+H]^+$ 232
$^1H$ NMR (400 MHz, DMSO-$d_6$) δ ppm 8.91 - 8.94 (m, 2 H) 8.04 - 8.08 (m, 1 H) 8.00 (d, *J*=1.01 Hz, 1 H) 7.85 (dd, *J*=8.62, 1.90 Hz, 1 H) 5.13 - 5.32 (m, 1 H) 3.88 (s, 2 H) 2.78 - 2.91 (m, 2 H) 2.62 - 2.76 (m, 1 H) 2.37 - 2.45 (m, 1 H) 2.09 - 2.26 (m, 1 H) 1.82 - 1.99 (m, 1 H)

**6-[(3,3-difluoropyrrolidin-1-yl)methyl]quinoxaline (5)**

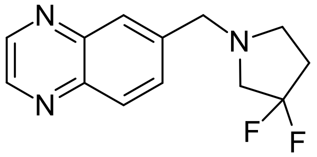

3,3-difluoropyrrolidine hydrochloride (55 mg, 0.38 mmol) was dissolved in DCM (3 ml). Quinoxaline-6-carbaldehyde (60.6 mg, 0.38 mmol) and titanium (iv) isopropoxide (0.11 ml, 0.38 mmol) and the mixture was stirred 5h at rt. Sodium triacetoxyborhydride (243.6 mg, 1.15 mmol) was added and the mixture was stirred 3 days at rt. The mixture was evaporated, the residue was dissolved in MeOH, filtered and the clear solution was purified by rp-chromatography (XBridge, gradient of ACN/water/NH3). Corresponding fractions were lyophilized to obtain the compound **5** (42.4 mg, 44 % yield)

LC-MS (*method 10*): $^{t}R$ = 0.78 min, $[M+H]^+$ 250
$^1H$ NMR (400 MHz, DMSO-$d_6$) δ ppm 8.92 - 8.95 (m, 2 H) 8.07 (d, *J*=8.49 Hz, 1 H) 8.00 (d, *J*=1.14 Hz, 1 H) 7.84 (dd, *J*=8.62, 1.90 Hz, 1 H) 3.89 (s, 2 H) 2.89 - 2.99 (m, 2 H) 2.74 - 2.80 (m, 2 H) 2.22 - 2.35 (m, 2 H)

**2-{2-methyl-7H-pyrrolo[2,3-d]pyrimidin-5-yl}ethan-1-amine (7)**

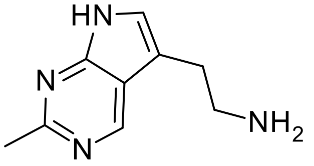

The solution of 5-(2-Isocyanato-ethyl)-2-methyl-7H-pyrrolo[2,3-d]pyrimidine **7A** (130 mg 0.64mmol) in 1M aq. NaOH (15.5 mL) was stirred at 55 °C for 10h. The mixture was evaporated. The residue was purified by flash chromatography (silica, DCM:MeOH=10:1 + 1 % NH3) to get the crude product. The crude product was purified again by flash chromatography (silica, DCM:MeOH= 10:1-5:1 +1% NH3) to obtain the compound **7** (40mg 34 % yield)
LC-MS (*method 10*): $^{t}R$ = 0.49 min, $[M+H]^+$ 177

1H NMR (400 MHz, MeOH-d4) δ ppm 8.89 (s, 1 H) 7.27 (s, 1 H) 3.12 (t, J=6.8 Hz, 1 H) 3.03 (t, J=6.8, 1 H) 2.70 (s, 3 H)

**5-(2-isocyanatoethyl)-2-methyl-7H-pyrrolo[2,3-d]pyrimidine (7A)**

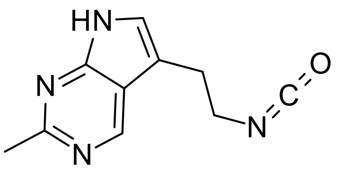

To the solution of **7B** (300mg, 1.46mmol) and TEA (221.48 mg, 2.19 mmol) in toluene (6 mL) and DMF (1 mL) was added DPPA (603.03 mg, 2.19 mmol). The white suspension was stirred at 25 °C for 2 h to give a orange suspension. t-BuOH (3 mL) was added. The mixture was stirred 8 h at 100 °C. The mixture was concentrated and poured into water (10 mL) slowly. The mixture was extracted with EA (10 mL) twice. The combined organic layers were washed with brine (20 mL) twice, dried over Na2SO4, filtered and evaporated. The residue was purified by flash chromatography (silica, EA:EtOH=10:1) to obtain compound **7A** (130 mg, 44 % yield)
LC-MS (*method 1*): $^{t}R$ = 2.46 min, $[M+H]^+$ 203

**3-{2-methyl-7H-pyrrolo[2,3-d]pyrimidin-5-yl}propanoic acid (7B)**

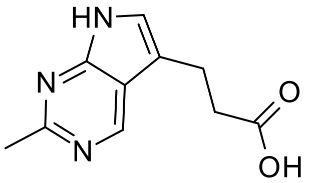

To the solution of **7C** (100,0 mg, 0.43 mmol) in EtOH (1.0 ml) and THF (2.0 ml) was added a solution of NaOH (85.74 mg, 2.14 mmol) in water (1.0 mL) at 25 $^{o}$C. The mixture was stirred at 25 $^{o}$C for 10 h.
The reaction mixture was concentrated to remove the organic solvent. The crude was diluted with EA (3 mL) and $H_2O$ (3 mL). The mixture was extracted with EA (5 mL). The aqueous phase was acidified with 1N HCl to pH=3. The crude aqueous phase was extracted with EA (5 mL). The aqueous phase was concentrated to dryness to give the crude product.
The residue was purified by flash chromatography (silica, DCM:MeOH=5:1) to get the compound **7B** (80.0 mg, 91 % yield).

LC-MS (*method 2*): $^{t}R$ = 1.31 min, $[M+H]^-$ 204

**ethyl 3-{2-methyl-7H-pyrrolo[2,3-d]pyrimidin-5-yl}propanoate 7C)**

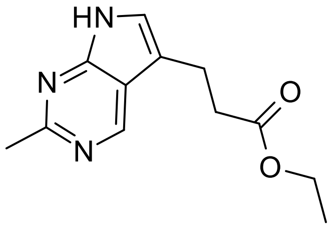

A mixture of **7D** (240.0 mg, 1.04 mmol) and 10% Pd/C (100.0 mg) in EtOH (10.0 ml) and THF (10.0 ml) was stirred under 15 psi of $H_2$ balloon at 25 $^{o}$C for 10 h.
The suspension was filtered through a pad of Celite and the pad was washed with EtOH (100 mL×2). The combined filtrates were concentrated to dryness to obtain crude compound **7C** (210.0 mg, 87 % yield) as yellow solid. The crude product was used directly for the next step without purification.

LC-MS (*method 3*): $^{t}R$ = 0.91 min, $[M+H]^+$ 234

**ethyl (2E)-3-{2-methyl-7H-pyrrolo[2,3-d]pyrimidin-5-yl}prop-2-enoate (7D)**

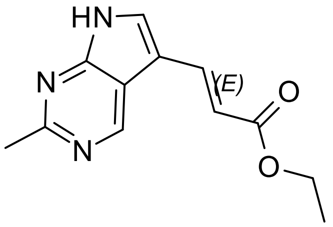

To the mixture of **7E** (2.65 g; 12.50 mmol), (E)-3-(4,4,5,5-Tetramethyl-[1,3,2]dioxaborolan-2-yl)-acrylic acid ethyl ester (3.39 g; 14.50 mmol), K3PO4 (5.30 g; 24.50 mmol) in THF (60.0 ml) and H2O (15.0 ml) was added chloro[(di(1-adamantyl)-n-butylphosphine)-2-(2-aminobiphenyl)]palladium(II) (500.9 mg, 0.75 mmol) under $N_2$. The mixture was stirred at 80 ºC for 16 h under $N_2$.

The mixture was poured into water (40 mL) slowly. The mixture was extracted with EA (50 mL) twice. The combined organic layers were washed with brine (100 mL) twice and dried over $Na_2SO_4$, filtered, and concentrated to dryness.

The residue was purified by silica gel column (PE:EA=2:1 to EA) to obtain compound **7D**. (2.0 g, 70 % yield)

LC-MS (*method 4*): $^tR$ = 1.79 min, $[M+H]^+$ 232

**5-bromo-2-methyl-7H-pyrrolo[2,3-d]pyrimidine (7E)**

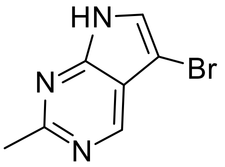

2-Methyl-7H-pyrrolo[2,3-d]pyrimidine (1.0 g, 7.51 mmol) was dissolved in DCM (25 mL). NBS (1.34 g, 7.51 mmol) was added and the mixture was stirred at rt o/n. The precipitate was filtered and the solid was dried at 50 °C to obtain compound **7E**. (1.24g, 78 % yield)

LC-MS (*method 10*): $^tR$ = 0.71 min, $[M+H]^+$ 212/214

**1-[2-(4-amino-2-methylpyrimidin-5-yl)ethyl]-3-[1-(quinoxalin-6-yl)-1H-1,2,3-triazol-4-yl]piperidin-3-ol (8)**

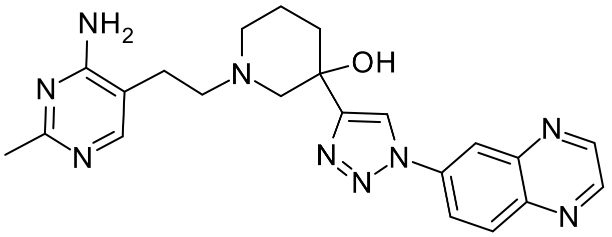

**8A** (10.0 mg, 0.02 mmol) was dissolved in acetic acid (0.25 mL). 33% HBr in acetic acid (0.4 mL, 2.32 mmol) was added and the mixture was stirred 4h at rt. The mixture was evaporated, the residue was dissolved in MeOH and the solution was purified by rp-chromatography (XBridge, ACN/water/NH3) to obtain the compound **8**. (3 mg, 39 % yield)

LC-MS (*method 10*): $^tR$ = 0.68 min, $[M+H]^+$ 432

$^1H$ NMR (400 MHz, DMSO-$d_6$) δ ppm 9.05 (d, *J*=1.77 Hz, 1 H) 9.01 - 9.03 (m, 1 H) 8.94 - 8.97 (m, 1 H) 8.63 (d, *J*=2.41 Hz, 1 H) 8.50 (dd, *J*=9.06, 2.47 Hz, 1 H) 8.32 (d, *J*=9.00 Hz, 1 H) 7.79 - 7.84 (m, 1 H) 6.63 (s, 2 H) 5.11 (s, 1 H) 2.96 (br d, *J*=10.65 Hz, 1 H) 2.64 (br d, *J*=10.90 Hz, 1 H) 2.53 (s, 4 H) 2.43 (br d, *J*=6.84 Hz, 1 H) 2.20 (s, 4 H) 1.69 - 1.85 (m, 2 H) 1.48 - 1.57 (m, 1 H)

**benzyl N-[5-(2-{3-hydroxy-3-[1-(quinoxalin-6-yl)-1H-1,2,3-triazol-4-yl]piperidin-1-yl}ethyl)-2-methylpyrimidin-4-yl]carbamate (8A)**

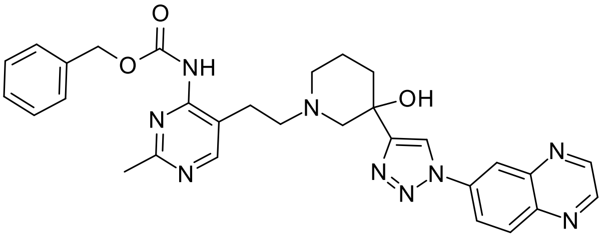

**8E** (70.0 mg, 0.25 mmol) and **8B** (72.7 mg, 0.25 mmol) were dissolved in DCM (5.0 mL) and acetic acid (0.05 mL). Molsieve and NaBH(OAC)3 (104.0 mg, 0.49 mmol) was added and the mixture was stirred at rt o/n. The mixture was evaporated, the residue was dissolved in DMF and purified by rp-chromatography (XBridge, ACN/water/NH3). Corresponding fractions were lyophilized to obtain compound **8A**. (59.0 mg, 43 % yield)

LC-MS (*method 10*): $^{t}$R = 0.91 min, $[M+H]^+$ 566

**3-[1-(quinoxalin-6-yl)-1H-1,2,3-triazol-4-yl]piperidin-3-ol (8B)**

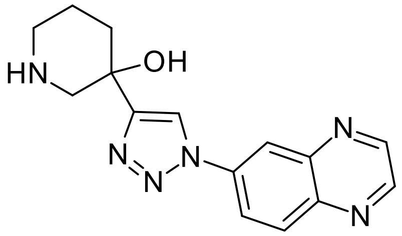

4M HCl in MeOH (150 mL) was added to **8C** (4.5 g, 11.35 mmol) at 0 °C. The mixture was stirred 2h at rt and was then evaporated. The residue was diluted with EA (100 mL) and was basified to pH=8 with NH3. The precipitate was filtered and washed with EA (30 mL) twice. The solid was diluted with MeOH (30 mL) and stirred 20min at rt. The precipitate was filtered, washed with cold MeOH (10mL) and dried to obtain compound **8B**. (5.0 g, 84 % yield)

LC-MS (*method 5*): $^{t}$R = 1.90 min, $[M+H]^+$ 297

**tert-butyl 3-hydroxy-3-[1-(quinoxalin-6-yl)-1H-1,2,3-triazol-4-yl]piperidine-1-carboxylate (8C)**

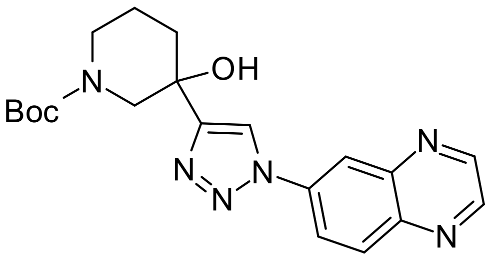

6-Bromo-quinoxaline (10.0g, 47.84 mmol), **8D** (10.78 g, 47.84 mmol, CuI (1.36 g, 7.18 mmol), NaN3 (6.22 g, 95.67 mmol), Na ascorbate 095 g, 4.78 mmol) were diluted with TEA (5.8 g, 57.40 mmol), trans-N,N'-dimethylcyclohexane-1,2-diamine (1.36 g, 9.57 mmol), water (20 mL) and MeOH (200 mL) under nitrogen. The mixture was stirred 16 h at 55 °C under nitrogen. The mixture was filtered at rt and the pad was washed with MeOH (50 mL) zwice. The combined filtrates were concentrated and extracted with EA twice. The organic layers were dried over Na2SO4, filtrated and evaporated. The residue was purified by flash chromatography (silica, PE/EA=2/1-0/100) to obtain compound **8C** (12g, 63 % yield)
LC-MS (*method 4*): $^{t}$R = 2.071 min, $[M+H]^+$ 397

**tert-butyl 3-ethynyl-3-hydroxypiperidine-1-carboxylate (8D)**

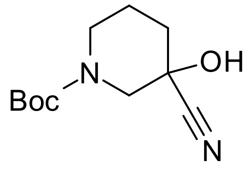

The solution of 1-boc-3-piperidone (65.0 g, 326.23 mmol) in THF (500 mL) was added dropwise 0.5 mol/L ethynylmagnesium bromide (978.7 mL, 489.34 mmol) at 0 °C under n2. The mixture was stirred 4h at 0 °C. The mixture was quenched with sat. NH4Cl (500 mL) and was extracted with EA (3x 600 ml). The organic layers were washed with brine (1000 mL), dried over Na2SO4, filtered and evaporated to obtain compound **8D**. (75 g, 100 % yield)

LC-MS (*method 8*): $^{t}$R = 1.048 min, M+Na+ 362

**benzyl 6-hydroxy-2-methyl-5H,6H,7H-pyrrolo[2,3-d]pyrimidine-7-carboxylate (8E)**

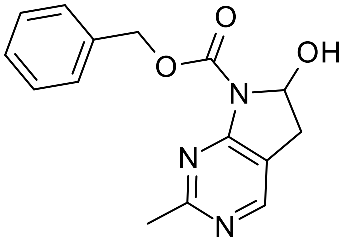

A solution of **8F** (1.05 g, 3.35 mmol) in 6M HCl (10 mL) was stirred 5h at 25 °C. The mixture was adjusted to pH=8~9 by aq. NaHCO3. The mixture was extraczed with EA (3x 10 mL). The combined organic layers were dried over Na2SO4, filtered and evaporated. The residue was purified by flash chromatography (silica, PE:EA=10:1-3:1) to obtain compound **8E** (0.6 g, 63 % yield)

LC-MS (*method 6*): $^{t}$R = 0.99 min, [M+Na]+ 308

**benzyl N-{5-[(1E)-2-ethoxyethenyl]-2-methylpyrimidin-4-yl}carbamate (8F)**

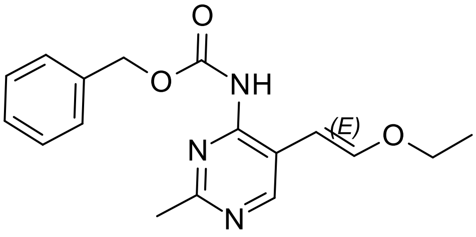

**8G** (3.0 g, 16.74 mmol) was dissolved in THF (30 mL). CbzCl (3.43 g, 20.09 mmol) and TEA (5.01 g, 50.22 mmol) were added and the mixture was stirred 5h at 25 °C. The mixture was evaporated. The residue was purified by flash chromatography (PE:EA=5:1-0:1) to obtain compound **8F**. (2.5 g, 48 % yield)
TLC: SiO2, PE:EA=1:1, Rf= 0.5

LC-MS (*method 7*): $^{t}$R = 1.229 min, $[M+H]^+$ 314

**5-[(1E)-2-ethoxyethenyl]-2-methylpyrimidin-4-amine (8G)**

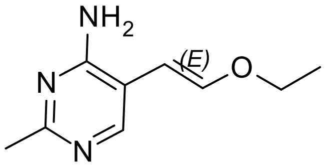

5-Bromo-2-methyl-pyrimidin-4-ylamine (5.0 g, 26,59 mmol) was dissolved in THF (100 mL) Water (25 mL), K3PO4 (11,27 g, 53,18 mmol), 2-((E)-2-Ethoxy-vinyl)-4,4,5,5-tetramethyl-[1,3,2]dioxaborolane (6,32 g, 31,91 mmol) and Chloro[(di(1-adamantyl)-N-butylphosphine)-2-(2-aminobiphenyl)]palladium(II) (0,57 g, 2,66 mmol) were added and the mixture was stirred 12h at 80 °C under N2.

The mixture was diluted with water (500mL), extracted with EA (3 x 150mL). The combined organic layers were dried over Na2SO4, filtered and evaporated.

The residue was purified by flash chromatography (silica, PE:EtOAc=10:1-2:1) to get the compound **8G** (3.0 g, 92 % yield)

LC-MS (*method 8*): $^{t}$R = 0.69 min, $[M+H]^+$ 180

**5-[(3-{[1-(1H-1,3-benzodiazol-2-yl)piperidin-4-yl]oxy}pyrrolidin-1-yl)methyl]-2-methylpyrimidin-4-amine (9)**

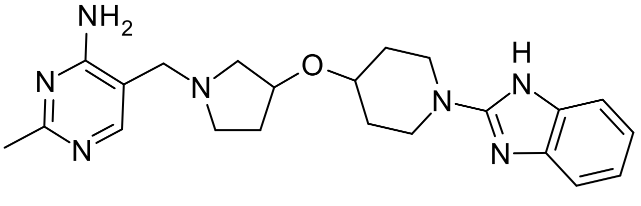

A mixture of 2-chlorobenzimidazole (30 mg, 0.20 mmol) and **9A** (57.3 mg, 0.20 mmol) was stirred at 120 °C o/n. The mixture was diluted with DMF and the solution was purified by rp-chromatography (XBridge, ACN/water/NH3) to obtain compound **9**. (42 mg, 52% yield)

LC-MS (*method 10*): $^{t}R$ = 0.72 min, $[M+H]^+$ 408
$^{1}H$ NMR (400 MHz, DMSO-$d_6$) δ ppm 7.86 (s, 1 H) 7.12 - 7.19 (m, 2 H) 6.86 - 6.94 (m, 2 H) 6.72 (s, 2 H) 4.16 - 4.24 (m, 1 H) 3.79 - 3.90 (m, 2 H) 3.44 - 3.56 (m, 2 H) 3.39 - 3.43 (m, 2 H) 3.16 (ddd, *J*=13.12, 10.01, 3.11 Hz, 2 H) 2.67 (dd, *J*=9.89, 6.21 Hz, 1 H) 2.52 - 2.60 (m, 1 H) 2.35 - 2.43 (m, 2 H) 2.29 (s, 3 H) 2.00 - 2.10 (m, 1 H) 1.82 - 1.94 (m, 2 H) 1.60 - 1.70 (m, 1 H) 1.38 - 1.49 (m, 2 H)

**5 2-methyl-5-{[3-(piperidin-4-yloxy)pyrrolidin-1-yl]methyl}pyrimidin-4-amine1 (9A)**

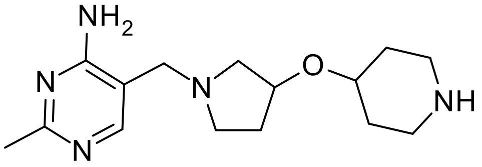

**9B** (420 mg, 1.09 mmol) was dissolved in MeOH (15 mL). 1M NaOH (5.4 mL, 5.42 mmol) was added and the mixture was stirred 4h at rt. Water was added and the mixture was extracted with DCM. Organic layers were dried over Na2SO4, filtrated and evaporated. The residue was purified by flash chromatography (silica, DCM:MeOH=99:1-87:13 to obtain compound **9A**. (195 mg, 62% yield)

LC-MS (*method 10*): $^{t}R$ = 0.70 min, $[M+H]^+$ 292

**1-[4-({1-[(4-amino-2-methylpyrimidin-5-yl)methyl]pyrrolidin-3-yl}oxy)piperidin-1-yl]-2,2,2-trifluoroethan-1-one (9B)**

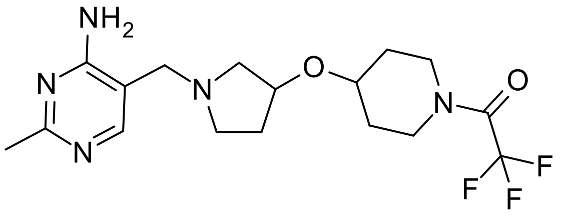

F3

**1A** (300 mg, 2.19 mmol) and 9C (582.5 mg, 2.19 mmol) were dissolved in acetic acid (0.12 mL, 2.19 mmol) and DCM (6 mL) and stirred 1h at rt. NaBH(OAc)3 (0.73 g, 3.28 mmol) was added and the mixture was stirred at rt o/n. Water was added and the mixture was extracted with DCM. Organic layer was dried over Na2SO4, filtrated and evaporated to obtain compound 14B. (420mg, 50% yield)

LC-MS (*method 10*): $^{t}R$ = 0.87 min, $[M+H]^+$ 388

**2,2,2-trifluoro-1-[4-(pyrrolidin-3-yloxy)piperidin-1-yl]ethan-1-one, hydrochloride salt (9C)**

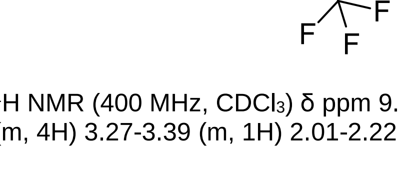

A mixture of **9D** (6.50g, 17.7 mmol) and HCl (50mL 4mol/L in dioxane, 20 mmol) was stirred 2h at 15 °C. The mixture was evaporated to obtain crude compound **9C** as HCl salt which was used without further purification

TLC: SiO2, PE:EA=2:1, Rf=0
LC-MS (*method 5*): $^{t}R$ = 2.03 min, M+H+ 267
$^{1}H$ NMR (400 MHz, $CDCl_3$) δ ppm 9.62 (br s, 1 H) 9.72 (br s, 1 H) 4.36 (s, 1 H) 3.62-3.79 (m, 4 H) 3.39-3.58 (m, 4H) 3.27-3.39 (m, 1H) 2.01-2.22 (m, 2 H) 1-63-1.94 (m, 4 H)

**tert-butyl 3-{[1-(2,2,2-trifluoroacetyl)piperidin-4-yl]oxy}pyrrolidine-1-carboxylate (9D)**

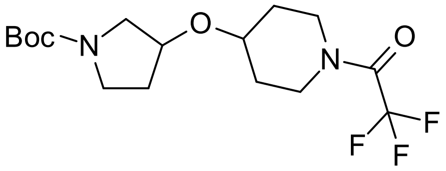


**9E** (7.0 g, 25.89 mmol) was dissolved in THF (80 mL) and TEA (3.92 g, 38.84 mmol). (CF3)2CO (7.1 g, 33.66 mmol) was added slowly at 0 °C. The mixture was stirred 8h at rt before quenched with water (50 mL). The mixture was extracted with EA (2x 150 mL), organic layers washed with brine (150 mL), dried over Na2SO4, filtrated and evaporated. The residue was purified by flash chromatography (silica, PE:EA=5:1-1:1 to obtain compound **9D**. (6.5 g, 69% yield).
TLC: silica, PE:EA=3:1, Rf=0.2

$^1$H NMR (400 MHz, $CDCl_3$) δ ppm 4.12-4.2 (m, 1 H) 3.60-3.80 (m, 4 H) 3.28-3.55 (m, 5H) 1.91-2.00 (m, 2H) 1.80-1.90 (m, 2 H) 1.65-1.75 (m, 4 H) 1.48 (s, 9 H)

**tert-butyl 3-(piperidin-4-yloxy)pyrrolidine-1-carboxylate (9E)**

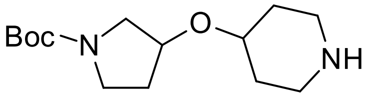


**9F** (2.3 g, 6.41 mmol) was dissolved in MeOH (40 mL). Pd/C (200 mg) was added and the mixture was hydrogenated 4h at 50 °C and 50 psi H2 pressure. The mixture was filtrated, evaporated and dried at high vacuo to obtain compound **9E** (1.7 g, 98% yield)
LC-MS (*method 12*): $^tR$ = 0.72 min, $[M+H]^+$ 271

**tert-butyl 3-(pyridin-4-yloxy)pyrrolidine-1-carboxylate (9F)**

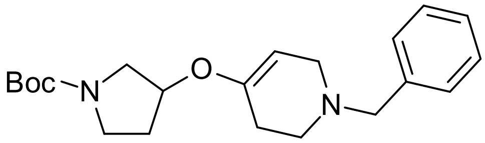


**9G** (1.6 g, 3.68 mmol) was dissolved in MeOH (20 mL). The mixture was cooled to 0°C and NaBH4 (470.3 mg, 12.13 mmol) was added slowly. The mixture was stirred o/n whereby rt was achieved. The mixture was quenched with water, sat. NaHCO3 was added and the mixture was extracted 3x with EA. Organic layers were dried over Na2SO4, filtrated and evaporated to obtain compound **9F**. (1.24 g, 94% yield)

LC-MS (*method 12*): $^tR$ = 0.83 min, $[M+H]^+$ 359

**tert-butyl 3-(pyridin-4-yloxy)pyrrolidine-1-carboxylate (9G)**

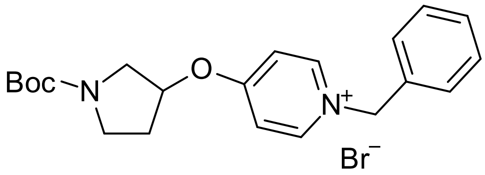


**9H** (1.7 g, 6.43 mmol) was dissolved in DCM (30 mL). (bromomethyl)benzene (0.84 mL, 7.05 mmol) was added and the mixture was stirred at rt o/n. The mixture was evaporated to obtain compound **9G** (2.8 g, 100% yield)

LC-MS (*method 12*): $^tR$ = 0.82 min, $[M+H]^+$ 355

**tert-butyl 3-(pyridin-4-yloxy)pyrrolidine-1-carboxylate (9H)**

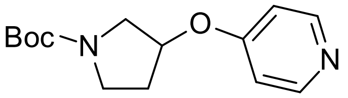

4-(pyrrolidin-3-yloxy)pyridine dihydrochloride (1.9 g, 8.01 mmol) was mixed with DCM (40 mL). TEA (3.37 mL, 24.04 mmol) and BOC2O (1.92 g, 8.81 mmol) were added and the mixture was stirred 3h at rt. Water was added and the mixture was extracted 3x with DCM. Organic layers were dried over Na2SO4, filtrated and evaporated to obtain compound 14H (2.08g, 98% yield)
LC-MS (*method 12*): $^{t}R$ = 0.72 min, $[M+H]^+$ 265

**6-[(3-{[1-(1H-1,3-benzodiazol-2-yl)piperidin-4-yl]oxy}pyrrolidin-1-yl)methyl]quinoxaline (10)**

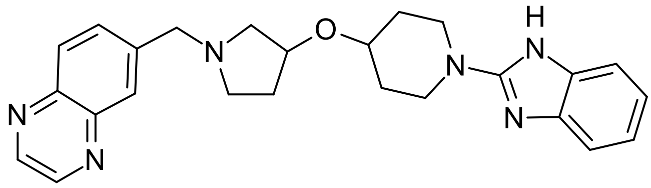

**10A** (120 mg, 0.37 mmol) was dissolved in THF (5 mL). Quinoxaline-6-carbaldehyde (58.8 mg, 0.37 mmol) was added and the mixture was acidified to pH=5 with acetic acid. The mixture was stirred 30 min at rt. NaBH(OAc)3 (98.5 mg, 0.47 mmol) was added and the mixture was stirred at rt o/n. NaBH(OAc)3 (98.5 mg, 0.47 mmol) was added again and the mixture was stirred 3 days at rt. The mixture was quenched with water and evaporated. The residue was dissolved in MeOH, filtrated, purified by rp-chromatography (XBridge, ACN/water/NH3) and lyophilized to obtain compound **10** (110mg, 69% yield)
LC-MS (*method 13*): $^{t}R$ = 0.39 min, $[M+H]^+$ 429
$^1H$ NMR (400 MHz, DMSO-$d_6$) δ ppm 8.92 - 8.94 (m, 1 H) 8.90 - 8.92 (m, 1 H) 8.06 (d, *J*=8.59 Hz, 1 H) 7.99 (s, 1 H) 7.81 - 7.87 (m, 1 H) 7.12 - 7.20 (m, 2 H) 6.89 (dd, *J*=5.68, 3.16 Hz, 2 H) 5.95 - 6.10 (m, 1 H) 4.24 (tt, *J*=7.07, 3.79 Hz, 1 H) 3.79 - 3.89 (m, 4 H) 3.53 (ddd, *J*=12.57, 8.53, 3.66 Hz, 2 H) 2.78 - 2.85 (m, 2 H) 2.60 - 2.68 (m, 2 H) 2.09 (td, *J*=13.77, 7.58 Hz, 1 H) 1.83 - 1.94 (m, 2 H) 1.64 - 1.73 (m, 1 H) 1.38 - 1.50 (m, 2 H)

**2-[4-(pyrrolidin-3-yloxy)piperidin-1-yl]-1H-1,3-benzodiazole, hydrochloride salt (10A)**

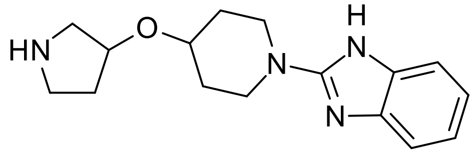

**10B** (200 mg, 0.52 mmol) was dissolved in DCM (10 mL) and MeOH (1 mL). HCl (3 mL 4mol/L in dioxane) was added and the mixture was stirred 1h at rt. The mixture was evaporated to obtain compound **10A** (167 mg, 100% yield)

LC-MS (*method 10*): $^{t}R$ = 0.82 min, $[M+H]^+$ 287

**tert-butyl 3-{[1-(1H-1,3-benzodiazol-2-yl)piperidin-4-yl]oxy}pyrrolidine-1-carboxylate (10B)**

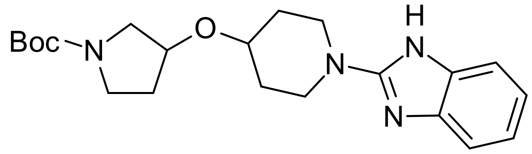

A mixture of 2-chlorobenzimidazole (1.0 g, 6.55 mmol) and **9E** (2.12 g, 7.87 mmol) in DMF ( 20mL) and DIPEA (3.38 mL, 20 mmol) was stirred at 120 °C o/n. The mixture was quenched with water and extracted with 3x EA. Organic layers were dried over Na2SO4, filtrated and evaporated. The residue was purified by flash chromatography (silica, DCM:MeOH=100:0-93:7). Fractions containing product were pooled and evaporated to obtain compound **10B** (1.14 g, 45% yield)

LC-MS (*method 10*): $^{t}R$ = 0.96 min, $[M+H]^+$ 387

**General syntheis of compounds 11-13 and BI-5232:**

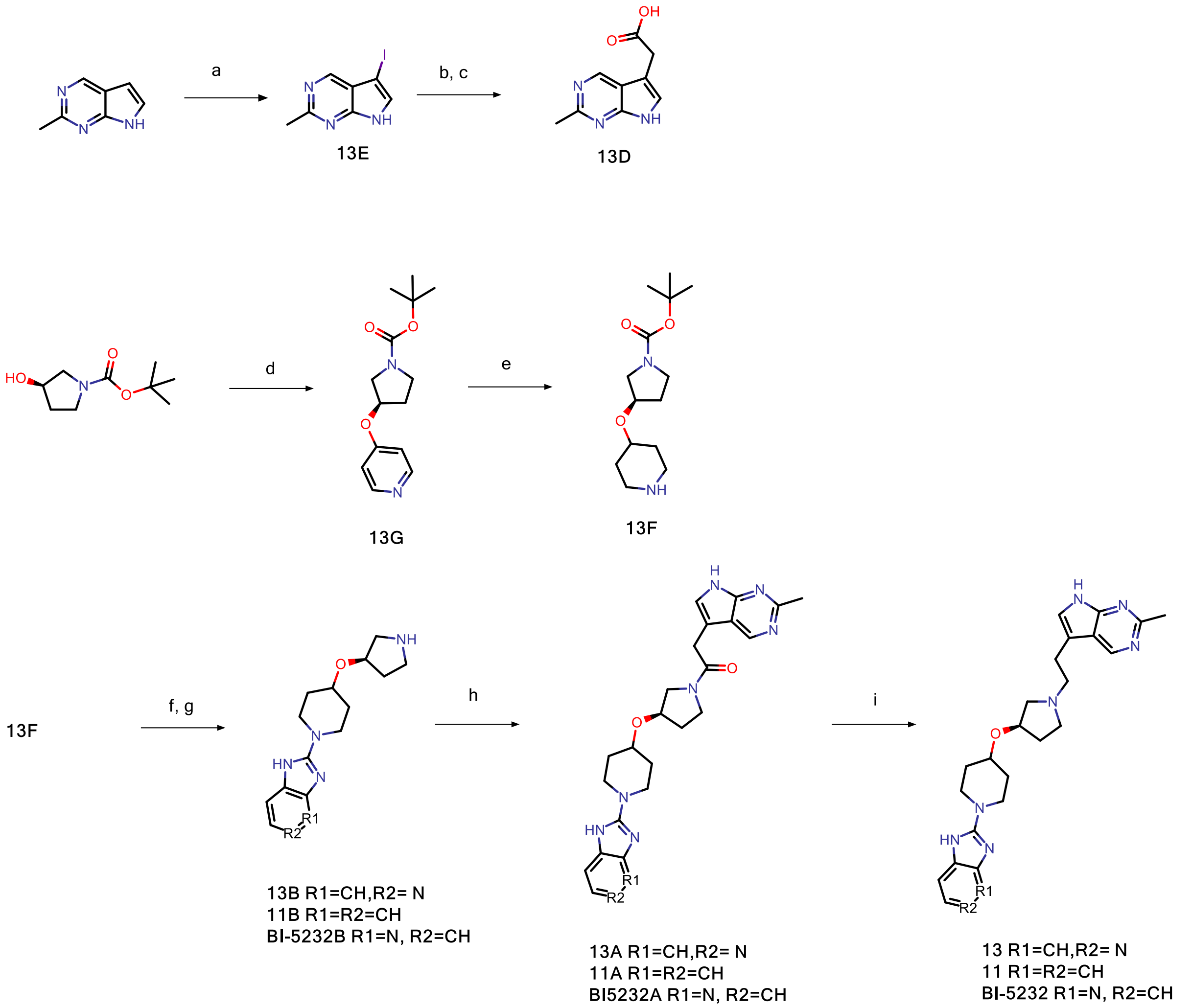


[a]Reagents and conditions (shown for compound **13**): (a) NIS, $CH_2Cl_2$, 81% yield; (b) 2-tert-butoxy-2-oxoethylzinc chloride, SPHOS Pd G3, THF; (c) TFA, $CH_2Cl_2$ (1:1), 98% yield over 2steps; (d) 4-fluoropyridine, NaH 50% dispersion, THF, 60°C, 94% yield; (e) Nishimura's catalyst, HOAc, MeOH, 98% yield; (f) 2-chloro-3H-imidazo[4,5-c]pyridine, DIPEA, MeOH, 160 °C, microwave irradiation, 42% yield; (g) 4N HCl in dioxane, $CH_2Cl_2$, 79% yield; (h) HATU, DIPEA, DMF, 57% yield; (i) $LiAlH_4$, THF/DMSO (10:1), 45% yield.

**1-{3H-imidazo[4,5-c]pyridin-2-yl}-4-{[(3R)-1-(2-{2-methyl-7H-pyrrolo[2,3-d]pyrimidin-5-yl}ethyl)pyrrolidin-3-yl]oxy}piperidine (13)**

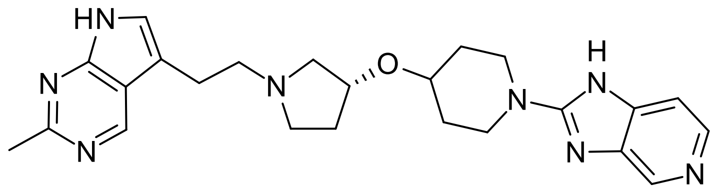

**BI-5232A** (150 mg, 0.33 mmol) was dissolved in THF ( 3mL) and DMSO (0.3 mL). LiAlH4 (120 mg, 3.16 mmol) was added slowly and the mixture was stirred 10 min at rt. The mixture was quenched with 120 µL water and 120 µL 4 mol/L NaOH and the mixture was stirred 3 min at rt. 360µL water was added and the suspension was filtered. The filtrate was evaporated and the residue was purified by rp-chromatography (XBridge, ACN/water/NH3). Corresponding fractions were evaporated to obtain compound **13** (66 mg, 45% yield)

LC-MS (*method 10*): $^tR$ = 0.74 min, $[M+H]^+$ 447

$^1H$ NMR (400 MHz, DMSO-$d_6$) δ ppm 11.36 - 11.68 (m, 1 H) 8.88 (s, 1 H) 8.35 (s, 1 H) 8.01 (d, *J*=5.32 Hz, 1 H) 7.22 (s, 1 H) 7.14 (d, *J*=5.45 Hz, 1 H) 4.20 (tt, *J*=6.95, 3.69 Hz, 1 H) 3.87 - 3.97 (m, 2 H) 3.56 (td, *J*=8.24,

4.06 Hz, 2 H) 2.76 - 2.88 (m, 3 H) 2.65 - 2.71 (m, 2 H) 2.60 (s, 4 H) 1.97 - 2.10 (m, 1 H) 1.90 (br dd, *J*=8.30, 3.87 Hz, 2 H) 1.59 - 1.69 (m, 1 H) 1.40 - 1.52 (m, 2 H)

**1-[(3R)-3-[(1-{3H-imidazo[4,5-c]pyridin-2-yl}piperidin-4-yl)oxy]pyrrolidin-1-yl]-2-{2-methyl-7H-pyrrolo[2,3-d]pyrimidin-5-yl}ethan-1-one (13A)**

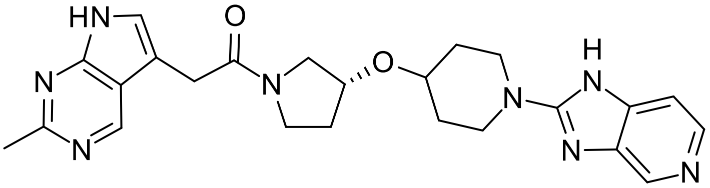


**13D** (110 mg, 0.57 mmol) was dissolved in DMF (4 mL). DIPEA (0.3 mL, 1.72 mmol) and **13B** (165 mg, 0.57 mmol) were added and the mixture was stirred 1 min at rt. HATU (218 mg, 0.57 mmol) was added and the mixture was stirred 1h at rt. The solution was purified by rp-chromatography (XBridge, ACN/water/NH3), fractions were evaporated to obtain compound **13A** (150 mg, 57% yield).
LC-MS (*method 10*): $^{t}$R = 0.68 min, $[M+H]^{+}$ 461

**1-{3H-imidazo[4,5-c]pyridin-2-yl}-4-[(3R)-pyrrolidin-3-yloxy]piperidine (13B)**

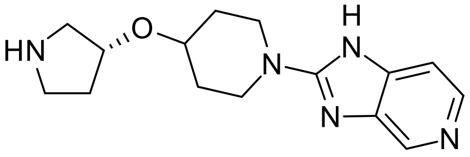


**13C** (290 mg, 0.75 mmol) was dissolved in DCM (5 mL). HCl (0.9 mL 4mol/L in dioxane, 4.81 mmol) was added and the mixture was stirred 1h at rt. 1 mol/L NaOH (1.8 mL, 1.8 mmol) was added and the mixture was extracted with DCM. Organic layers were dried over Na2SO4, filtrated and evaporated to obtain compound **13B** (170 mg, 79% yield).

LC-MS (*method 10*): $^{t}$R = 0.64 min, $[M+H]^{+}$ 288

**tert-butyl (3R)-3-[(1-{3H-imidazo[4,5-c]pyridin-2-yl}piperidin-4-yl)oxy]pyrrolidine-1-carboxylate (13C)**

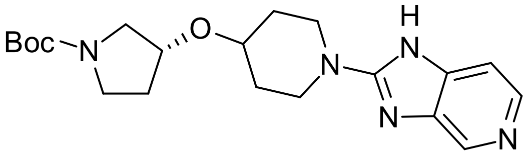


2-chloro-3H-imidazo[4,5-c]pyridine (312 mg, 2.03 mmol) and **13F** (500 mg, 1.85 mmol) were dissolved in MeOH (20 mL) and DIPEA (0.95 mL, 5.55 mmol). The mixture was stirred 2.5 h at 160 °C in the microwave. The mixture was evaporated and the residue was purified by rp-chromatography Sunfire, ACN/water/TFA) to obtain compound **13C** (300 mg, 42% yield.
LC-MS (*method 10*): $^{t}$R = 0.86 min, $[M+H]^{+}$ 388

**2-{2-methyl-7H-pyrrolo[2,3-d]pyrimidin-5-yl}acetic acid (13D)**

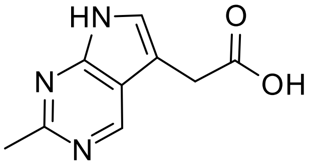


Step 1:
A mixture of 2-tert-butoxy-2-oxoethylzinc chloride (20 mL 0.5 mol/L in THF, 10.0 mmol), **13E** (500 mg, 1.93 mmol) and SPHOS Pd G3 (75 mg, 0.09 mmol) was stirred 4h at 70°C. The mixture was evaporated. 2-tert-butoxy-2-oxoethylzinc chloride (20 mL 0.5 mol/L in THF, 10.0 mmol) and SPHOS Pd G3 (75 mg, 0.09 mmol) were added, and the mixture was stirred 2h at 70°C. The mixture was evaporated. The residue was diluted with 50 mL EA and 50 mL sat. NH4Cl-solution, Celite was added, and the mixture was filtrated. The filtrate was extracted 3x with EA, organic layers were combined, dried over Na2SO4, filtrated and evaporated. The residue was suspended in diethylether and the precipitate was isolated by filtration to obtain crude intermediate **13D-1** (803 mg, 168% yield), which was used directly in the next step.
LC-MS (*method 12*): $^{t}$R = 0.62 min, $[M+H]^{+}$ 248

Step 2:
Intermediate **13D-1** was diluted with DCM:TFA=1:1 (15 mL) and stirred 1h at rt. The mixture was evaporated. The residue was suspended in ACN, the precipiitate was filtered and washed with little ACN and diethylether to obtain compound **13D** (284 mg, 77% yield).

LC-MS (*method 12*): $^{t}R$ = 0.18 min, $[M+H]^+$ 192

**5-iodo-2-methyl-7H-pyrrolo[2,3-d]pyrimidine (13E)**

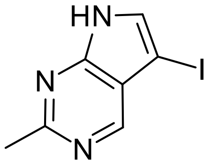


2-Methyl-7H-pyrrolo[2,3-d]pyrimidine (300 mg, 2.25 mmol) was suspended in DCM (10 mL). NIS (510 mg, 2.27 mmol) was added and the mixture was stirred at rt o/n. The mixture was concentrated to half of volume under reduced pressure, the precipitate was filtered and washed with diethylether to obtain compound **13E** (470 mg, 81% yield).
LC-MS (*method 12*): ): $^{t}R$ = 0.38 min, $[M+H]^+$ 260

**tert-butyl (3R)-3-(piperidin-4-yloxy)pyrrolidine-1-carboxylate (13F)**

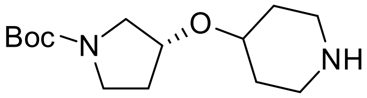


**13G** (3.0 g, 11.35 mmol) was dissolved in MeOH (50 mL) and acetic acid (0.6 mL, 10.47 mmol). Nishimura catalyst (300 mg) was added and the mixture was hydrogenated with 60 psi H2 pressure at rt o/n. The mixture was filtrated. Active charcoal was added and the filtrate was filtrated again. The filtrate was evaporated and the residue was dissolved in DCM. The solution was washed with 0.5 mol/L NaOH and water, dried over Na2SO4, filtrated and evaporated to obtain compound **13F** (3 g, 98% yield).

LC-MS (*method 10*): $^{t}R$ = 0.87 min, $[M+H]^+$ 271

**tert-butyl (3R)-3-(pyridin-4-yloxy)pyrrolidine-1-carboxylate (13G)**

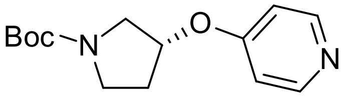


4-fluoropyridine hydrochloride (2.0 g, 14.98 mmol) and tert-butyl (3R)-3-hydroxypyrrolidine-1-carboxylate (2.8 g, 14.98 mmol) were dissolved in THF (50 mL). NaH 50% dispersion (1.8 g, 37.44 mmol) was added. The mixture was stirred 15 min at rt and was then stirred at 60°C o/n. The mixture was quenched with water at rt and THF was distilled off. The mixture was extracted 3x with EA.Organic layer was dried over Na2SO4, filtrated and evaporated. The residue was purified by flash chromatography (silica, DCM:MeOH=100:0-95:5) to obtain compound **13G** (3.73 g, 94% yield).

LC-MS (*method 10*): $^{t}R$ = 0.92 min, $[M+H]^+$ 265

Compounds 11, 12 and BI-5232 were synthesized using the general procedure for compound 13 with exceptions as noted.

**2-(4-{[(3R)-1-(2-{2-methyl-7H-pyrrolo[2,3-d]pyrimidin-5-yl}ethyl)pyrrolidin-3-yl]oxy}piperidin-1-yl)-1H-1,3-benzodiazole (11)**

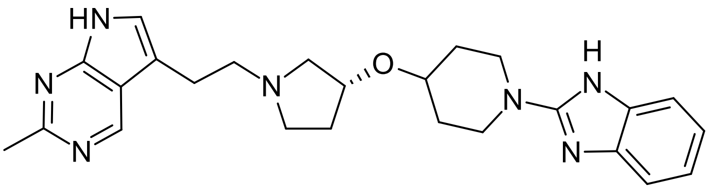

Synthesis is analogous to that described for compound **13** using **11A** (65 mg, 0.14 mmol) as starting material to obtain compound **11** (22 mg, 35% yield).
LC-MS (*method 13*): $^{t}R$ = 0.33 min, $[M+H]^+$ 446
Chiral chromatography (*Chiral SFC method 1*): $^{t}R$ = 5.44 min,>98 %ee
$^{1}$H NMR (400 MHz, DMSO-$d_6$) δ ppm 11.15 - 11.61 (m, 2 H) 8.88 (s, 1 H) 7.22 (s, 1 H) 7.13 - 7.19 (m, 2 H) 6.90 (br s, 2 H) 4.16 - 4.23 (m, 1 H) 3.81 - 3.90 (m, 2 H) 3.51 - 3.59 (m, 1 H) 3.12 - 3.22 (m, 2 H) 2.76 - 2.88 (m, 3 H) 2.64 - 2.71 (m, 2 H) 2.58 - 2.63 (m, 4 H) 1.99 - 2.09 (m, 1 H) 1.90 (br dd, *J*=7.98, 3.93 Hz, 2 H) 1.59 - 1.68 (m, 1 H) 1.39 - 1.51 (m, 2 H)

**1-[(3R)-3-{[1-(1H-1,3-benzodiazol-2-yl)piperidin-4-yl]oxy}pyrrolidin-1-yl]-2-{2-methyl-7H-pyrrolo[2,3-d]pyrimidin-5-yl}ethan-1-one (11A)**

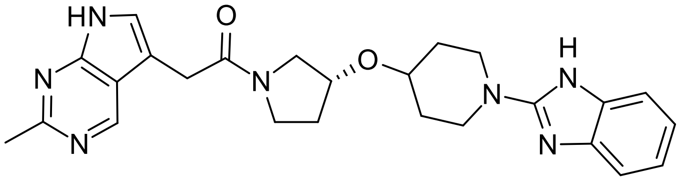

Synthesis is analogous to that described for compound **13A** using **13D** (133 mg, 0.70 mmol) and **11B** (250 mg, 0.70 mmol) as starting materials to obtain compound **11A** (176 mg, 55% yield).
LC-MS (*method 10*): $^{t}R$ = 0.78 min, $[M+H]^+$ 460

**2-{4-[(3R)-pyrrolidin-3-yloxy]piperidin-1-yl}-1H-1,3-benzodiazole dihydrochloride salt (11B)**

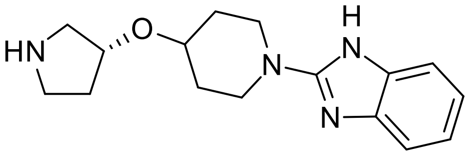

Synthesis is analogous to that described for compound **13B** using **11C** (370 mg,0.96 mmol) as starting material to obtain compound **11B** (250 mg, 73% yield).
LC-MS (*method 12*): $^{t}R$ = 0.58 min + 0.62 min, $[M+H]^+$ 287

**tert-butyl (3R)-3-{[1-(1H-1,3-benzodiazol-2-yl)piperidin-4-yl]oxy}pyrrolidine-1-carboxylate (11C)**

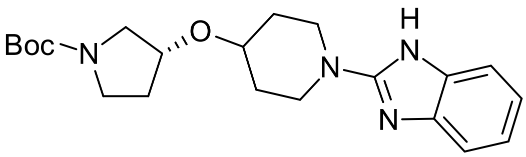

Synthesis is analogous to that described for compound **13C** using 2-chloro-benzimidazole (2.5 g, 16.39 mmol) and **13F** (4.43 g, 16.39 mmol) as starting materials to obtain compound **11C** (2.3 g, 36% yield).
LC-MS (*method 10*): $^{t}R$ = 0.97 min, $[M+H]^+$ 387

**2-(4-{[(3S)-1-(2-{2-methyl-7H-pyrrolo[2,3-d]pyrimidin-5-yl}ethyl)pyrrolidin-3-yl]oxy}piperidin-1-yl)-1H-1,3-benzodiazole (12)**

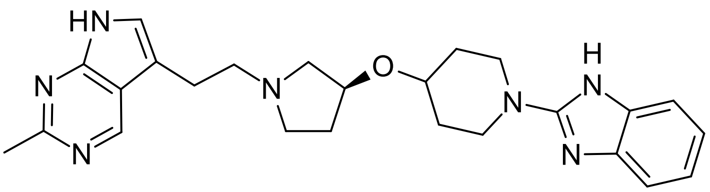

Compound 12 was prepared via the same route as 11, starting with the commercially available enantiomeric tert-butyl (3S)-3-hydroxypyrrolidine-1-carboxylate. Compound 12 was obtained in the final step from 1-[(3S)-3-{[1-(1H-1,3-benzodiazol-2-yl)piperidin-4-yl]oxy}pyrrolidin-1-yl]-2-{2-methyl-7H-pyrrolo[2,3-d]pyrimidin-5-yl}ethan-1-one (11A) (130 mg, 0.283 mmol) to give compound 12 (29.6 mg, 23% yield).
LC-MS (*method 10*): $^{t}R$ = 0.73 min, $[M+H]^+$ 446

$[M+H]^+$ 446; Chiral chromatography (*Chiral SFC method 1*): $^{t}R$ = 4.5 min,>98 %ee

H NMR (400 MHz, DMSO-d6) δ ppm 11.51 (br s, 1 H) 11.27 (br s, 1 H) 8.87 (s, 1 H) 7.22 (s, 1 H) 7.13 - 7.19 (m, 2 H) 6.84-696 (m, 2 H) 4.15 - 4.25 (m, 1 H) 3.80 - 3.91 (m, 2 H) 3.51 - 3.59 (m, 1 H) 3.34-3.49 (m, 2H) 3.10 - 3.23 (m, 2 H) 2.74 - 2.94 (m, 4 H) 2.62 - 2.74 (m, 2 H) 2.57 - 2.61 (m, 4 H) 1.97 - 2.10 (m, 1 H) 1.90 (br dd, J=8.24, 3.68 Hz, 2 H) 1.56 - 1.70 (m, 1 H) 1.38 - 1.53 (m, 2 H)

**1-{1H-imidazo[4,5-b]pyridin-2-yl}-4-{[(3R)-1-(2-{2-methyl-7H-pyrrolo[2,3-d]pyrimidin-5-yl}ethyl)pyrrolidin-3-yl]oxy}piperidine (BI-5232)**

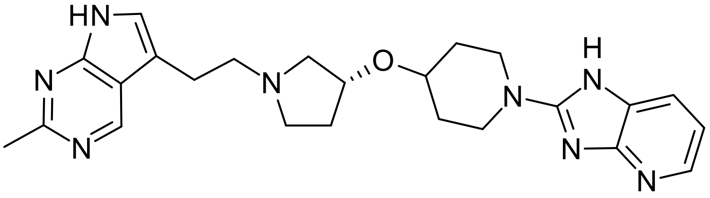

Synthesis is analogous to that described for compound 13 using **BI-5232A** (120 mg, 0.14 mmol) as starting material to obtain compound **BI-5232** (26 mg, 22% yield).
LC-MS (*method 10*): $^{t}R$ = 0.66 min, $[M+H]^+$ 447
$^1H$ NMR (400 MHz, DMSO-$d_6$) δ ppm 11.50 (br s, 1H) 8.88 (s, 1 H) 7.88 (dd, *J*=5.07, 1.52 Hz, 1 H) 7.40 (dd, *J*=7.60, 1.52 Hz, 1 H) 7.22 (s, 1 H) 6.86 (dd, *J*=7.67, 5.01 Hz, 1 H) 4.16 - 4.24 (m, 1 H) 3.88 - 3.96 (m, 2 H) 3.52 - 3.60 (m, 2 H) 2.76 - 2.88 (m, 3 H) 2.64 - 2.72 (m, 2 H) 2.58 - 2.63 (m, 4 H) 1.99 - 2.09 (m, 1 H) 1.85 - 1.95 (m, 2 H) 1.59 - 1.69 (m, 1 H) 1.39 - 1.51 (m, 2 H)

**1-[(3R)-3-[(1-{1H-imidazo[4,5-b]pyridin-2-yl}piperidin-4-yl)oxy]pyrrolidin-1-yl]-2-{2-methyl-7H-pyrrolo[2,3-Ad]pyrimidin-5-yl}ethan-1-one (BI-5232A)**

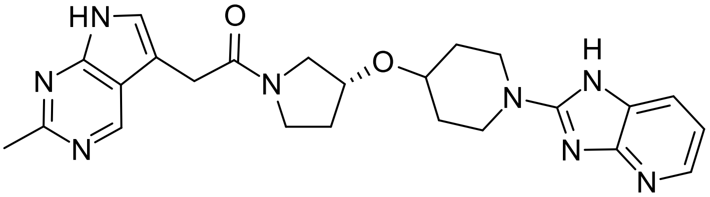

Synthesis is analogous to that described for compound **13A** using **13D** (100 mg, 0.52 mmol) and **BI-5232B** (160 mg, 0.56 mmol) as starting materials to obtain compound **BI-5232A** (130 mg, 54% yield).
LC-MS (*method 10*): $^{t}R$ = 0.61 min, $[M+H]^+$ 461

**1-{1H-imidazo[4,5-b]pyridin-2-yl}-4-[(3R)-pyrrolidin-3-yloxy]piperidine (BI-5232)**

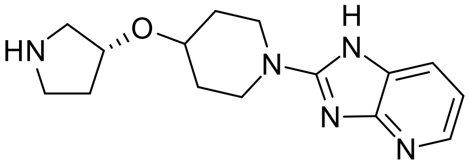

Step 1:
A mixture of 2-chloro-1H-imidazo[4,5-b]pyridine (250 mg, 1.32 mmol), **13F** (280 mg, 1.04 mmol) in ACN (3 mL) and TEA (0.44 mL, 3.16 mmol) was heated 30min at 160 °C in the microwave.The mixture was diluted with water and extracted with DCM. The organic layer was separated via phase separation cartidge and evaporated to obtain crude intermediate.
Step 2:
Crude intermediate was dissolved in DCM:TFA=3:1 (10 mL) and the mixture was stirred 1h at rt. The mixture was evaporated and the residue was purified by rp chromatography (XBridge, ACN/water/NH3) to obtain compound **BI-5232B** (161 mg, 54% yield)
LC-MS (*method 10*): $^{t}R$ = 0.60 min, $[M+H]^+$ 288

#### 5. HPLC traces of key compounds

Compound 1 (method 10):

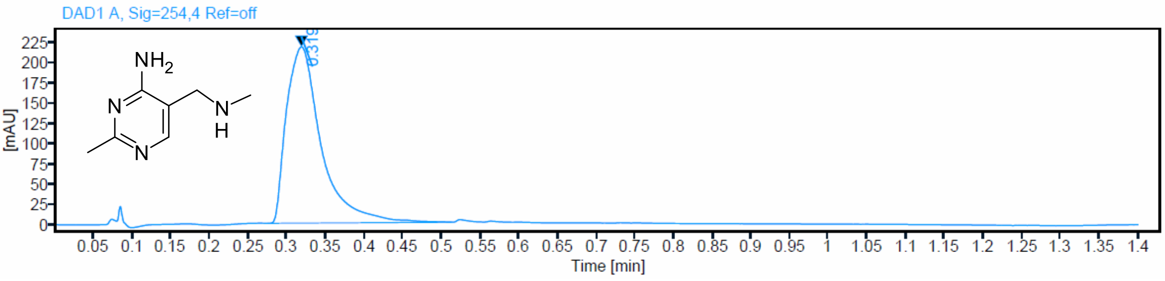


Compound 2 (method 10):

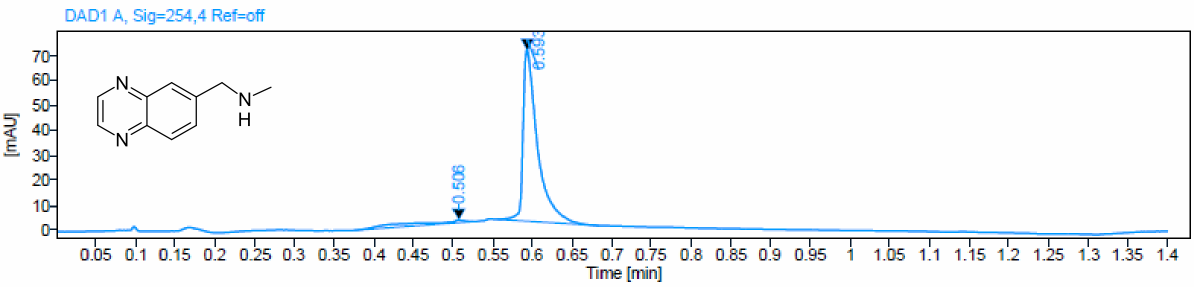


Compound 3 (method 10):

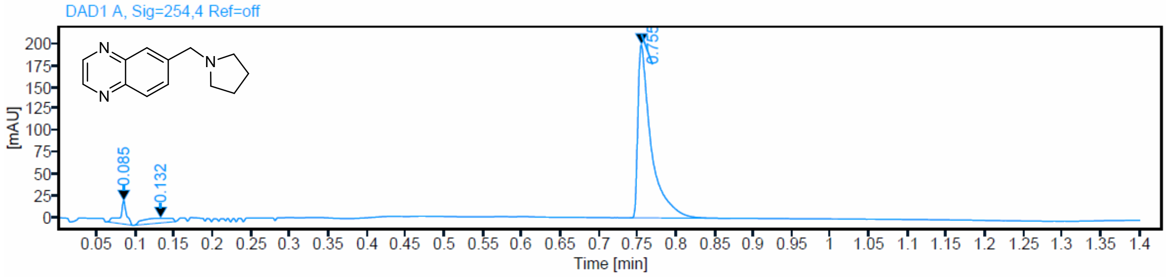


Compound 4 (method 10):

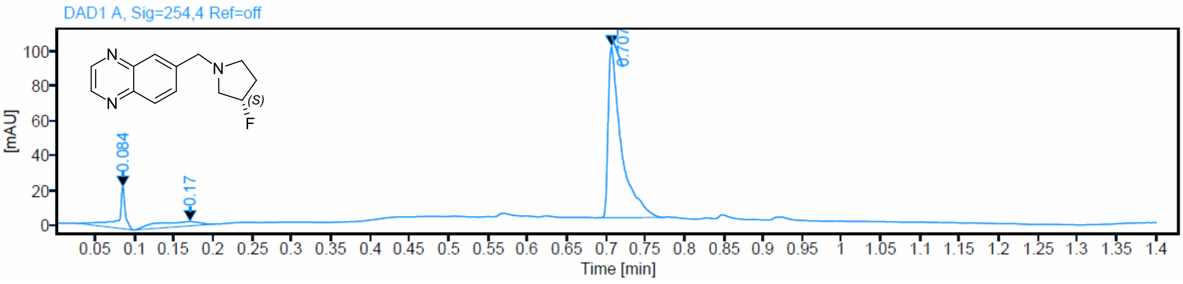


Compound 5 (method 10):

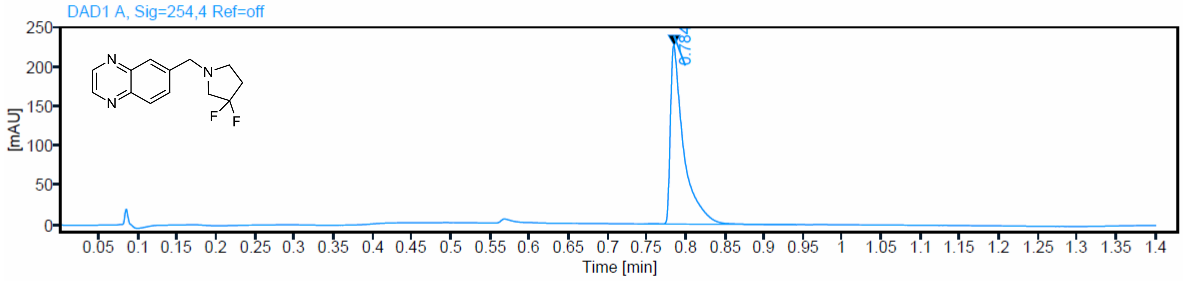


Compound 7 (method 10):

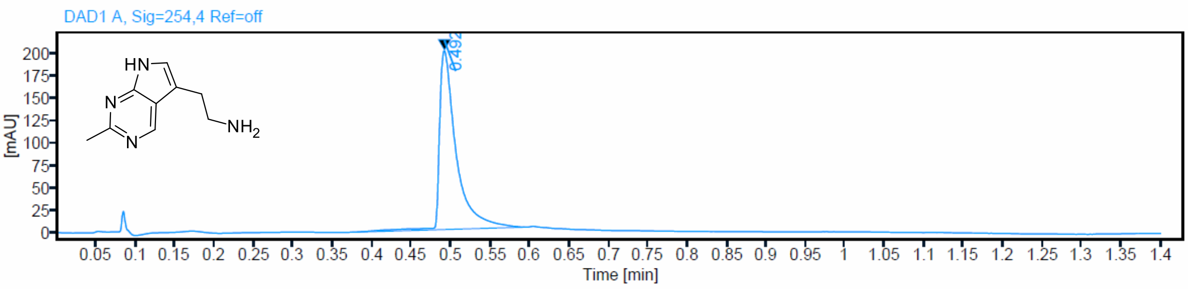


Compound 8 (method 10):

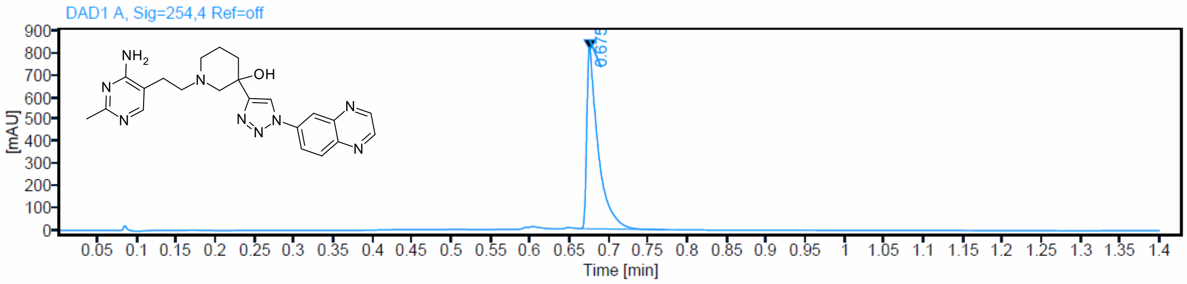


Compound 9 (method 10):

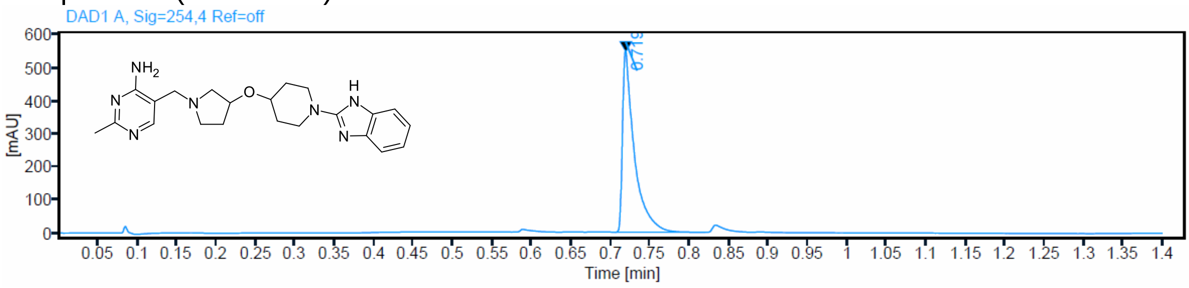

Compound 10 (method 13):

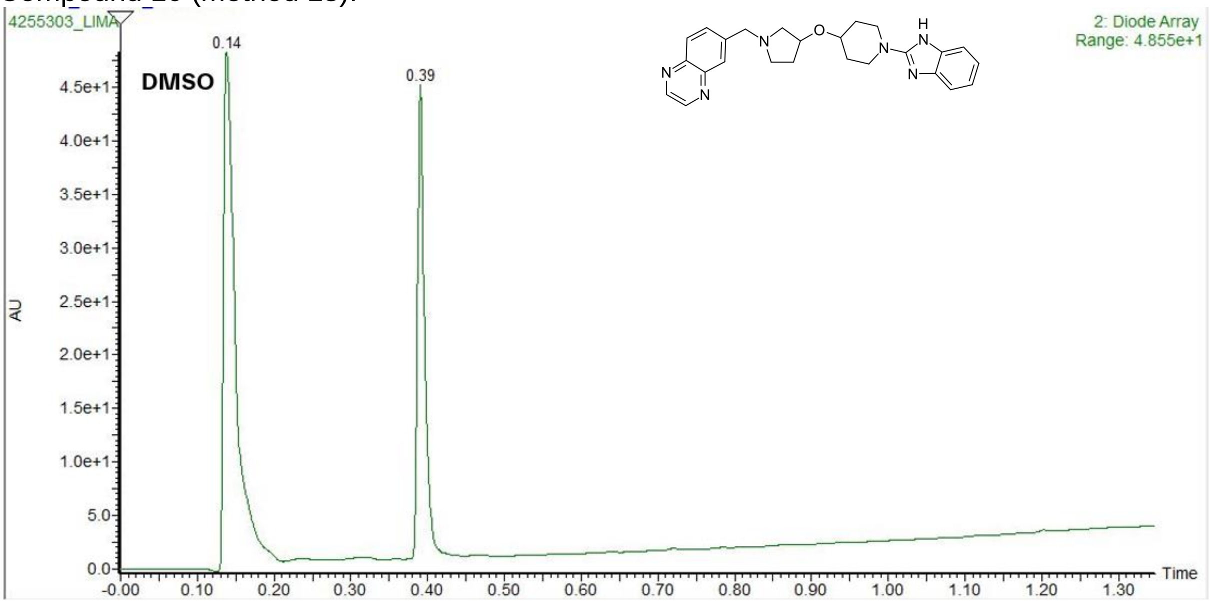


Compound 11 (method 13):

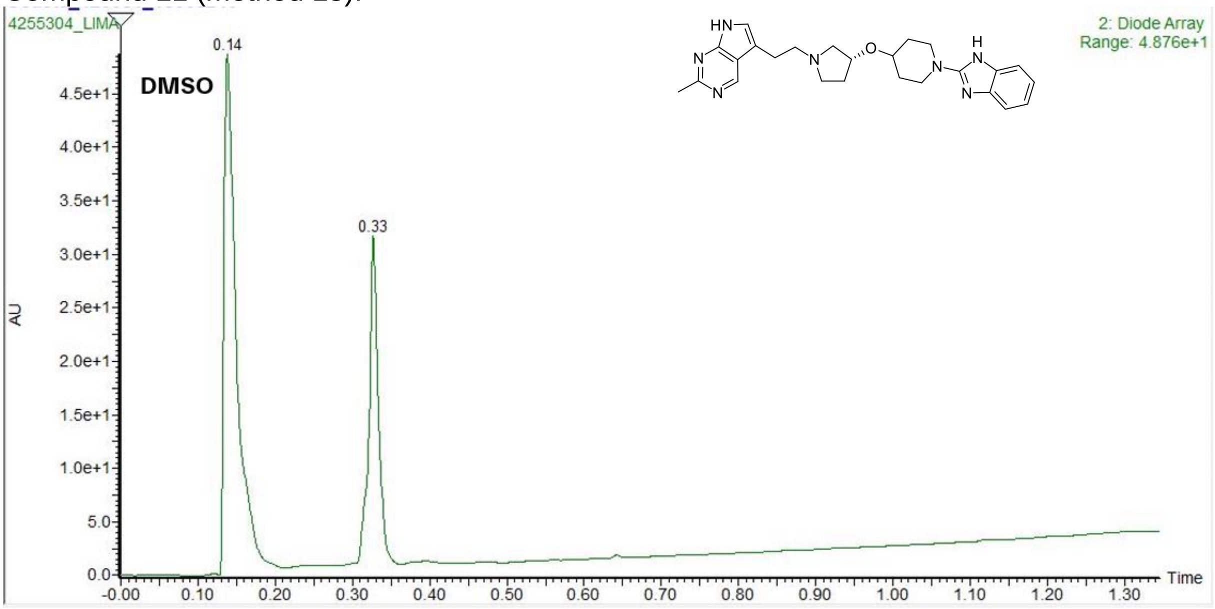


Compound 12 (method 10):

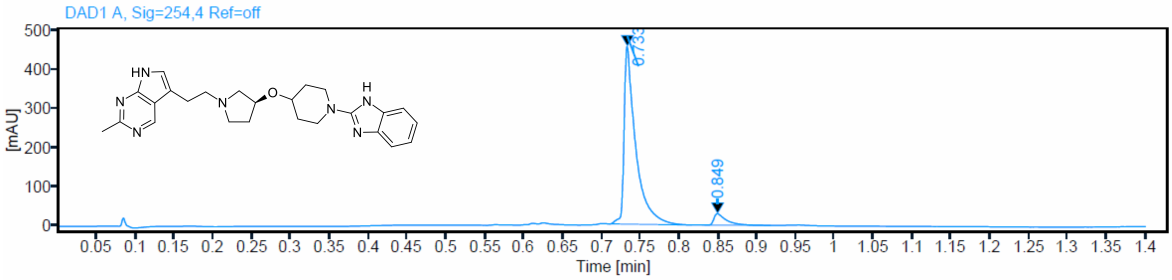

Compound 13 (method 10):

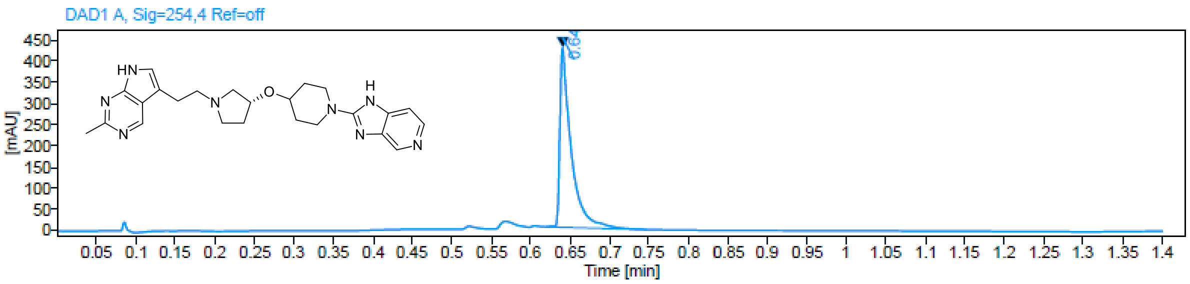


BI-5232:

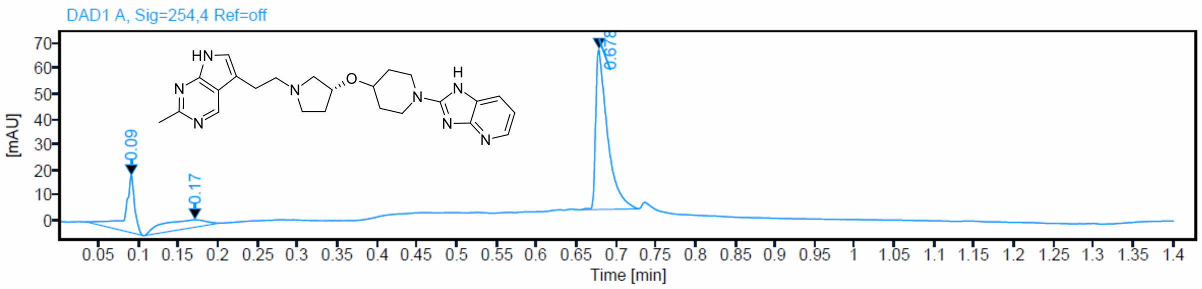


#### 6. [1]H NMR of key compounds

Compound 1

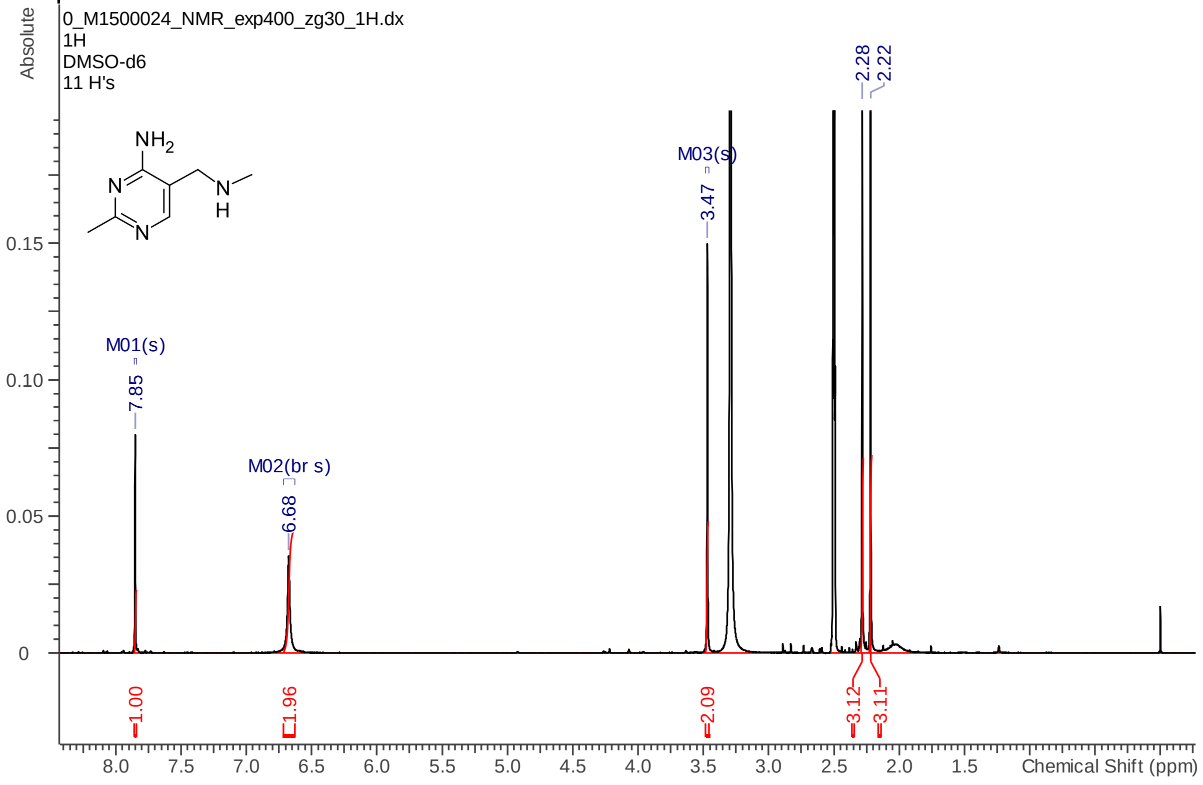


Compound 2:

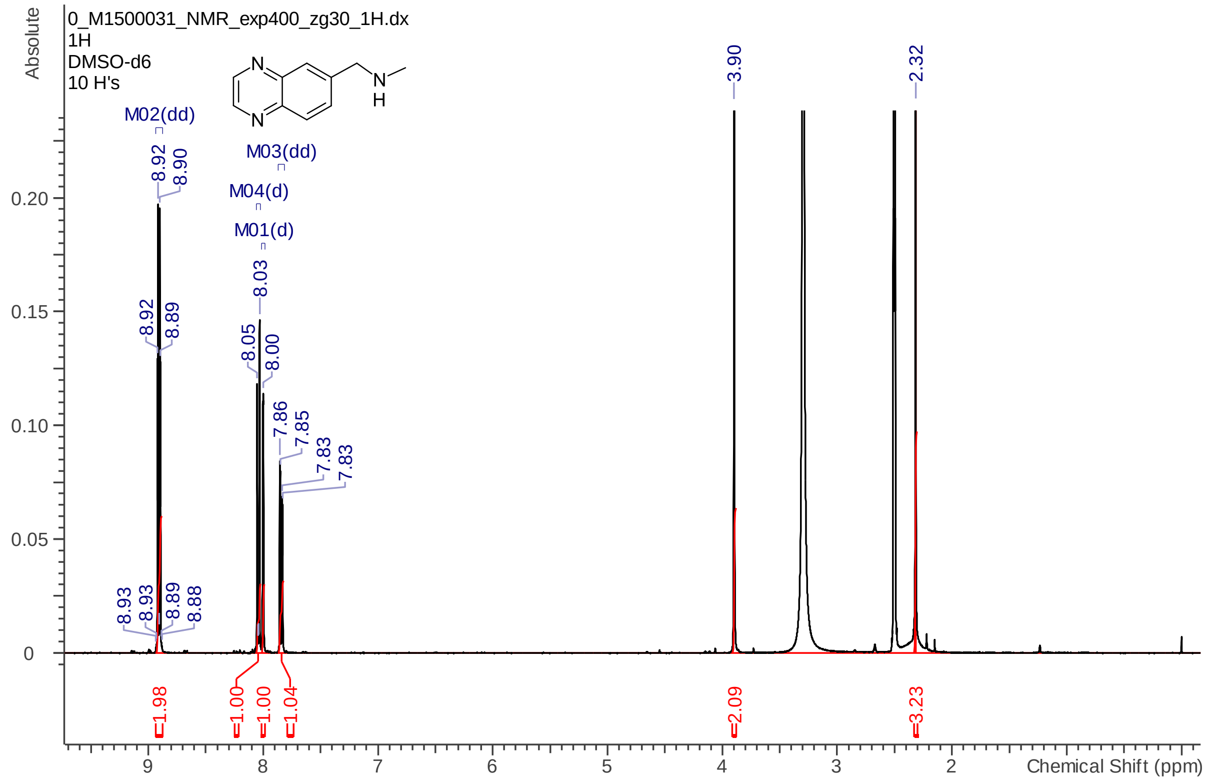
0_M1500031_NMR_exp400_zg30_1H.dx
1H
DMSO-d6
10 H's
Absolute
M02(dd)
M03(dd)
M04(d)
M01(d)
3.90
2.32
1.98
1.00
1.00
1.04
2.09
3.23
Chemical Shift (ppm)

Compound 3:

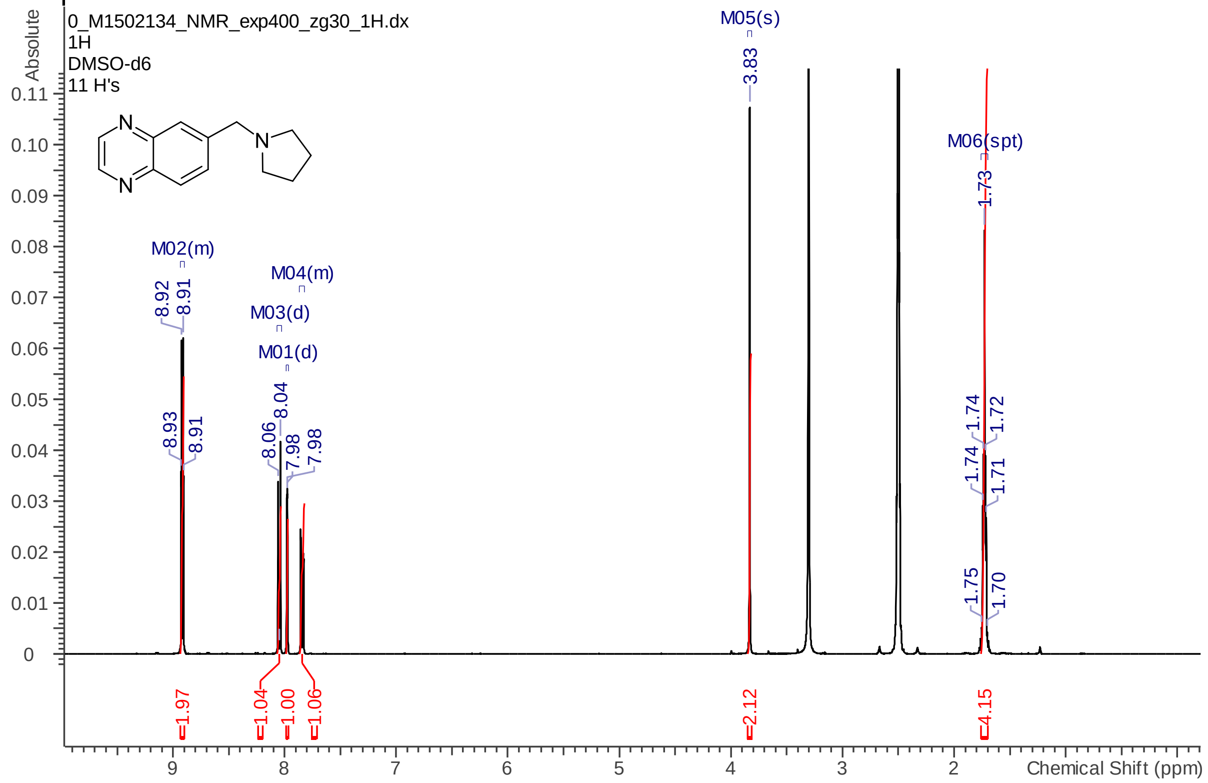
0_M1502134_NMR_exp400_zg30_1H.dx
1H
DMSO-d6
11 H's
Absolute
M05(s)
3.83
M06(spt)
1.73
M02(m)
M04(m)
M03(d)
M01(d)
1.97
1.04
1.00
1.06
2.12
4.15
Chemical Shift (ppm)

Compound 4:

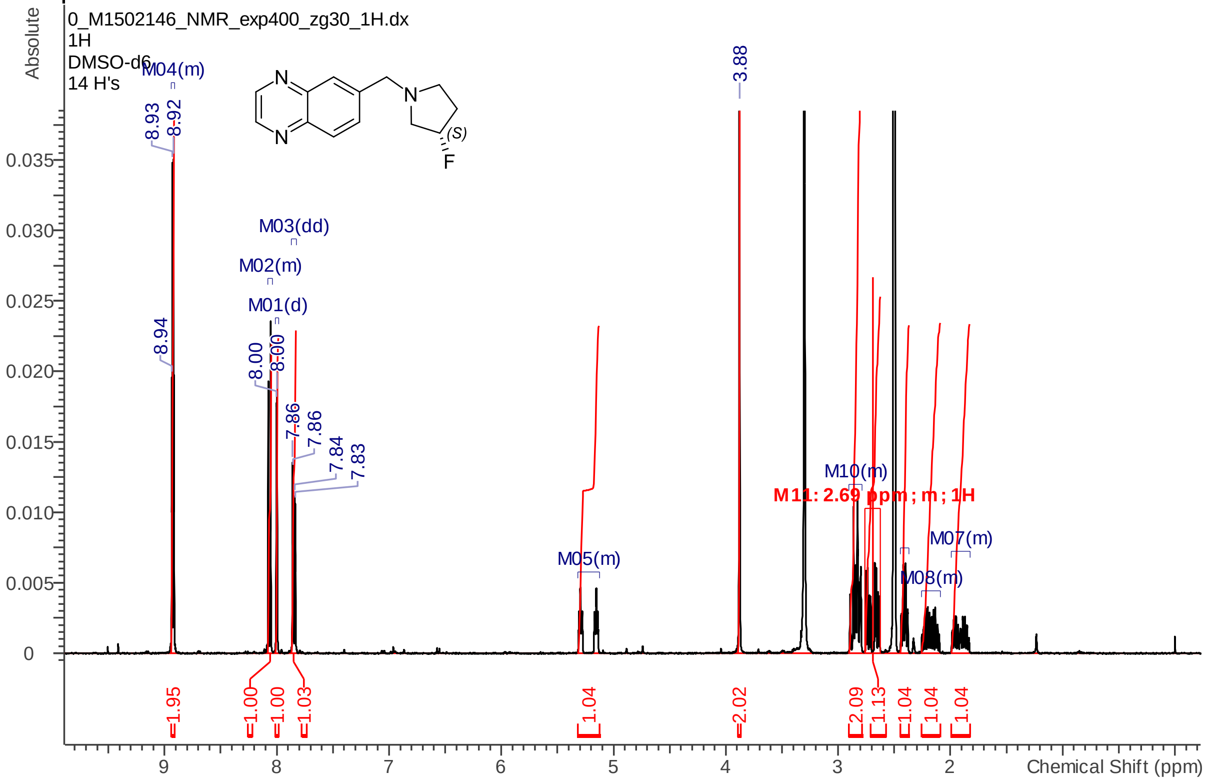


Compound 5:

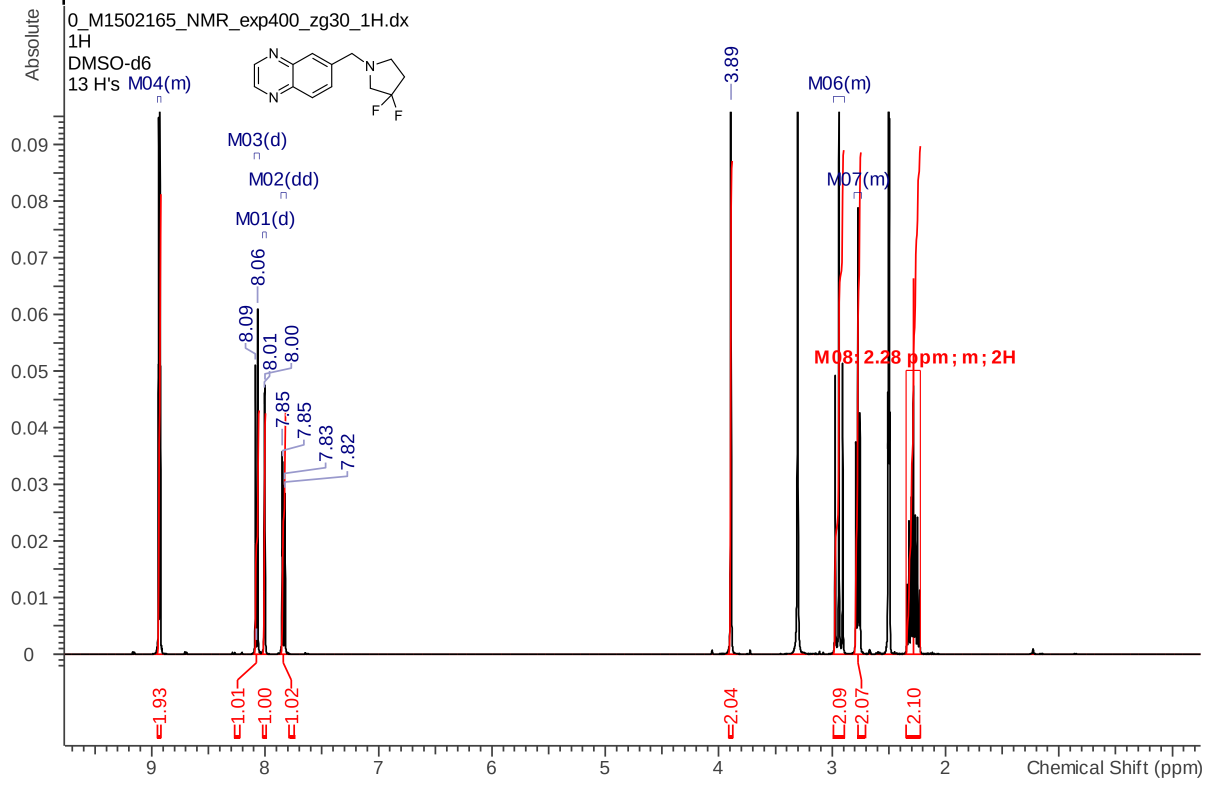

Compound 7:

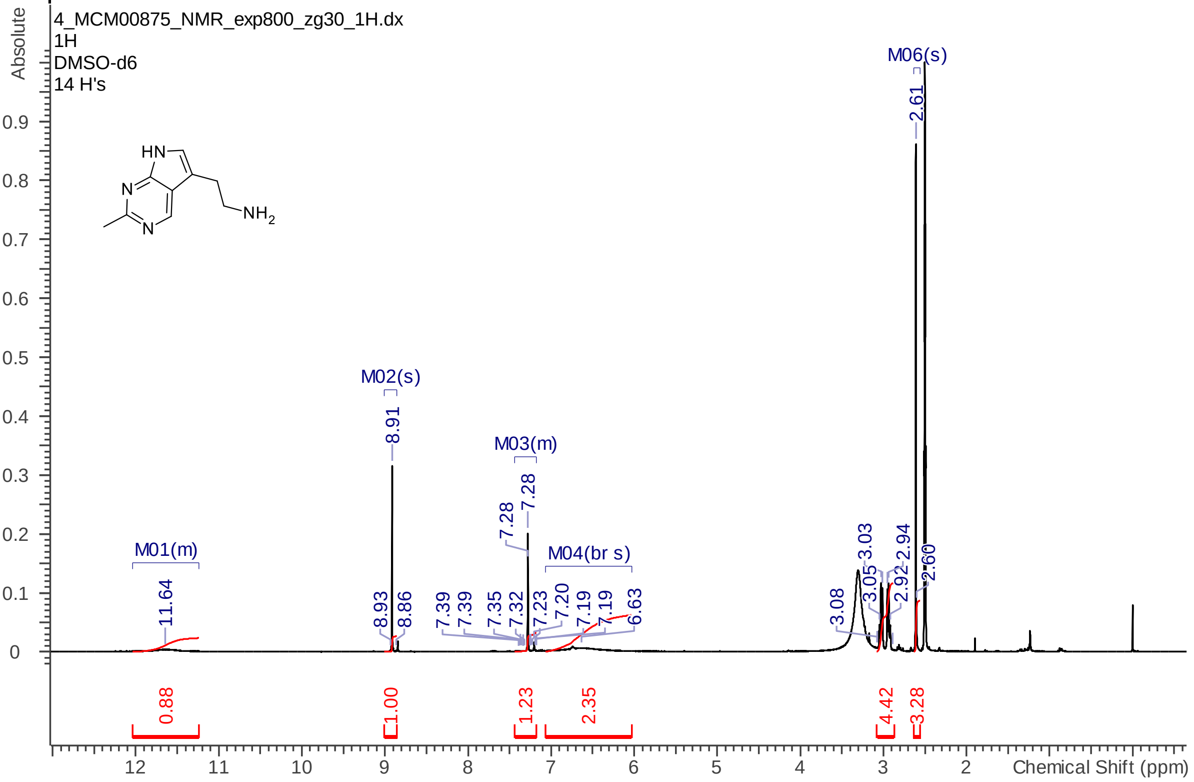


Compound 8:

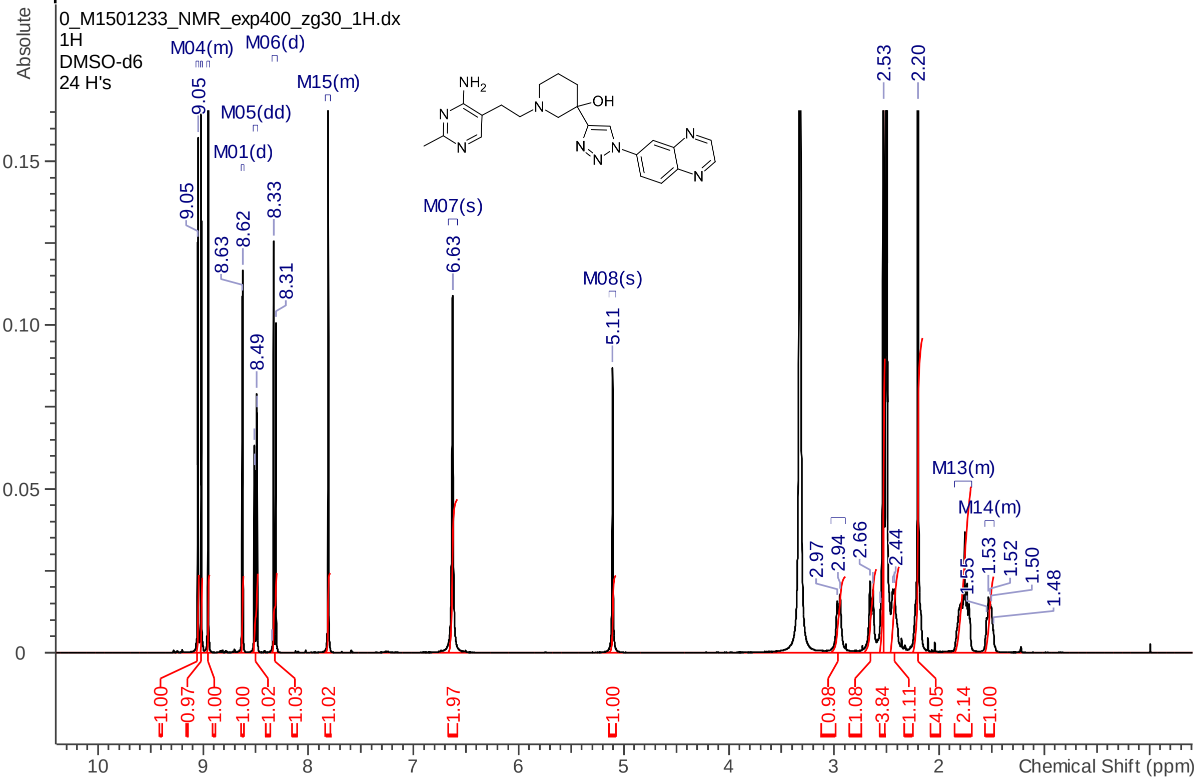


Compound 9:

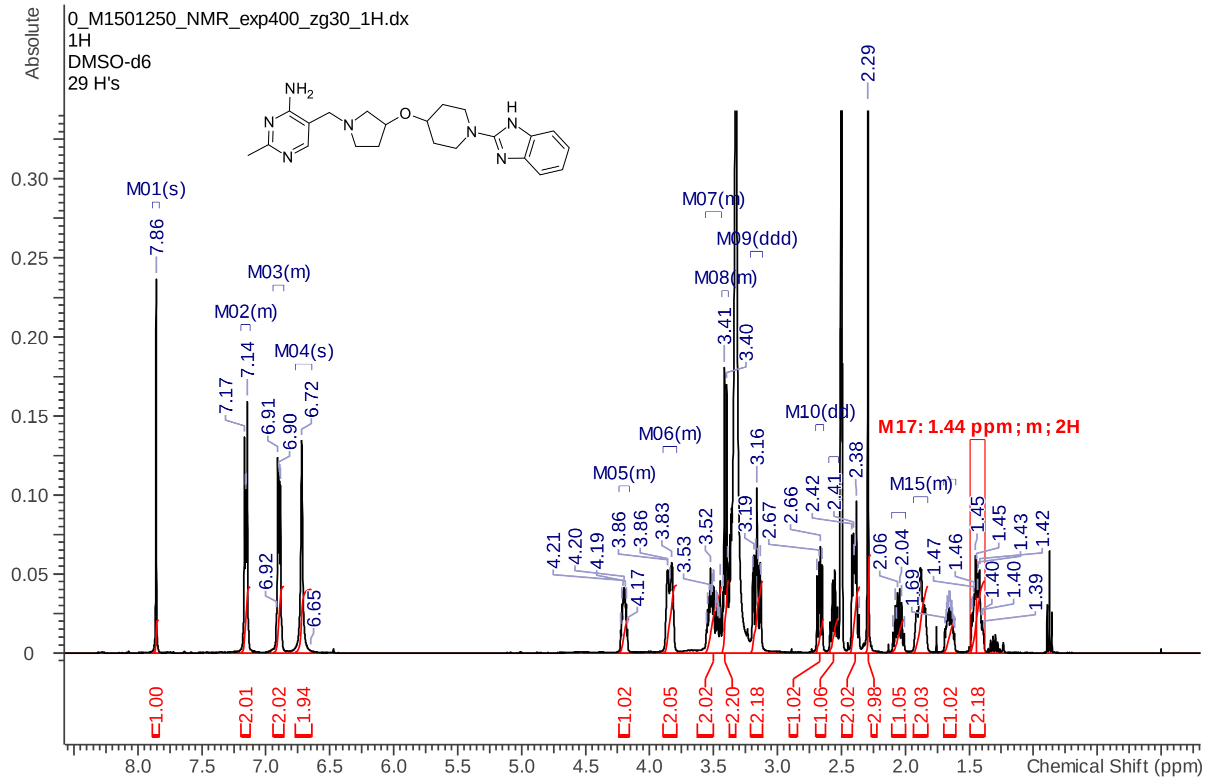
0_M1501250_NMR_exp400_zg30_1H.dx
1H
DMSO-d6
29 H's
M17: 1.44 ppm ; m ; 2H
Chemical Shift (ppm)
Absolute

Compound 10:

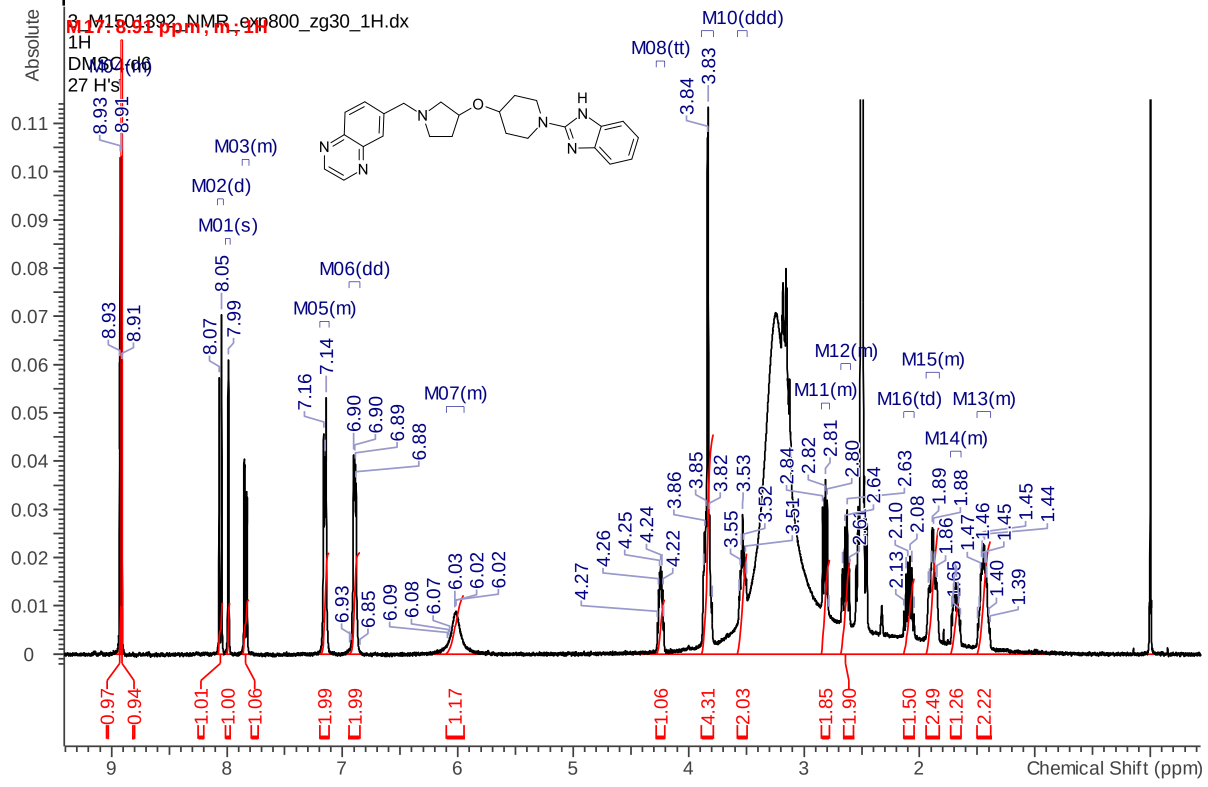
3_M1501392_NMR_exp800_zg30_1H.dx
1H
DMSO-d6
27 H's
M17: 8.91 ppm ; m ; 1H
Chemical Shift (ppm)
Absolute

Compound 11:

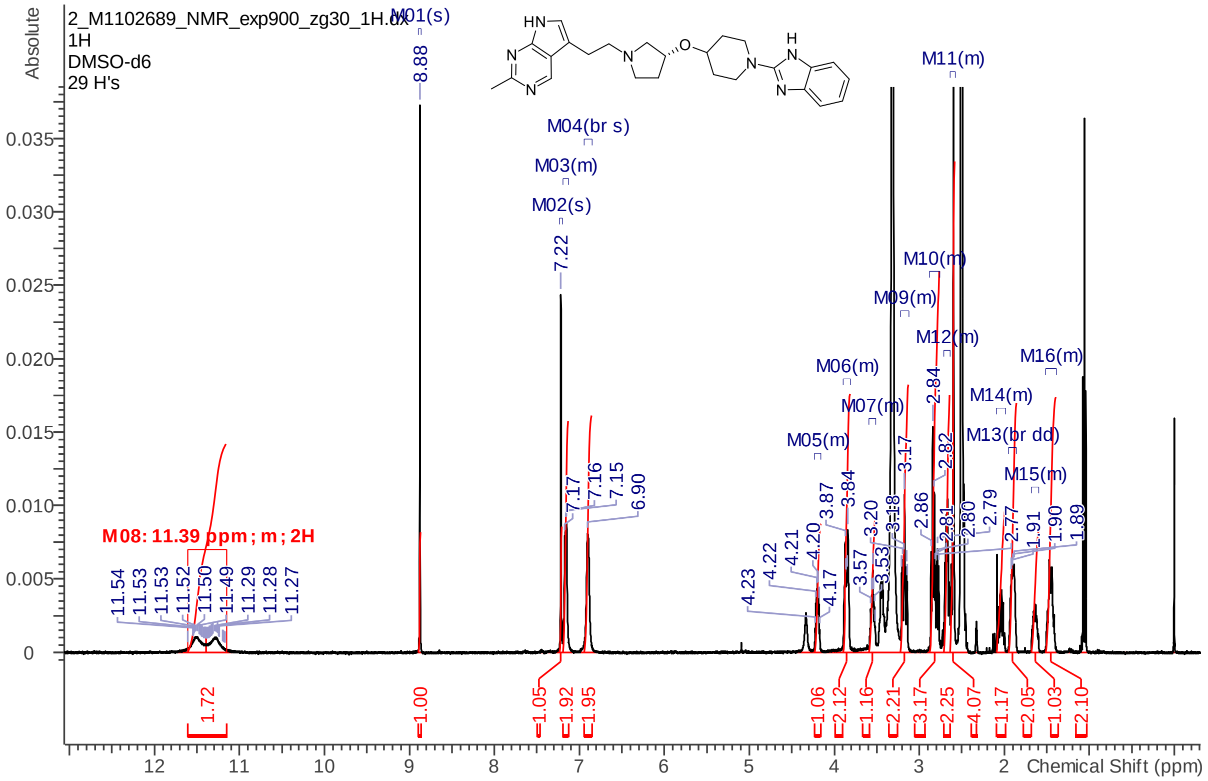


Compound 12:

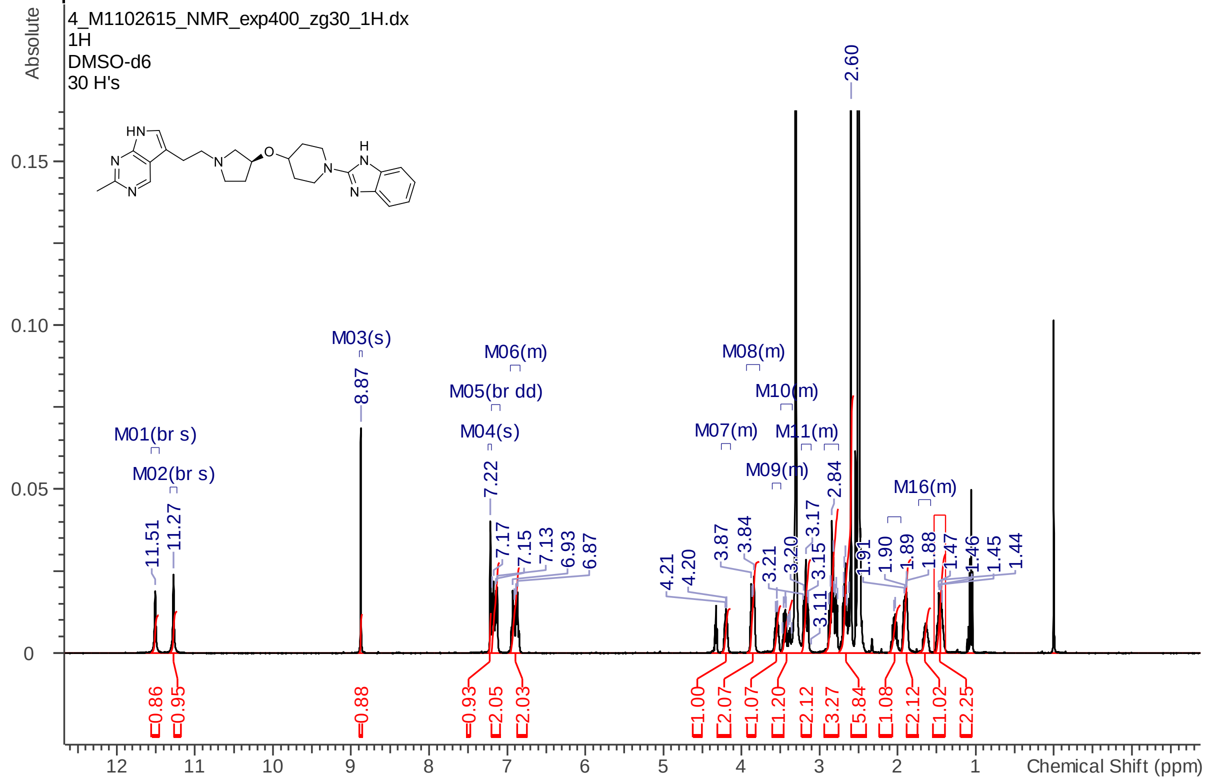

Compound 13:

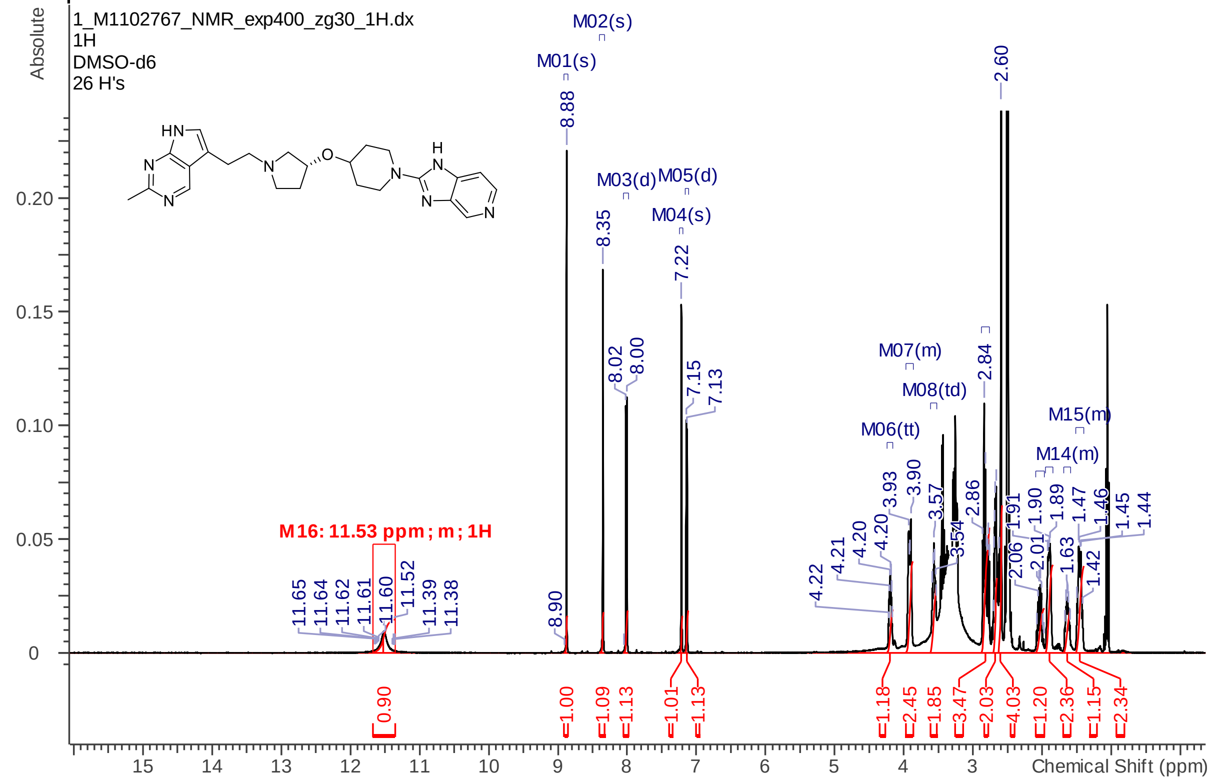


BI—5232:

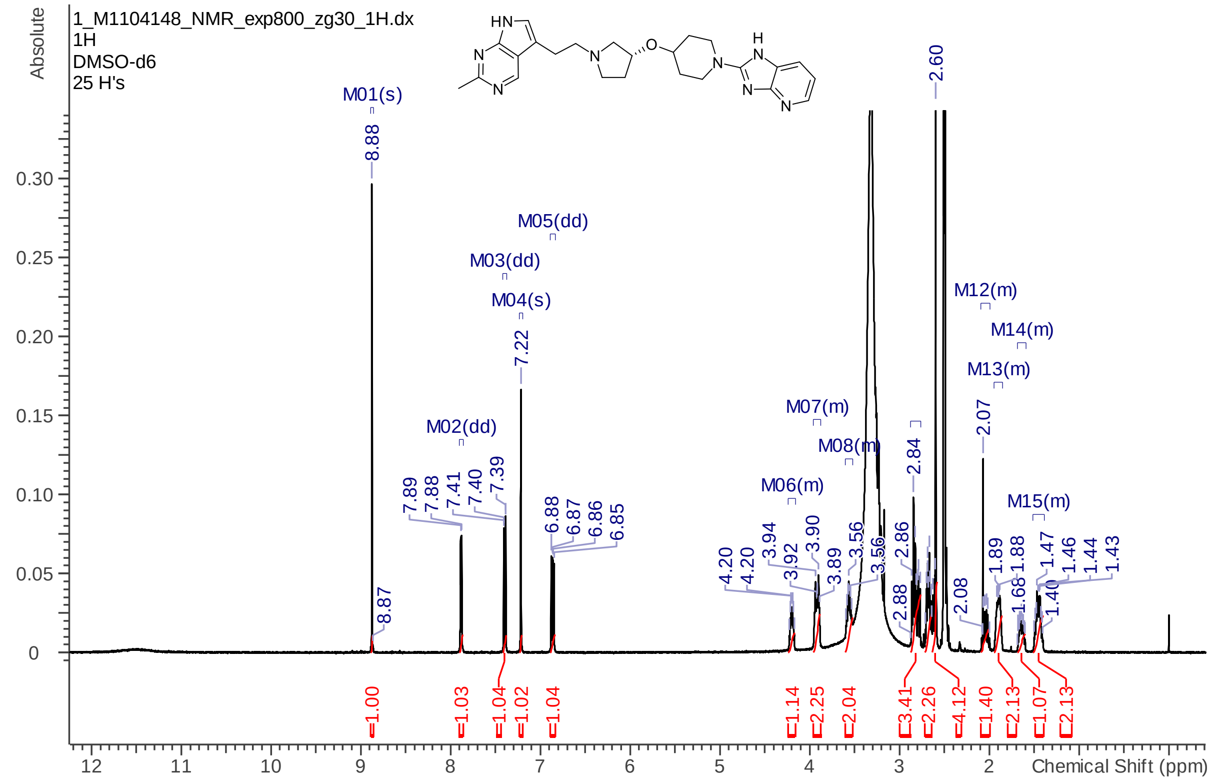

#### 7. Chiral chromatography for compounds 11 and 12

Racemic mixture compounds 11 & 12:

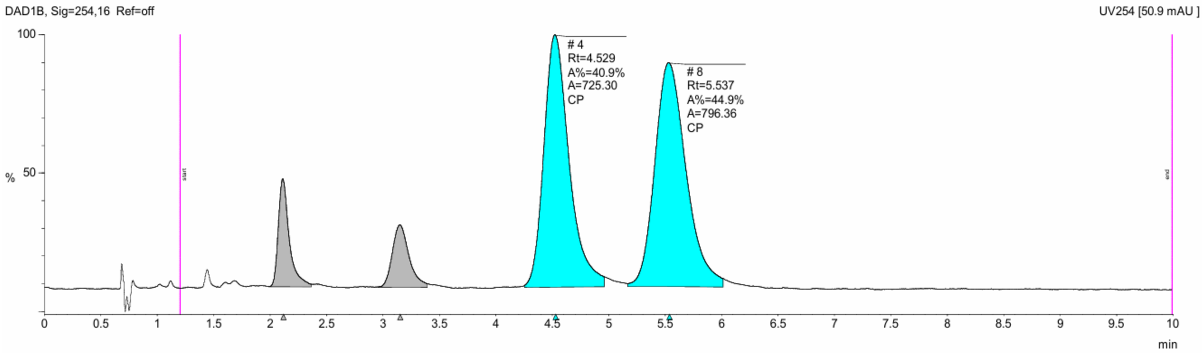


Compound 11:

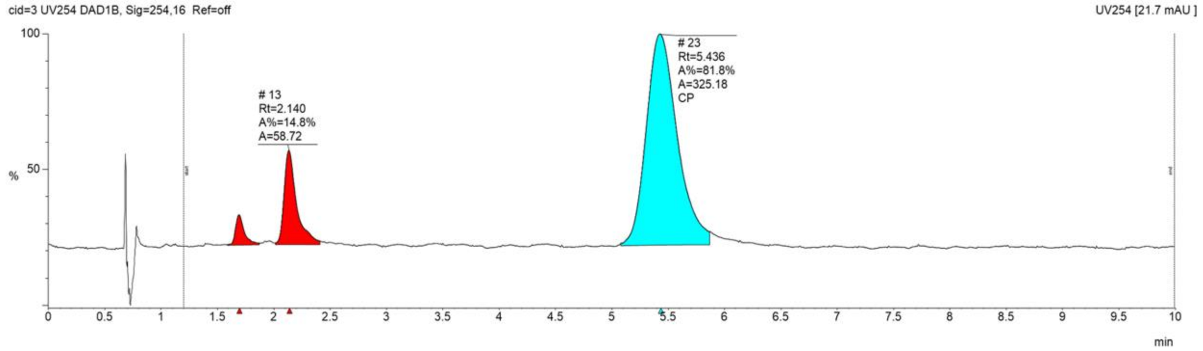


Compound 12:

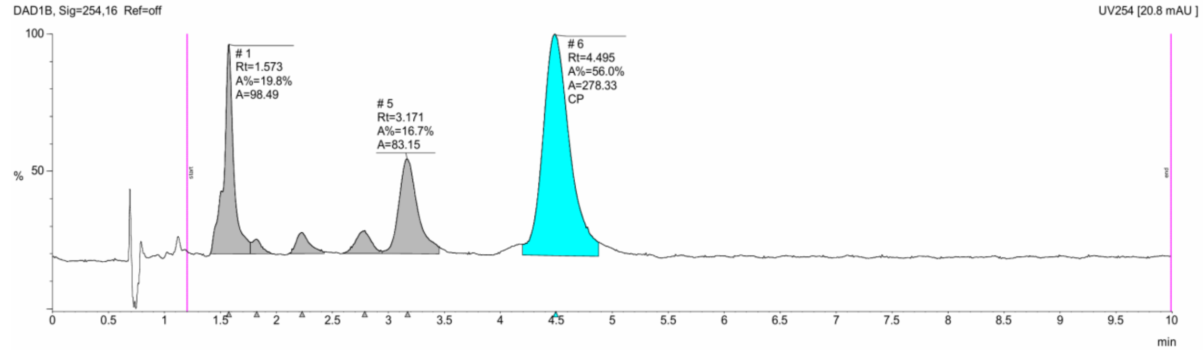
